# Chromosome segment scanning for gain- or loss-of-function screening (CHASING) and its application in metabolic engineering

**DOI:** 10.1101/2024.01.31.578163

**Authors:** Yan Xia, Lichao Sun, Zeyu Liang, ZhongRao Han, Jing Li, Yingjie Guo, Pengyu Dong, Yi-Xin Huo, Shuyuan Guo

## Abstract

Constructing a library of thousands of single-gene knockout or interference strains is a powerful tool to understand the relation between genotype and phenotype, but it is labor and cost intensive. Powered by the computer-aided gene annotation and functional grouping of non-essential genes, we showed that targeting a single gene directly to a specific observed phenotype could be quickly achieved for a specific microorganism via constructing a library of strains containing single chromosome-segment-deletion per strain. As a proof-of-concept, a genome-scale library consisting of 70 chromosome-segment-deletion strains for *B. subtilis* was constructed by CRISPR-based methods and strains with six loss-of- and gain-of-function phenotypes were screened out. To facilitate the rapid genotyping, we developed a web tool to visualize the potential targets of each chromosome segment associated with a particular function, successfully identifying the genes for valuable representative phenotypes. To apply the library to metabolic engineering, the hosts with improved production capacity of acetoin and lycopene were screened in the presence of pathway genes. This work demonstrated the significance of our strategy of **<u>ch</u>**romosome segment scanning for g**<u>a</u>**in- or lo**<u>s</u>**s-of-function screen**<u>ing</u> (CHASING)** on functional genomics investigation, robust chassis engineering, and chemical overproduction.

TOC

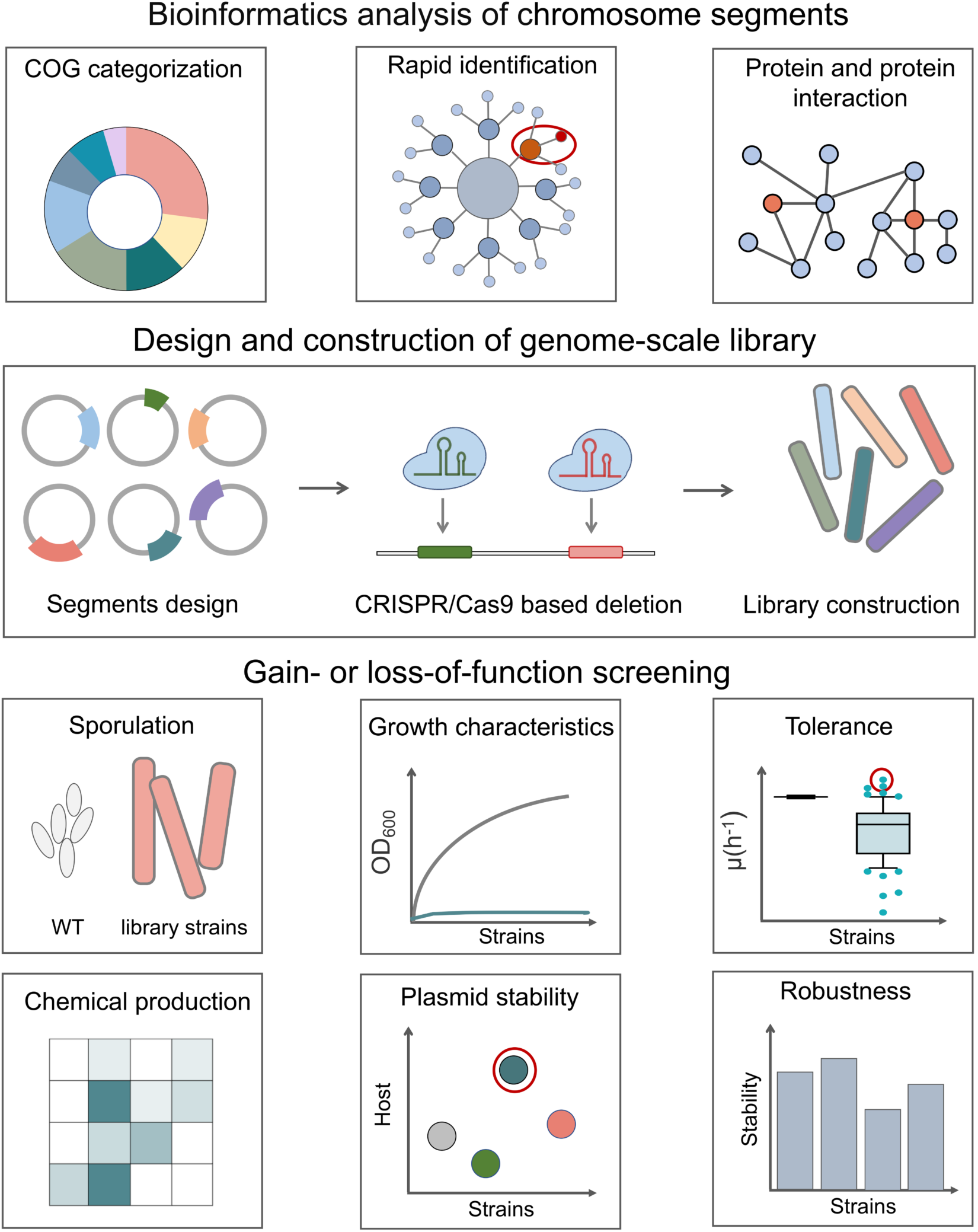

## 1. Introduction

Native bacteria maintain homeostasis through regulation of the expression of the functional proteins at transcriptional and translational levels. The deactivation of a gene could lead to the loss or destroy of a function while the deactivation of a regulator could lead to function restore via de-repression or de-regulation of a previously depressed enzyme or pathway. Dissecting the genotype-phenotype map, which implies the relationship between genotype variations and phenotype changes(de VienneCapy, 2022), provides valuable insights into the engineering of robust chassis and is essential for industrial chemical production.

Developing high-throughput approaches is important for understanding the sequence-function as well as gene-network architecture(BrochadoTypas, 2013). Constructing randomly-generated mutant libraries has been widely employed using transposon-induced mutation strategies(van Opijnen, et al., 2009). However, the transposon-mediated mutation resulted in insertion bias toward genes with long coding regions(Kazi, et al., 2020; T. Wang, et al., 2018). With the advance of CRISPR technique, pooled CRISPR interference (CRISPRi) technique has been applied for screening by designing the genome-scale sgRNA libraries in microorganisms such as *Escherichia coli*(Rousset, et al., 2018; T. Wang, Guan, et al., 2018), *Synechocystis* sp PCC 6803(Yao, et al., 2020) and *Eubacterium limosum*(Shin, et al., 2023), despite the limitations such as the variation of the binding affinity of sgRNAs and the fluctuated repression level during sampling(de Bakker, et al., 2022; Doench, et al., 2016; T. Wang, Guan, et al., 2018).

An alternative high-throughput approach for mapping phenotypes is the genome-arrayed single-gene mutant libraries containing thousands of strains. By knocking out the non-essential genes individually, collections of single-gene-deletion strains have been reported for elaborating genome architecture and stability in *E. coli*(Baba, et al., 2006; Yamamoto, et al., 2009), *Bacillus subtilis*(Koo, et al., 2017), and *Saccharomyces cerevisiae*(Puddu, et al., 2019). Single-gene-knockdown strains have also been obtained using CRISPRi technique, revealing a systematic essential-gene network of *B. subtilis*(Peters, et al., 2016). Although exploring these libraries avoids the unknown targeting effect, the library construction requires enormous labor work and the operating processes for screening and storage are challenging(Costanzo, et al., 2019; Henser-Brownhill, et al., 2017).

With decades of efforts, various databases have been constructed to aid the functional genomic analysis of the organism of interested(Galperin, et al., 2021). As an example, the database of Clusters of Orthologous Genes (COGs) includes 26 COG categories such as amino acid metabolism and transport, transcription, translation, cell wall/membrane/envelop biogenesis, and signal transduction. The setting of category number to 26 is a trade-off between applicability and precision since the 26 categories could not specify all the cellular functions(Galperin, et al., 2019).

To provide an alternative simple strategy for large-scale genotype-phenotype mapping, we sought to establish an arrayed chromosome-segment-deletion (CSD) library and grouped the genes within segments to a specific cluster of function. By taking advantage of the well-established database of COG, our analysis showed that the CSD library could cover all non-essential genes, having 46.1%, 64.6%, and 45.8% of probability to link a single gene directly to an observed phenotype in *B. subtilis* 168, *Bacillus thuringiensis* BMB171, and *E. coli* MG1655, respectively. Using CRISPR-based large-fragment deletion methods, we yielded a genome-scale library consisting of 70 CSD strains for *B. subtilis* in this work. As a proof-of-concept, we applied this arrayed genome-wide library to rapidly identify the genetic modifications underlying the loss-of- and gain-of-function phenotypes. We also evaluated the applicability of the CSD library in metabolic engineering applications. Specifically, we successfully screened out two lycopene-overproducers reaching 54.1% and 18.2% increase in yields, and five sporulation-defective mutants reaching as high as 20.8% increase in acetoin titers. Our work highlighted the significance of the CHASING strategy in diverse applications.

## 2. Method

### 2.1 Plasmid construction for chromosome segment deletion

The relevant plasmids and primers can be found in Table S3 and S4. All primers were synthesized by Genewiz (Tianjin, China). The knockout plasmid backbone, p8999-2N20, used in our study, contains Cas9 under mannose-induced promoters and two sgRNA expression cassettes driven by the constitutive promoter *P_veg_*. To target the chromosome segment, we carefully selected two highly efficient N20 sequences located at the beginning and end of the target region using the Zhanglab website. For the rapid and simple construction of various knockout plasmids, we simultaneously introduced two sgRNAs using Gibson assembly. Next, the plasmid containing the target sgRNA serves as the template for the second round of PCR. The upstream and downstream homology arm sequences are amplified from the *B. subtilis* genome, and then fragments are generated using overlap extension PCR (OEPCR). Subsequently, the backbone and fragments are joined together through Gibson assembly, resulting in the formation of a complete knockout plasmid.

### 2.2 Construction of genome-scale chromosome-segment deletion library

In order to construct a genome-scale deletion library for *B. subtilis*, we designed the N20 sequences to target the proposed regions and constructed the plasmids using the highly efficient sgRNAs along with 2 kb repair template. The resulting plasmids were transformed into *B. subtilis* 168 by natural transformation method. 20 μL of cells was plated on the LB agar plates with 20 μg/mL kanamycin and 0.2% mannose, and incubated at 30°C for editing. After 15 h, we selected 8 colonies from each plate for deletion validation. Two pairs of validation primers were designed for each deleted segment, the first pair was designed inside the deleted segment and the second pair was designed outside the region of homology arm. Theoretically, the positive colony would yield PCR products with the correct length using the first pair and no product using the second pair. The editing efficiency was calculated as the ratio of the number of positive colonies to the selected ones. The results were further validated through Sanger sequencing. Once the single colony that containing the deleted segment was verified, plasmid elimination was carried out using the method as previously(Tian, et al., 2022). The chromosome-segment-deletion strains listed in Table S5 were preserved at −80°C.

### 2.3 Bioinformatics analysis of the genome-scale deletion library

COG annotation was accomplished using eggnog-emapper. All relevant code were displayed in Note S1. All available data of protein-protein interactions in *B. subtilis* were retrieved from the STRING database(Szklarczyk, et al., 2023). Data of protein-protein interactions with a combined score greater than 800 were considered reliable. Python was employed to generate the graph of protein-protein interaction while NetworkX and Matplotlib was used for graph construction and visualization, respectively. The distribution of undeleted genes and deleted genes was embedded using a 2D representation by the t-SNE projections.

### 2.4 Phenotype screening

The survival capability of the resulting 75 strains were examined in various conditions, including LB medium, GMII medium, GMII medium under acidic conditions (pH 5), and GMII medium with 60 g/L urea. The strains in the library were streaked onto LB agar plates and incubated at 37°C for 12 h. Then, three single colonies were picked from each plate and inoculated into the fresh culture, followed by incubation at 37°C and 220 rpm for 12 h. Subsequently, they were transferred into 20 mL of LB broth in shake flasks at a 1% of inoculation ratio and incubated at 37°C and 220 rpm for 24 h. During the incubation period, growth measurements were taken every 4 h. For growth measurement, 200 μL of the culture was transferred into a 96-well plate, and the OD_600_ was measured using a spectrophotometer. Blank LB medium was used as a control, and the OD_600_ values were obtained by subtracting the blank control values. Growth curve of each strain was plotted based on these values.

### 2.5 Transcriptomic analysis

The cell cultures were collected in logarithmic phase (OD_600_ of 1.0) by centrifugation at 5,000 rpm for 10 min under 4°C. The cell cultures were then subjected to liquid nitrogen frozen and stored at –80°C. Total RNA was extracted from the collected cells using the methods reported previously(Peng, et al., 2020). The library construction, purification, and Illumina sequencing were completed by GENEWIZ company (Tianjin, China). To ensure the quality of sequencing data, thorough pre-processing of the raw data was conducted, including the meticulous filtering of low-quality data, meticulous removal of contaminants, and precise trimming of adapter sequences. The gene expression was analyzed with the HTseq software (V 0.6.1) using the FPKM (Fragments Per Kilobase per Million reads) method, which was proposed by Mortazavi in 2008(Mortazavi, et al., 2008). The differential expression analysis was conducted using the DESeq2 (V1.6.3) package from Bioconductor. The cutoff of the differentially expressed gene was set at a log2-fold change > |1| with a q-value (FDR, padj) < 0.05.

### 2.6 Screening for plasmid stability

We constructed the plasmid pXY-sfGFP and transformed it into each strain of the genome-scale deletion library. The resulting strains were inoculated into 5 mL of LB liquid medium that containing antibiotics and cultured at 37°C and 200 rpm for 8 h. Subsequently, the seed culture was transferred to LB medium without antibiotics at a inoculation ratio of 1%. Subculturing was performed every 24 h and 200 μL samples were taken at 8 h, 32 h, and 56 h for fluorescence measurement at OD_474_. The plasmid stability was calculated using the formula: Plasmid stability = (Fluorescence value in LB without antibiotics / Fluorescence value in LB with antibiotics) ×100%.

### 2.7 Acetoin fermentation and product detection

The acetoin fermentation plasmid was obtained from our lab stock. Strains of interested were transformed with these plasmids. Single colonies were selected and inoculated into 5 mL of LB liquid medium for incubation at 30°C and 200 rpm for 8 h. Subsequently, the seed culture was transferred to 20 mL fermentation medium with or without antibiotic as required. The fermentation process was conducted at 30°C and 200 rpm, and the sample was taken every 24 h. To detect the acetoin product, gas chromatography was performed using an instrument (PANNA GCA91) equipped with an FID detector. All experiments were performed in triplicates, and the data was presented as an average value.

### 2.8 Screening for lycopene production

The lycopene biosynthetic genes including *crtE*、*crtB* and *crtI* from *Pantoea ananas* were expressed in the plasmid, yielding the strain pHT01-crtEBI. The seed culture was with transferred at an inoculum of 5% to 20 mL of LB liquid medium with the corresponding antibiotics. The fermentation was conducted at 50 mL shake flasks and 1 mL fermentation broth was taken every 24 h. To determine the cell dry weight and lycopene production, the sample was subjected to centrifugation to separate the supernatant, and the pellets were dried in a 65°C oven. The biomass was weighed to calculate the lycopene production per unit dry weight. The lycopene extraction was conducted using the acetone extraction method. The bacterial culture was centrifuged at 12,000 rpm for 2 min and the pellet was resuspended twice using 1 mL of ddH_2_O. Then, 200 μL acetone was added to the pellet, and the cells were lysed with beads using a cell disruptor. This process was performed for 20 min with 25 s of operation followed by 5 s of interval. The samples were transferred to a water bath of 55°C and lycopene was extracted for 1 h. Then, the resulting supernatant was collected and the lycopene titer was calculated by measuring the absorbance at 474 nm in the spectrophotometer.

## 3. Results

### 3.1 Concept of the CHASING strategy

Functional genomic analysis and function assignments for proteins are important for tracing the corresponding genotype underlying an interested phenotype. Theoretically, every gene along the chromosome could be assigned to a specific function, yielding personalized clusters of function (PCOF). For a specific chromosome segment, the genes in the segment are likely to belong to diverse PCOFs. If a segment does not contain an essential gene, it could be deleted and the loss of functional non-essential genes might lead to certain phenotypes. If any phenotype is desirable to the metabolic engineering purpose, the change of phenotype is likely due to the loss of a gene or genes that fall into the PCOF category that related to the phenotype. Thus, we could rapidly identify the gene or genes that functionally link to a specific phenotype.

### 3.2 Genome-wide analysis for large-scale genotype-phenotype mapping

By exploring the PCOF distribution profile in the chromosome using the genome-wide COG distribution as a representative, we did *in silico* chromosome segment scanning in model microorganisms. Taking *B. subtilis* 168 as an example, there are 257 essential genes and 4018 non-essential genes in the genome (Fig. 1a). The 4018 non-essential genes that belongs to 22 COGs are distributed dispersedly among these essential genes. If each essential gene was set as a dividing point, the whole genome could be divided into 257 segments. Taking into account that in 109 cases no non-essential gene exists between two essential genes, the whole genome was finally divided into 148 large segments by essential genes. The 148 segments were analyzed (Fig. 1b) and all the genes in each segment were allocated into a specific COG. For example, segment 82 contains COGs from eleven categories, and segment 31 contains COGs from fifteen categories (Fig. 1c).

**Fig. 1.**
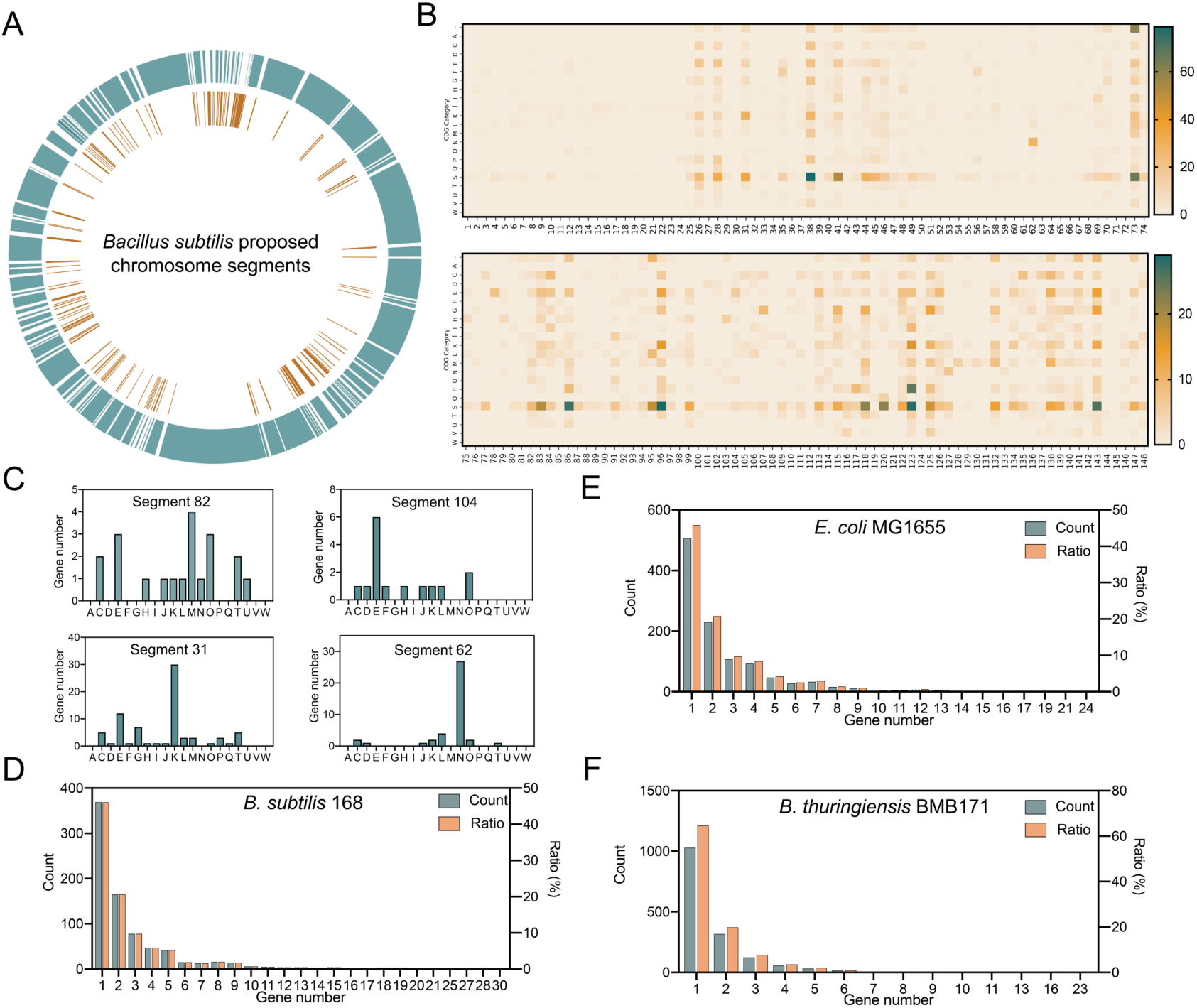
Bioinformatic analysis of chromosome segments in three model strains. (a) The proposed chromosome segments in *B. subtilis* 168. The inner orange lines indicate the position of essential genes. The 148 proposed chromosome segments were indicated in blue. (b) COGs distribution in the proposed chromosome segments in *B. subtilis* 168. (c) Gene number analysis of each COG category in segments 82, 104, 31, and 62. (d) Accumulated COG categories with different gene numbers in *B. subtilis* 168. The data was calculated based on the COGs in all the proposed chromosome segments. (e) Accumulated COG categories with different gene numbers in *E. coli* MG1655. (f) Accumulated COG categories with different gene numbers in *B. thuringiensis* BMB171.

We then analyzed the gene numbers of every category in every chromosome segment. As shown in Fig. 1d, the probability of having one or two genes belongs to a random COG category in a random segment is 46.1% or 20.6%, respectively. The probability of having more than two genes belongs to a random COG category in a random segment is less than 34%. For example, eight COG categories in segment 82 or segment 104 contains only one or two genes (Fig. 1c). In contrast, 30 genes in segment 31 belong to COG category K and 27 genes in segment 62 belong to COG category N, indicating that gene clusters exist in some large segments. If segment 31 or 62 was deleted, then the probability of observing the K- or N-related phenotype would be high. Similar results have been obtained for *E. coli* (Fig. 1e) and *B. thuringiensis* (Fig. 1f) by analyzing the genome-wide COG distribution profile. Specifically, the probability of having one gene belongs to a random COG category in a random segment is 45.8% and 64.6% in *E. coli* and *B. thuringiensis*, respectively.

Taken together, a library of 148 CSD strains could cover all non-essential genes in *B. subtilis* 168, having 46.1% of probability to link a single target gene directly to an observed phenotype. Besides this simplest scenario, this library has 20.6% of probability to zoom the candidates into two genes for an observed phenotype, thus is easy to identify the target gene(s) via another round of knockout. A library that fits into a 96-well plate could rapidly identify the target gene for a desired phenotype, showing great potential in large-scale genotype-phenotype mapping. Considering the positive control, negative control and necessary blank, a library of 70 to 80 strains is a reasonable size for achieving this purpose.

### 3.3 Constructing a genome-scale deletion library in *B. subtilis*

Taking advantage of the ten *Bacillus* large-fragment knockout strains in our lab(Tian, Xing, et al., 2022), we designed a 75-CSD library for *B. subtilis* to provide a proof-of-concept demonstration for large-scale genotype-phenotype mapping. Herein, 65 chromosome-segment deletions were performed individually in this study (Fig. 2a). For every specific segment deletion, two sgRNAs targeting both ends of the segment and a repair template were designed. For example, a 50.6 kb segment spanning from *yefB* to *yetK* was deleted in the Δ8 strain using sgRNAs targeting the upstream of *yefB* and the downstream of *yetK* along with a 2000-bp repair template, reaching a positivity rate of 100% (Fig. 2b). Similarly, a smaller segment of 17.3 kb spanning from *cotT* to *yilB* genes was deleted in the Δ17 strain using two sgRNAs targeting the upstream of *cotT* and the downstream of *yilB*, yielding a positivity rate of 75%. Among all these segments, 48 segments were deleted with positivity rates higher than 60%, including 31 segments that were deleted with positivity rates of 100% (Fig. 2c). The length of the designed segments ranges from 3.7 to 201 kb (Fig. 2d). These results indicated the high efficiency of our editing method in large-fragment deletion. To be noted, there is no direct correlation between the deletion length and the editing efficiency. Despite numerous attempts with diverse sgRNAs, five segments (Δ4, Δ12, Δ26, Δ40, and Δ62) ranging from 5.6 to 22.7 kb could not be deleted, suggesting that certain non-essential genes or the combination of non-essential genes could not be knocked out, probably attributed to their metabolic connections with cell viability. Taken together, we have yielded a genome-wide library consisting of 70 CSD strains, including 60 strains that were obtained in this work and ten strains that we reported earlier.

**Fig. 2.**
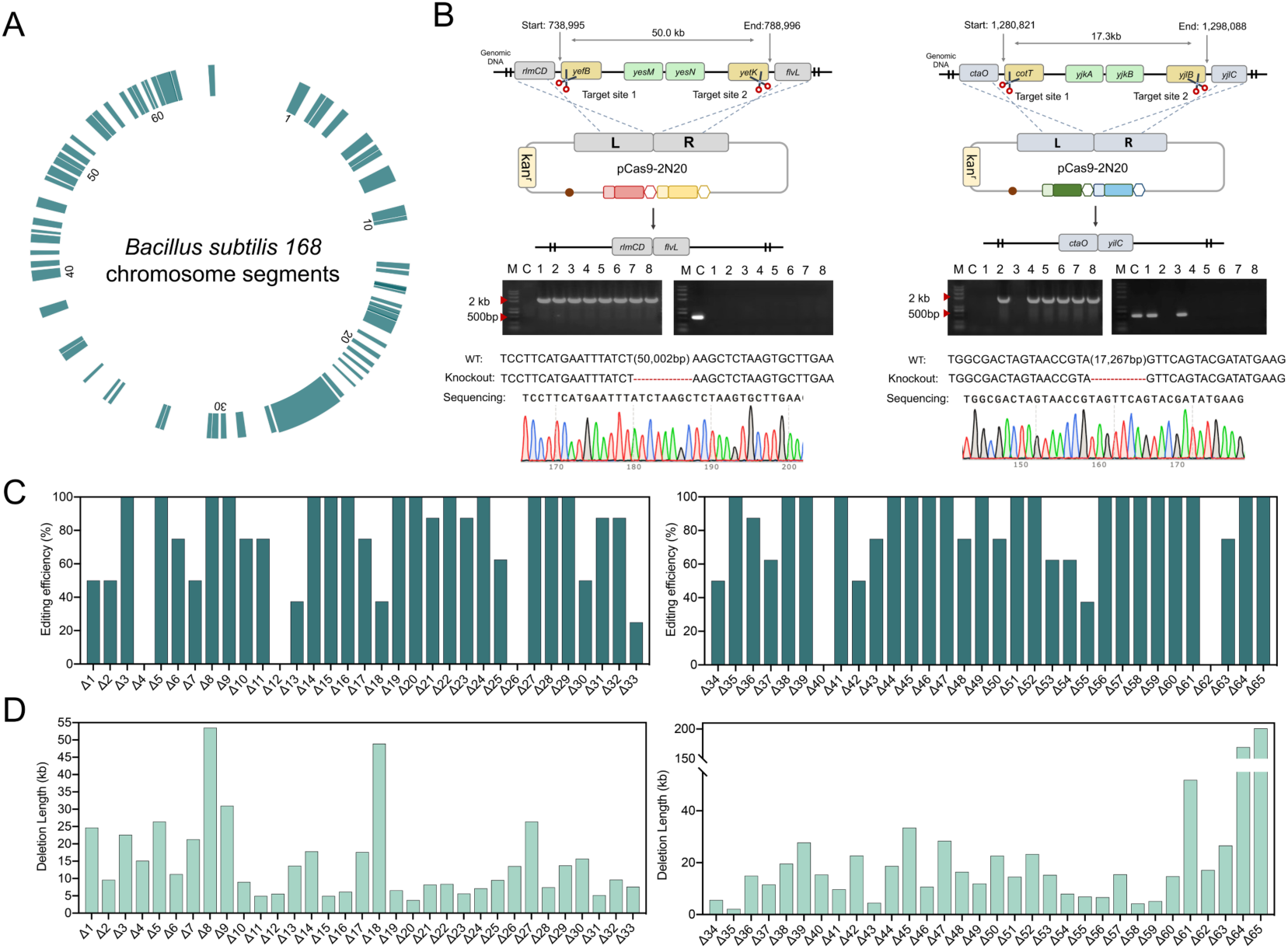
Design and construction of a genome-wide chromosome-segment-deletion library in *B. subtilis* 168. (a) 65 chromosome segments divided by essential genes. (b) Design and validation of 50.0 kb- and 17.3 kb-chromosome-segment-deletion. (c) The editing efficiency of 65 chromosome-segment-deletions. (d) The size of the 65 deleted segments constructed in this study.

### 3.4 Bioinformatic analysis of the genome-wide deletion library

To evaluate whether this library is large enough to be the representative of the whole strategy, we performed bioinformatic analysis for the deleted chromosome segments. The deleted genomic regions of this library contain 1301 *Bacillus* genes which could be grouped to 22 COG categories, among which carbohydrate metabolism and transport (G, 11.5%), transcription (K, 8.5%), amino acid metabolism and transport (E, 7.7%), cell wall/membrane/envelop biogenesis (M, 5.5%), and inorganic ion transport and metabolism (P, 5.0%) rank the top five categories with annotated functions (Fig. 3a). By comparing the number of COGs in deleted regions with that of undeleted regions, we identified that the highest proportion of deleted COGs appeared in the G category (Fig. 3b and Fig. S1). This suggested that the cumulative deletion of above segments may generate a profound effect in the carbohydrate metabolism and transportation. Further characterization of the deleted segments showed that each segment contains COGs with diverse functions (Fig. 3c). For example, both segment 5 and 67 contains COGs of six functional categories, while the categories are different in these two segments.

**Fig. 3.**
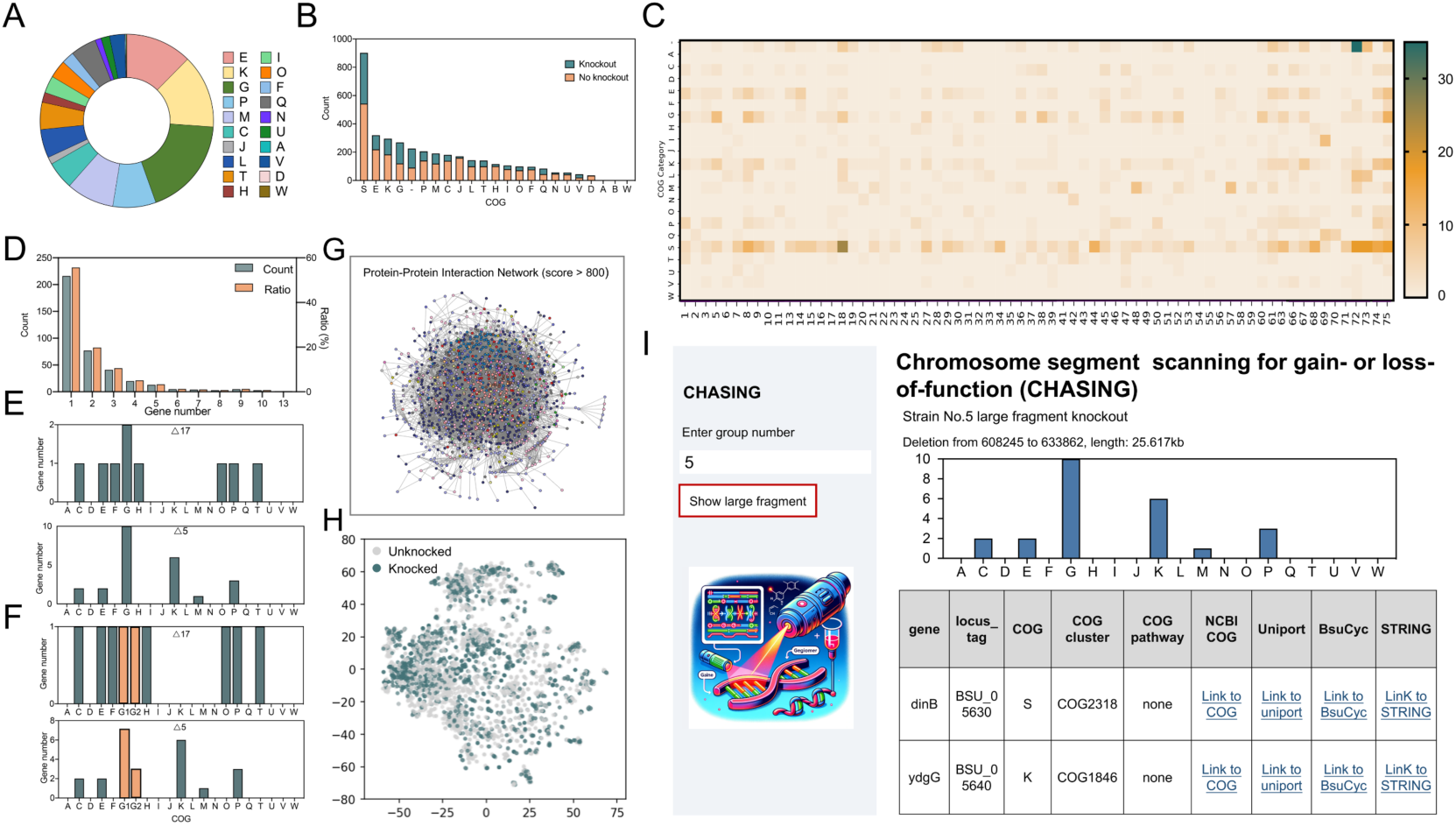
Bioinformatic analysis of the deleted chromosome segments in *B. subtilis* 168. (a) Characterization of COG categories for all deleted genes in the genome-wide deletion library. (b) The number of un-deleted and deleted genes in each COG category. (c) COG distribution pattern in each deleted segment. (d) Accumulated COG categories with different gene numbers in *B. subtilis* 168. The data was calculated based on the COGs in all deleted chromosome segments. (e) Gene number analysis of each COG category in segments 17 and 5. (f) Gene number analysis of each COG category in segments 17 and 5 by partitioning the G category into the category of carbohydrate metabolism (G1) and the category of carbohydrate transport (G2). (g) The interaction network of the proteins translated from the deleted genes. All available data of protein-protein interactions in *B. subtilis* were retrieved from the STRING database. Data of protein-protein interactions with a combined score greater than 800 were considered reliable. Each point represents a protein and the lines connecting the points depict the interaction. (h) The functional distribution of proteins in a two-dimensional space. The high-dimensional data was embedded onto a two-dimensional space by the t-SNE projections. The distance of inter-points reflects the function similarity. Gray and green points represent the proteins encoded by the undeleted and deleted genes, respectively. (i) The web server of CHASING. The COG distribution profile of each deleted segment of *B. subtilis* was displayed. With the serial number of the chromosome segment as input, the deleted genes along with the corresponding COG category would be presented. The associated metabolism of each deleted gene and the protein interaction network could be obtained by clicking the relevant web links. The schematic diagram on the left side of the web server was generated using chatGPT-4.

Next, we analyzed the COGs distribution profile in the deleted segments. Similar to that of the whole-genome analysis, the probability of having one or two genes belongs to a random COG category in a random segment is 55.7% or 19.8%, respectively (Fig. 3d). As examples, the probabilities of having one or two genes in the corresponding COG categories of segment 17 and segment 5 are 100% and 50%, respectively (Fig. 3e and Fig. S2). Theoretically, if the genes along each chromosome segment could be re-assigned into PCOF with highly diversified function categories, then the probability of associating a few genes to a certain phenotype would be increased. As a proof-of-concept, we partitioned the G category into two subcategories, *i.e.* the category of carbohydrate metabolism (G1) and the category of carbohydrate transport (G2). We observed that the probability of allocating one gene to a specific category of segment 17 was increased from 87.5% to 100% (Fig. 3f). The probability of allocating no more than three genes to a specific category of segment 5 was also increased from 66.6% to 71.4%. Therefore, the CHASING strategy could be extensively exploited by using PCOFs with more diversified categories.

To further explore the genomic features of this large-scale CSD library, we evaluated the effects of the deleted genes on metabolic network. By retrieving the available data of protein-protein interactions in *B. subtilis* from the STRING database (https://string-db.org/), we established the interaction network for the proteins translated from the deleted genes. Proteins with closer distance to the center are presumed to have a higher degree of interaction within the network and are likely to be involved in biological processes associated with the central metabolism. The uniform distribution of proteins in this map represented the varied level of participation of the deleted genes in the interaction network (Fig. 3g). A highly crosstalk has been observed in the whole protein map. The simultaneous deletions of several nodes are unlikely to cause a catastrophic effect on the network due to the function crosstalk among remaining genes in the absence of the deleted genes, reflecting the high robustness of our deletion library. This was also evidenced by the highly interconnection of proteins which transcribed from the genes within each segment (Fig. S3). To further depict the protein distribution profile in the whole network, we performed the t-Stochastic Neighbor Embedding (t-SNE) projections of protein-protein interaction from high-dimensional space onto a two-dimensional space, where the inter-point distance reflects the function similarity (Fig. 3h). The points aggregated together reflected the proteins with closely related functions. Proteins encoded by the deleted genes (green points) and the remaining genes (gray points) exhibited a uniform distribution in the map, indicating that no significant bias existed in designing the deletion segments. To be noted, the green and the gray points overlapped to some extent in the plot, suggesting that the construction of the CSD library yielded interference on the whole network of *B. subtilis*.

To assist the application of the CHASING strategy in *B. subtilis*, we developed a web tool (https://chasing.cloudmol.org/) to visualize the potential targets of each chromosome segment associated with a particular function. As shown in Fig. 3i, the COG distribution pattern along with the deleted genes and the corresponding COG categories would be reported after inputting any segment number of our library. The web tool also presented useful information for each deleted gene by providing the relevant web links, such as the associated metabolism and the protein interaction network. The potential candidates responsible for an interested phenotype could be preliminarily analyzed by dissecting the genes of the corresponding functions. This web tool will highlight the potential of our CSD library and the CHASING strategy in metabolic engineering applications.

### 3.5 Screening the genome-wide library for loss-of-function phenotype

To demonstrate the utility of this library in rapidly locating the target genes for loss-of-function, we screened the library for sporulation-deficient phenotype. The observation of morphological growth in LB medium showed that sporulation was significantly impaired in deletion strains Δ5, Δ15, Δ24, and Δ34 (Fig. S4). By analyzing the absent segments of these strains using our web tool, we preliminarily identified the promising targets of sporulation-related COGs in these deletion strains. For example, the deleted segment in strain Δ24 contains *sigE* and *sigF*, in agreement with the previous report that deletion *sigE* and *sigF* severely affected the formation of spores(Ju, et al., 1998; Rodríguez Ayala, et al., 2020). These two genes encode the critical sigma factors that control the transcription of sporulation genes. The deleted segment in Δ5 or Δ15 contains the spore cortex lytic gene *spoL*, and the coat assembly and spore resistance gene *spoVIF*, respectively, likely to be the only gene responsible to the loss-of-sporulation phenotype. The deleted segment in Δ34 contains a sporulation gene cluster, including *spoIIIAA*, *spoIIIAB*, *spoIIIAC*, *spoIIIAD*, *spoIIIAE*, *spoIIIAF*, *spoIIIAG*, and *spoIIIAH*.

Then we evaluated the survival capability of the resulting 70 strains in LB medium. The majority of the deletion strains exhibited comparable growth rates to that of the wild-type strain, suggesting that the large-scale deletion of non-essential genes did not show significant impact on cell growth in nutrient-rich medium (Fig. 4a). As shown in Fig. 4b, the specific growth rates of the deletion strains ranged from 0.107 to 0.148 h^-1^ within 24 h and the specific growth rate of wild-type strain was 0.142 h^-1^. Several strains, such as Δ15, Δ24, Δ34, and Δ39, showed retarded growth after 12 hours of cultivation, resulting in low 24-h specific growth rates, *i.e.* 0.121, 0.118, 0.113, and 0.107 h^-1^, respectively. Interestingly, three of the four slow-growing strains (Δ15, Δ24 and Δ34), have sporulation-deficient phenotypes, suggesting that spore formation in *B. subtilis* is closely related to the growth in the late stationary phase.

**Fig. 4.**
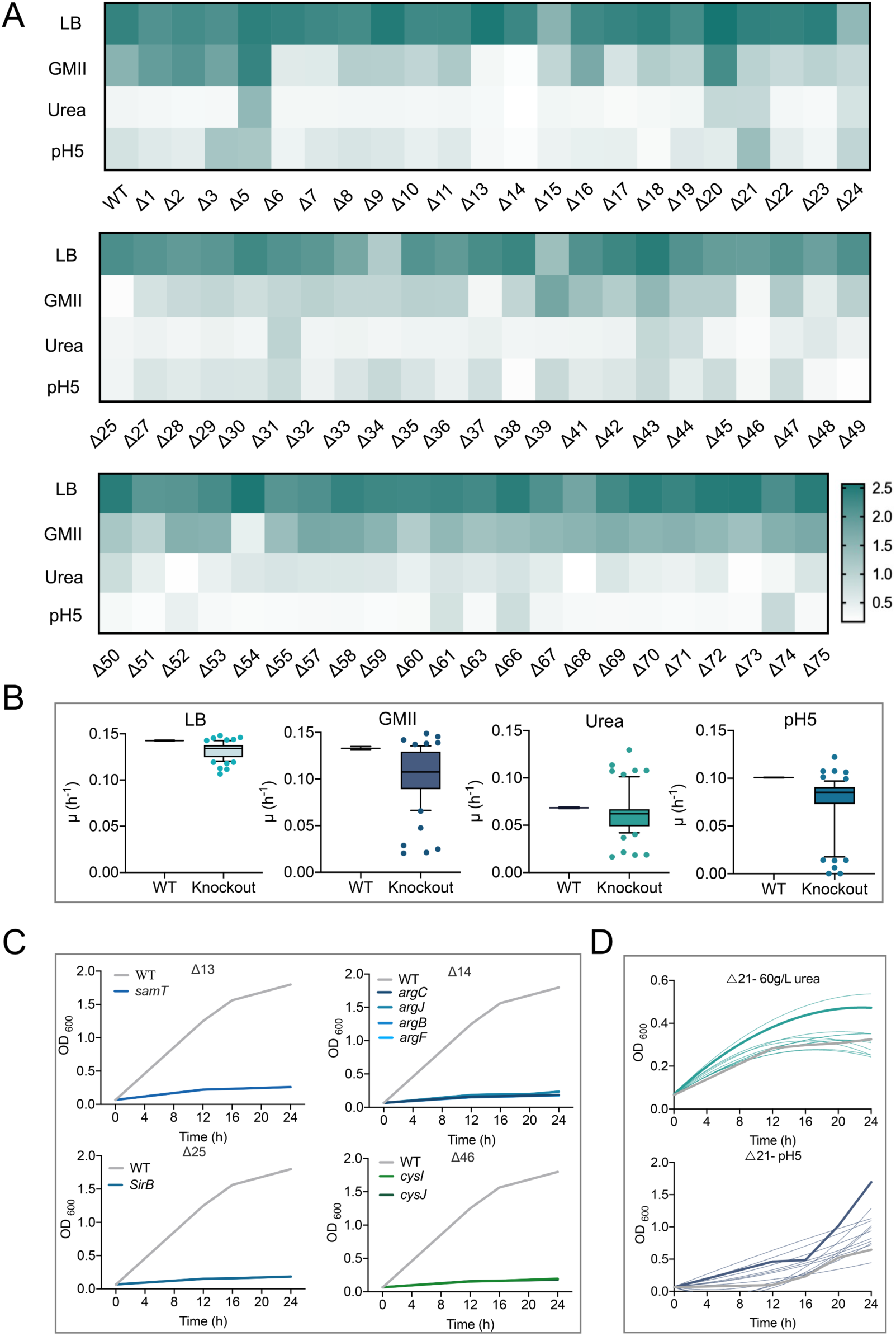
Cellular fitness of the genome-wide deletion library and genes identification for the interested phenotypes. (a) Heatmap representation of morphological patterns of chromosome-segment-deletion strains in LB, GMII, high urea, and pH 5 medium. (b) The specific growth rates of the deletion strains in LB, GMII, urea and pH 5 medium. The best and the worst growth rates were also displayed in the scatter plot. (c) The growth curves of single-gene-deletion strains in GMII medium. Genes responsible for the cell growth were identified for the strain Δ13, Δ14, Δ25, and Δ46, respectively. (d) The growth curves of the strain Δ21 and the corresponding single-gene-deletion strains under high urea and pH 5 condition.

Genome-wide identification of conditionally essential genes is of great significance in comprehending metabolic pathways and network regulations. To this end, we screened the library in GMII medium to identify the genes essential for cell growth under nutrition-poor condition. Compared with the wild-type strain, the majority of the deletion strains exhibited normal growth in GMII medium, displaying an OD_600_ value of approximately 1.5 at 24 hours (Fig. 4a and Fig. S5). These strains presented rapid growth rates from 4 to 12 hours, and slow growth rates thereafter, which might be attributed to carbon source depletion. Notably, four strains, namely Δ13, Δ14, Δ25, and Δ46, showed negligible growth in GMII medium with specific growth rates less than 0.02 h^-1^, suggesting that the corresponding chromosome segment contain genes or pathways synthesizing essential substances for cell growth in nutrient-poor medium (Fig. 4b). Then we examined the COGs function in the deleted segments using our web tool and identified the genes involved in central metabolism, including *samT* that was deleted in strain Δ13, *argC*, *argJ*, *argB*, and *argF* that were deleted in strain Δ14, *sirB* that was deleted in strain Δ25, *cysI* and *cysJ* that were deleted in strain Δ46 (Table S1). These genes were involved in the metabolism of homocysteine, arginine, cysteine, and siroheme, respectively. To verify whether these genes were conditionally essential, we performed the single-gene knockout experiments for these candidates individually. As expected, all the eight single-gene knockout strains could not grow in GMII medium, reproducing the phenotypes of the corresponding CSD strains, respectively (Fig. 4c).

### 3.6 Screening the genome-wide library for gain-of-function phenotype

*B. subtilis* is a proficient chassis for the expression of exogenous proteins and has been commonly used in the secretion of urease for the hydrolysis of urea to NH_3_. Capacity of utilizing high concentration of urea is important for this application. However, high concentration of urea could change the osmotic pressure, causing damages to the cell membrane and proteins, and ultimately affecting the survival rate of *B. subtilis*. Although normal growth was observed for *B. subtilis* in the presence of 40 g/L and 50 g/L of urea, the cell growth was significantly inhibited in the presence of 60 g/L of urea (Fig. S6a). To improve the tolerance of *B. subtilis* to urea, we conducted the growth measurement experiments under 60 g/L urea stress using the CSD library. Notably, the strains Δ5, Δ20, Δ21, Δ31, and Δ43 exhibited better growth performance with specific growth rates of 0.130, 0.108, 0.107, 0.109, and 0.104 h^-1^, respectively (Fig. 4b and Fig. S7a). These values were significantly higher than that of wild-type strain (0.068 h^-1^), indicating their potential as excellent chassis in urea utilization.

Using the strain Δ21 as the example, we aim to identify the gene responsible to the gain-of-growth phenotype. There are eleven genes in the deleted segment of the strain Δ21 (Table S2). Among them, the six genes with known functions do not encode proteins that likely related to this phenotype (Fig. S8). Therefore, the five genes with unknown functions, *ykzR*, *ykvR*, *ykvS*, *ykzS*, and *ykvU* are probably responsible to the urea-tolerance phenotype. We then deleted the five genes individually and tested the growth of the mutants in the presence of urea. The strain Δ*ykvS* exhibited vigorous growth under 60 g/L of urea during the initial 16 h with a growth rate of 0.123 h^-1^, which was 29.5% and 6.03% higher than that of the wild type strain (0.095 h^-1^) and stain Δ21 (0.116 h^-1^), respectively (Fig. 4d). The other strains could not restore the growth of cells in the presence of 60 g/L urea. To rule out the probability that any other gene in the deleted segment of Δ21 may be related to the urea-resistant phenotype, all the six genes with known functions were knockout individually and these deletions had no effect on the cell growth in the presence of 60 g/L urea (Fig. 4d). Taken together, this result indicated that the single-gene deletion of *ykvS* in the wild-type strain could reproduce the gain-of-resistance phenotype of strain Δ21 under urea-rich condition.

In industrial process, organic acids accumulated during fermentation, leading to intracellular acid shock which could not be completely overcome by extracellular pH adjustment. Hence, screening strains tolerant to organic acids accumulation-induced low pH is crucial to industrial production. To demonstrate the practicality of the genome-scale deletion library, we evaluated the survival capability of the library strains in GMII medium under acidic conditions. To this end, we first evaluated the tolerance capability of wild type *B. subtilis* at three different pH conditions, including pH 3, pH 4, and pH 5. No cell growth was observed at pH 3 and pH 4, indicating completely inhibition of growth under the extremely low pH conditions (Fig. S6b). In contrast, partially inhibited growth was observed at pH 5. Therefore, we selected pH 5 to screen the strain chassis that tolerate low pH using the CSD library. Interestingly, strain Δ5 and strain Δ21 showed increased fitness under pH 5 condition compared with the wild-type *B. subtilis*, reaching OD_600_ of 1.20 and 1.69, respectively (Fig. 4a). Since the single gene-knockout strains of Δ21 have been constructed in above experiments, we directly measured the growth curves of these strains under pH 5 condition. Among these strains, strain Δ*ykvQ* that delete the gene encoding a putative sporulation-specific glycosylase exhibited a growth curve similar with strain Δ21, reaching a OD_600_ of 1.26 at 24 h under acidic conditions (Fig. 4d and Fig. 7b). In addition, the deletion of other sporulation-related genes such as *ykvP* and *ykvT*, which encode spore protein and cell wall hydrolase related to spore cortex-lytic enzymes, respectively, also resulted in enhanced growth fitness than wild type. These results suggested that the cellular fitness in strain Δ21 was probably introduced by the synergistic deletion effect of multiple sporulation genes, highlighting the potential of large-scale deletion library in screening robust chassis.

### 3.7 Potential mechanisms underlying the improved cellular fitness

To visualize the variance in chassis fitness, we performed principal component analysis (PCA) by plotting the growth dataset in different conditions to two most descriptive principal components (Fig. 5a). In the PCA plot, the variable points were scattered around the origin and were grouped into four categories with different growth patterns, showing significantly varied fitness of the CSD strains. By projecting each data point to the first principal component, we identified three groups with distinct growth patterns. Group one contains the strain with the highest growth fitness under GMII, pH 5 and urea-rich conditions, *i.e.* the strain Δ5. Group two consists of strains Δ21, Δ43, Δ20, Δ31, and Δ3, which displayed better growth fitness than the wild-type strain (red dot in PCA plot) but inferior pattern than strain Δ5. On the contrary, group three included the strains that showing the worst growth under GMII, pH 5 and urea-rich conditions. By projecting each data point to the second principal component, it was easily to identify the group four, in which the strains showed inferior growth pattern under nutrient-rich condition such as LB medium.

**Fig. 5.**
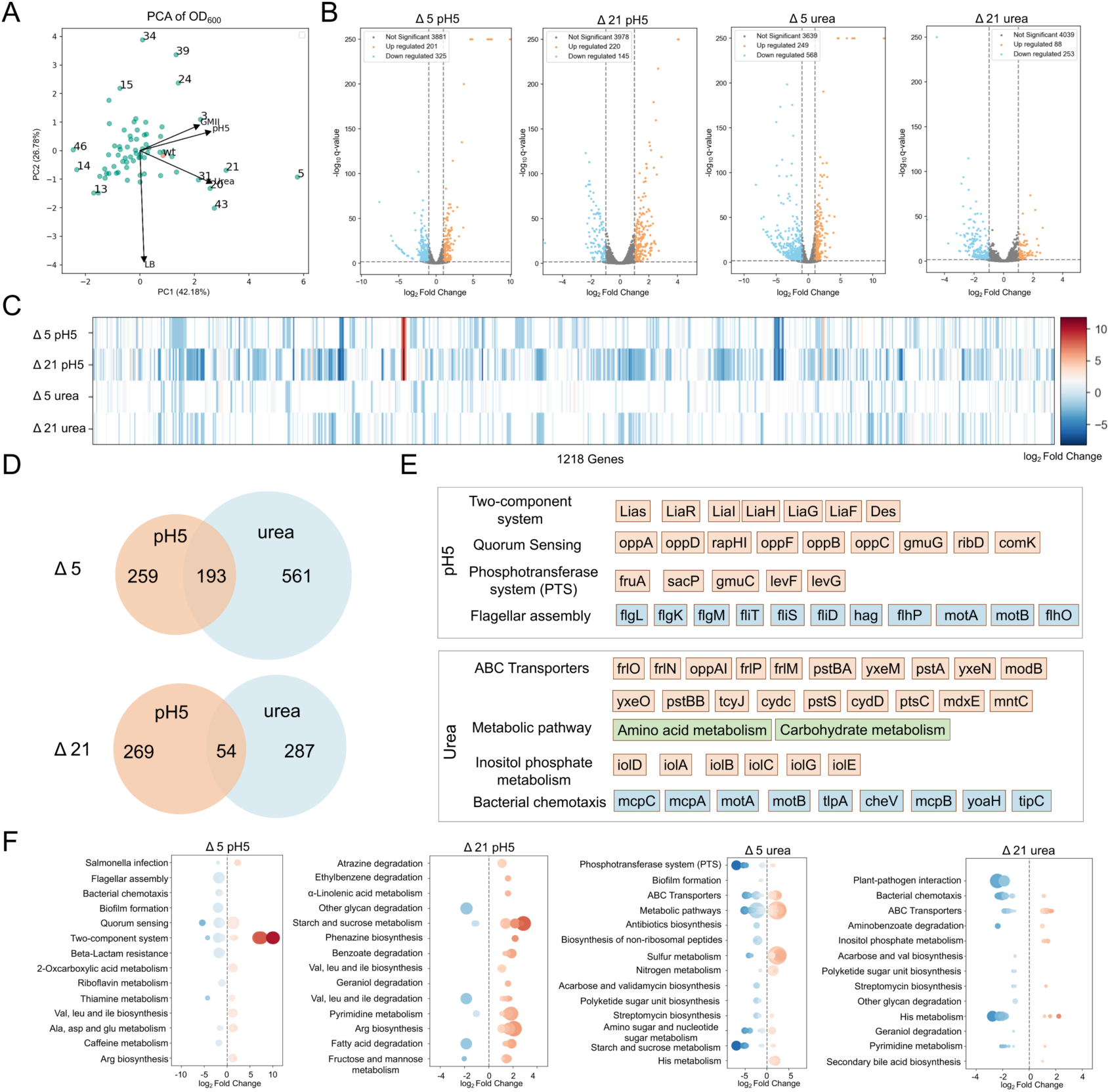
Transcriptome analysis of the strains Δ5 and Δ21 under pH 5 and high urea condition. (a) Principal component analysis (PCA) of the growth fitness under different conditions. (b) Volcano plot of transcriptomic data of the strains Δ5 and Δ21 under pH 5 or high urea conditions. The blue dots represent the downregulated genes and the orange dots indicate the upregulated ones. The log_2_ fold-change in differentially expressed genes was compared to the wild-type *B. subtilis*. The cutoff of the differentially expressed gene was set at a log2-fold change > |1| with a q-value (FDR, padj) < 0.05. (c) Heat map of upregulated and downregulated genes under pH5 and high urea condition. (d) Venn diagram representation of the differentially expressed transcripts that are specific for pH 5 and high urea condition. The orange and blue colors represent the condition of pH 5 and urea, respectively. (e) The major pathways with the largest number of differentially-expressed genes. The pathways in upregulation and downregulation were indicated in orange and blue color, respectively. The pathways with both upregulated and downregulated genes were indicated in green color. (f) Scatter plot of differentially expressed genes in KEGG enrichment. Larger size of the dot represents higher number of differentially expressed genes. Deeper color of the dot represents higher fold-change in differential expressed genes.

To illustrate the potential mechanisms for chassis robustness, we performed the transcriptome analysis by selecting the strain Δ5 in group one and the strain Δ21 in group two as representatives, identifying the differentially expressed transcripts under pH 5 and urea-rich conditions, respectively (Fig. 5b, c, Note1). To exclude the effect caused by the deletion of unrelated non-essential genes, we further performed the cluster analysis for the strain Δ5 and Δ21, respectively, identifying the differentially expressed transcripts and the corresponding metabolism that are specific for pH 5 (Fig. 5d) and urea-rich condition (Fig. 5e), respectively. For the strain Δ5, the most affected metabolism under acid shock is the two-component system and quorum sensing system (Fig. 5f), among which the LiaRS system (Jordan, et al., 2008) and the Opp system (Liu, et al., 2017)have been reported to be involved in the adaptive responses to the pH stress. These results suggested that the improved fitness in strain Δ5 may be acquired by upregulating the genes of these pathways, leading to a fast and differentiated response under pH 5 condition. Under urea shock, the most affected metabolism in the strain Δ5 are amino acid and carbohydrate metabolism (Fig. 5e, f). This could be explained by the feedback inhibition of urea on the production of NH_4_^+^ and the degradation of amino acids, leading to the rewiring of amino acid and carbohydrate metabolism(Chandel, 2021).

To be noted, the metabolic changes were totally different in the strain Δ21 (Fig. 5e, f). For example, the most significantly upregulated and downregulated genes under pH 5 condition are the starch and sucrose metabolism, and glycan degradation pathways, respectively. This could be attributed to the deletion effect of putative sporulation-specific glycosylase gene *ykv*Q, which is responsible for the enhanced tolerance of the strain Δ21 to acid shock (Fig. 4d). It has been well documented that glycosylase play important roles in the degradation of glycan(2009). It is possible that the deletion of *ykv*Q resulted in an upregulated level of glycan in Δ21, increasing the abundance of lipopolysaccharide and the toughness of cell wall, thus leading to an improved response to acid stress. Under urea-rich condition, the most significantly affected genes were associated with the plant-pathogen interaction pathway and His metabolism. Based on this, the absence of hypothetical YkvS protein may enhance the urea tolerance via these pathways.

### 3.8 Apply the genome-scale deletion library to screen for strains with increased plasmid stability

Plasmids could introduce genetic parts to cells, thus offering a powerful tool for biotechnological applications. However, introducing exogenous plasmids often causes metabolic burden and extra fitness costs, inducing the protective mechanisms of the cells such as the interference in plasmid replication and the clearance of plasmid. To address this, we focused on the strain chassis and aimed at deleting the factors that affect plasmid stability. To this end, we established a high-throughput report system for evaluating plasmid stability using a plasmid that carrying repA origin and sfGFP reporter. Each strain in the CSD library was individually transformed with the sfGFP plasmid and was successive cultivated without antibiotics selection for three generations (Fig. 6a). After each transfer, the fluorescence of each strain was detected at 8 h for calculating the plasmid preservation rates. After the first transfer, the plasmid preservation rates ranged from 55.8% to 96.8% (Fig. 6b). Most strains exhibited similar fluorescence intensities with that of the wild-type *B. subtilis*, except the strains Δ5, Δ11, Δ49, and Δ71 showing low fluorescence intensity. However, after the second transfer, the plasmid preservation rates decreased significantly in most strains, with the wild type strain exhibiting a preservation rate of 4.0%. This indicated that the plasmid was difficult to sustain during the successive cultivation without antibiotics selection. In contrast, the strains Δ1, Δ27, Δ28, Δ44, Δ52, Δ65, Δ66, and Δ70 displayed high fluorescence intensity after the second transfer, reaching an average preservation rate of 48.9%. Even after the third transfer, the strains Δ28, Δ44, Δ52, Δ65, Δ66, and Δ70 still displayed high plasmid preservation rates, *i.e.* 18.4%, 29.5%, 13.5%, 18.3%, 15.2% and 25.1%, respectively. These strains were termed as the strain chassis with enhanced plasmid stability.

**Fig. 6.**
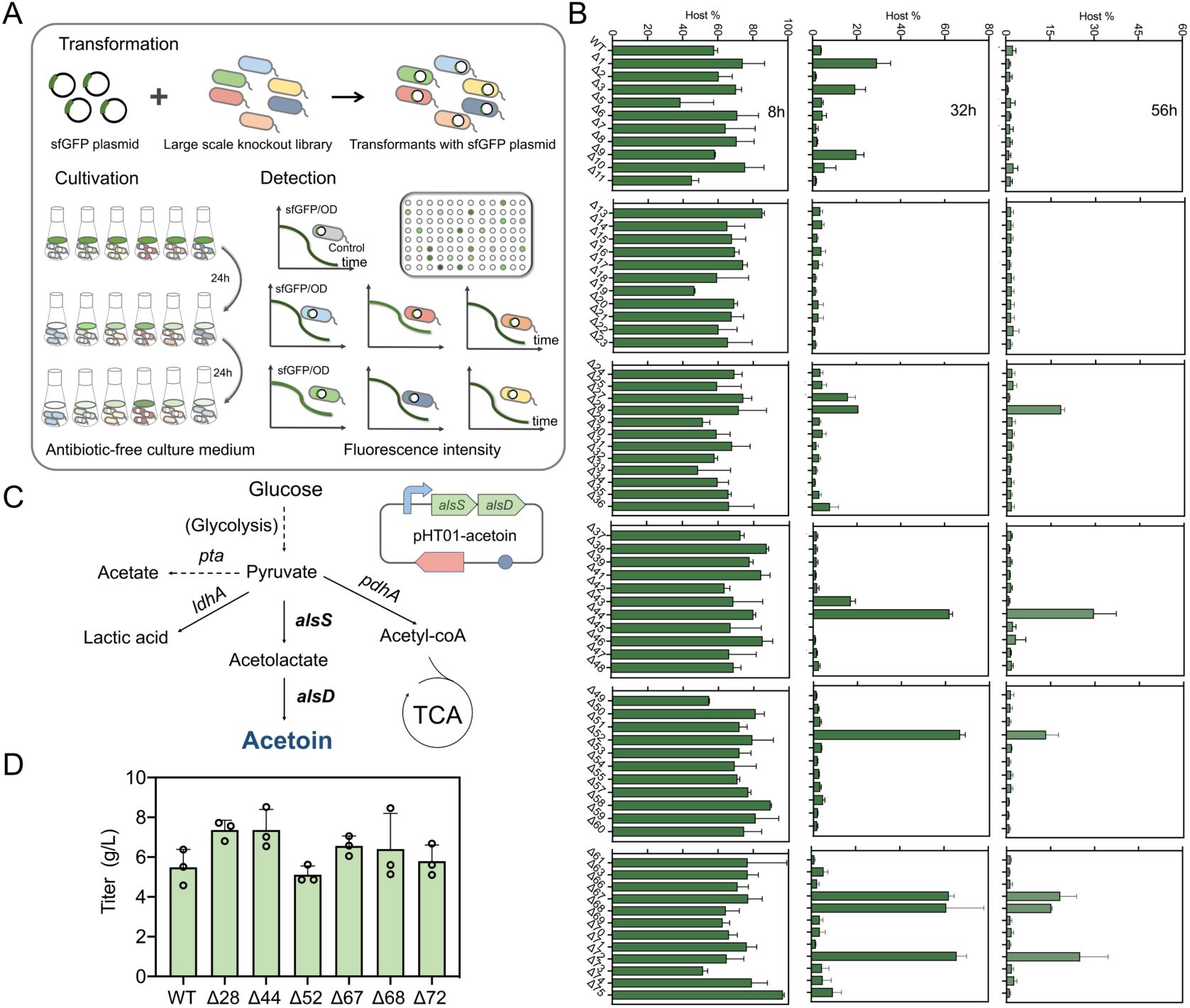
Plasmid stability of the CSD strains under non-selective condition. (a) Schematic diagram of the strain screening for plasmid stability using a high-throughput system. Subculturing of each sfGFP-harboring strain was performed every 24 h and the fluorescence was measured at OD_474_. Higher fluorescence intensity represents higher plasmid stability. (b) Plasmid stability analysis of the deletion strains. Sample was collected at 8, 32, and 56 h, respectively, and the fluorescence was measured at OD_474_. (c) Metabolic pathway of acetoin production in *B. subtilis*. *alsS*, acetolactate synthase; *alsD*, acetolactate decarboxylase. (d) Acetoin production of the deletion strains with higher plasmid stability. The fermentation was performed in LB medium without antibiotic. All data was represented using the average value of three independent experiments.

To demonstrate the significance of these strains in plasmid maintaining and chemical production, we applied the screened-out strains to acetoin production using an antibiotic-free system. The acetoin biosynthetic genes, *alsS* and *alsD*, were expressed in this plasmid instead of the reporter gene *sfGFP* (Fig. 6c). As expected, the acetoin production in the strains with enhanced plasmid stability were improved. Specifically, the strains Δ28, Δ44, Δ52, Δ65, Δ66, and Δ70 reached titers of 7.36, 7.36, 5.11, 6.56, 6.40, and 5.80 g/L, respectively, which were higher than the 5.49 g/L titer in the wild-type *B. subtilis* (Fig. 6d). These results showed that the chassis with high plasmid stability is also a potential host for chemical production.

### 3.9 Apply the genome-scale deletion library to screen for strains with improved lycopene production

In the present study, the food colorant lycopene, which possess antioxidant, anti-aging, and anti-inflammatory properties, was chosen as a target product to show the potential of genome-scale deletion library in chemical production. Although *B. subtilis* harbors the capability to synthesize intermediate metabolites of lycopene (IPP and DMAPP) and is able to produce GPP and FPP through *ispA* catalysis, it lacks the pathway toward lycopene production (Fig. 7a). To establish a complete biosynthetic pathway of lycopene in *B. subtilis*, genes including *crtE*, *crtB*, and *crtI* from *Pantoea ananatis* were cloned and expressed. Then, the lycopene production plasmid expressing *crtEBI* was transformed into the strains of CSD library individually. After fermentation, the culture color of each strain was compared with that of the wild-type *B. subtilis* and mutant strains with a significantly deeper color were further subjected to lycopene extraction and yield calculation. Although most mutant strains exhibited similar color depth with the control strain, the mutant strains Δ27 and Δ28 had a notable increase in lycopene production and the yield reached 4.12 and 3.24 mg/g dry weight, respectively, representing 47.7% (Fig. 7b) and 16.1% (Fig. 7c) increase in yields, respectively.

**Fig. 7.**
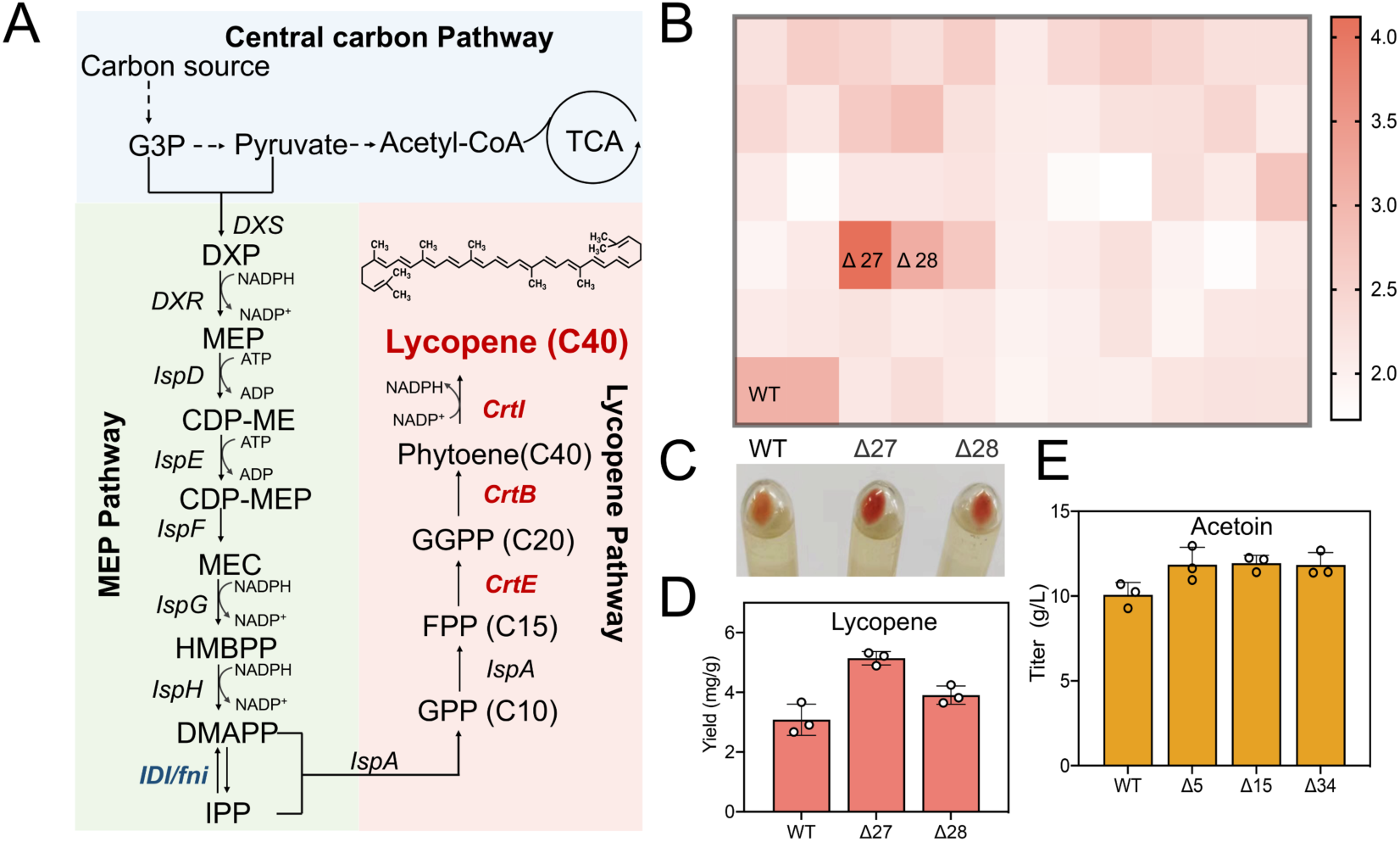
Application of the genome-wide deletion library in metabolic engineering. (a) The engineered pathway of lycopene biosynthesis in *B. subtilis*. The genes in red color are lycopene biosynthetic genes, including the heterologous genes encoding geranylgeranyl diphosphate synthase (*CrtE*), phytoene synthase (*CrtB*) and phytoene desaturase (*CrtI*) from *Pantoea ananas*. (b) Heatmap representation of lycopene production in the deletion strains that overexpressing lycopene biosynthetic genes. (c) The cell pellets of lycopene fermentation samples from the strains Δ27 and Δ28 overexpressing lycopene biosynthetic genes. (d) Lycopene production in the superior chassis overexpressing lycopene biosynthetic genes and the rate-limiting gene *fni*. (e) Acetoin production in the sporulation-deficient strains.

To make full use of above chassis for lycopene production, we have tried to increase the supply of intermediate IPP by overexpressing the biosynthetic enzymes. We first transformed the production plasmids overexpressing the endogenous genes of potential rate-limiting enzymes, *dxs* and *fni* (a homolog of *idi* from *E. coli*), into the wild-type *B. subtilis*. By measuring the yield, we identified that the overexpression of *dxs* did not have a significant impact on lycopene production whereas the overexpression of *fni* increased the yield by 43.8%. Leveraging the best strain chassis, we overexpressed the *fni* gene in strains Δ27 and Δ28. The shake-flask fermentation results showed that the lycopene yield in these two strains was further increased to 5.27 and 4.04 mg/g dry weight, respectively, reaching 54.1% and 18.2% increase in yield compared to the one of the wild-type strain (Fig. 7d).

### 3.10 Improved chemical production using the screened-out sporulation-defective strains

During the fermentation process of *B. subtilis*, the nutrient becomes deficient as the fermentation prolongs and sporulation occurs as a response mechanism, inducing a state of dormancy and a decrease in production yield(Klausmann, et al., 2021). Motivated by the previous study, in which a Δ*sigE*Δ*sigF* mutant has been proved to be effective in increasing acetoin production(Q. Wang, et al., 2022), we sought to test the chemical production ability using the screened-out spore-deficient strains Δ5, Δ15, Δ24, and Δ34 (Fig. S4). These strains were transformed with acetoin-producing genes, *alsS* and *alsD*, yielding the engineered acetoin production strains. The fermentation experiments showed that Δ5, Δ15, Δ24, and Δ34 produced acetoin with titers of 11.9 g/L, 12.0 g/L, 12.2 g/L, and 11.8 g/L, respectively, while the wild-type *B. subtilis* strain only obtained a titer of 10.1 g/L at 72 h (Fig. 7f). These values represent 17.6%, 18.4%, 21.5%, and 17.4% increases in acetoin production, respectively. These results further demonstrated the believed trade-off between growth and production(Zhu, et al., 2023).

## 4. Discussion

Profiling of arrayed genome-wide deletion or interference libraries lay the foundation for the engineering of robust chassis. However, the shortage of high-throughput libraries for screening the interested phenotype and the limited understanding of the genotype-phenotype relationship hampered this purpose, especially for non-model strains. CRISPR-based technology breaks the limit in genome editing and increases the feasibility of genome-wide library construction(Bock, et al., 2022). CRISPR-based long-fragment deletion strategy has been widely reported in diverse organisms, achieving the scarless deletion of genomic fragments up to 134.3 kb, 186.7 kb, and 38 kb in *B. subtilis*(Tian, Xing, et al., 2022), *E. coli*(Huang, et al., 2020), and *Saccharomyces cerevisiae*(Hao, et al., 2016; Li, et al., 2018), respectively. In the present study, we employed CRISPR/Cas9-based long-fragment deletion technique to generate assembled genes knockout strains for phenotyping, successfully establishing an arrayed CSD library covering 31.6% of the *B. subtilis* 168 genome and 32.4% of the non-essential genes. The CRISPR-based assembled genes knockout strategy could also be applied to other microorganisms.

To give an overview of the CHASING, Keio collection and the CRISPRi methods, we compared the three aspects including library construction, screening and application. For library construction, the cost savings were calculated by measuring the primer costs for constructing the deletion strains, including that for amplifying N20 and homologous arm as well as for sequencing. Compared with the Keio collection, the CHASING strategy exhibited great cost- and time-savings in building an arrayed library (Table 1, Table S6). Only 70 strains besides the controls were involved, making the screening scale fitted well to the 96-well plate and the process easy to manipulate. To be noted, our library also assists the pooled screening for some particular applications. By designing the chromosome segment-specific PCR primers, the deleted segment in the screened-out mutant could be rapidly identified by PCR reactions. This makes our library more versatile and time-saving in applications.

**Table 1.**
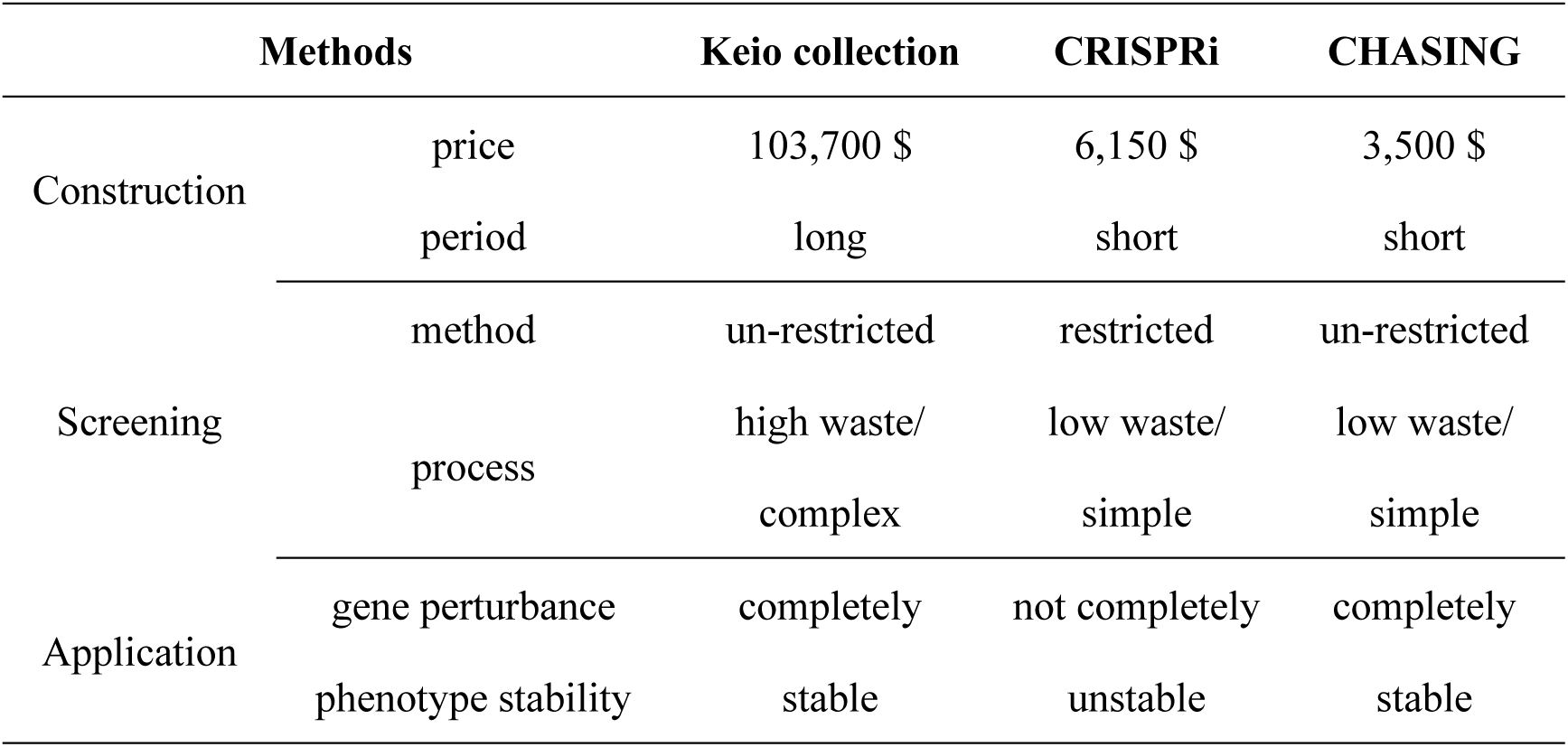
Duration and Minimal cost of realizing the phenotype-genotype mapping with a Keio collection, a CRISPRi array, or a CHASING library using the state-of-the-art technologies.

The cost savings of CHASING strategy is comparable to that of CRISPRi taking account of the genome coverage of the arrayed CSD library (Table 1). The advantages of CHASING mainly included the un-restricted screening method, the phenotypically stable strains and the accurate perturbance level in gene expression as compared with the CRISPRi method. Beyond this, the arrayed CSD library is amenable to single-pot assay in contrast to the pooled CRISPRi library, thus providing precise readout in experiments such as growth measurements and morphology phenotyping.

To assist the genotyping of interested phenotypes, we designed a user-friendly web server to scan the COG distribution profile of the deleted chromosome segment in *B. subtilis*, and showed that the probability to link a single gene directly to an observed phenotype is 55.7%. Regarding the simplified COG category, some functions such as stress resistance, chemical production, or sporulation are not assigned to any specific category of COG family, leading the corresponding genes be shadowed using the CHASING strategy. This could be addressed by generating a PCOF with more diversified categories by incorporating more protein databases such as the InterPro database (https://www.ebi.ac.uk/interpro/) into the PCOF. Another challenge is the huge number of hypothetical and unknown genes in the genome, and especially the difficulties to annotate them using homology-based approaches. With the explosion of protein sequences and the mature of AlphaFold database, this might be addressed by leveraging the machine learning revolution in protein bioinformatics(Durairaj, et al., 2023). For every chromosome segment, more specified categorization generated a higher probability of linking one gene to an interested phenotype (Fig. 3f). Therefore, the advance of cutting-edge technologies and the advent of interactive resources give promise into employing CHASING strategy for rapid genotyping. As an available accessory, our web server could be further upgraded by incorporating the renewed PCOF.

It should be noted that about 34% of the accumulative COG categories in the *Bacillus* genome contain more than two genes, some of which were annotated into a gene cluster. This yielded cases such that a robust phenotype of a particular strain may not be reproduced by the single-gene deletion due to the synergistic effect of multiple genes. For example, we found that the cellular fitness of strain Δ21 under pH 5 condition could not simply be reproduced by single-gene knockouts. Such a robust phenotype may be overshadowed using the single-gene perturbation library owing to the gene redundancy effect. In this regard, the CHASING strategy shows a unique potential in screening the robust chassis.

Eliminating a large quantity of non-essential genes significantly affect the transcription and translation in the cell, thus adjusting the resource allocation and reducing the metabolism complexity for diverse applications(KimOh, 2023; Reuß, et al., 2017; Zhang, et al., 2023). Regarding the complicated network interconnections (Fig. 3g), future cumulative deletion of different chromosome segments may have profound effects on the strain metabolism. In this regard, our library provided versatile resources for yielding diverse combinations of the CSD strains, offering promise to gain more comprehensive knowledge toward cellular life and interactive metabolism beyond the currently existed genome-minimized strains.

The applicability of CHASING strategy was further evaluated by analyzing the genome of *E. coli* and *B. thuringiensis*. For *E. coli*, the whole genome could be divided into 184 large segments by essential genes (Fig. S9). The probability of having one or two genes belongs to a random COG category in a random segment is 45.8% or 20.8%, respectively (Fig. 1e). For *B. thuringiensis*, the whole genome could be divided into 437 large segments by essential genes (Fig. S10). The probability of having one or two genes belongs to a random COG category in a random segment is 64.6% or 19.8%, respectively (Fig. 1f). These results further demonstrated the applicability and generality of the CHASING strategy.

Taken together, we presented a simplified CHASING strategy to establish a systematic arrayed library by assembled gene knockout. An arrayed genome-wide library was established in *B. subtilis* and was successfully applied to obtain strains with stress-tolerance, enhanced plasmid stability, sporulation mutants, and improved chemical production. Our work demonstrated the effectiveness of this platform in genome-wide network perturbation and high-throughput function screening. Our CHASING strategy shows great potential in screening the interested phenotype and illustrating the genotype-phenotype relationship.

## Data availability

Data associated with this research could be accessed at NCBI under BioProject PRJNA1031996. The transcriptomic data of strains under pH 5 condition could be found under BioSample SAMN37977971 (the first parallel sample of strain Δ5), SAMN37977972 (the second parallel of strain Δ5), SAMN37977973 (the first parallel sample of the strain Δ21), SAMN37977974 (the second parallel sample of the strain Δ21), SAMN37977975 (the first parallel sample of the wild-type), and SAMN37977976 (the second parallel sample of the wild-type), respectively. The transcriptomic data of strains under urea-rich condition could be found under BioSample SAMN37977977 (the first parallel sample of strain Δ5), SAMN37977978 (the second parallel sample of strain Δ5), SAMN37977979 (the first parallel sample of the strain Δ21), SAMN37977980 (the second parallel sample of the strain Δ21), SAMN37977981 (the first parallel sample of the wild-type), and SAMN37977982 (the second parallel sample of the wild-type), respectively. The authors declare that the data are available upon request.

## Acknowledgements

This work was supported by the National Key R&D Program of China (2021YFC2100500, and 2019YFA0904104) and the Beijing Natural Science Foundation (grant number 2222026). Part of the experiments was carried out in the Biological & Medical Engineering Core Facilities of the Beijing Institute of Technology.

## Author contributions

S.Y.G., and Y.-X.H. generated the idea. S.Y.G., Y.-X.H., Y.X. and L.C.S. conceptualized the project. Y.X., Z.Y.L., Z.R.H., J.L., Y.J.G., and P.Y.D. carried out the experiments. Y.X. completed bioinformatics analysis and website development. Y.X., and L.C.S. analyzed the experiments and transcriptome data, and wrote the manuscript. S.Y.G., Y.-X.H., Y.X., and L.C.S. revised and edited to obtain the final manuscript.

## Competing interests

The authors declare no competing interests.

## Notes

### Competing Interest Statement

The authors have declared no competing interest.

