## Supplemental information for "Chromosome segment scanning for gain- or loss-of-function screening (CHASING) and its application in metabolic engineering"

**Table S1. The genes information of the strain Δ21.**

| Gene | Position | Function | COG category |
|---|---|---|---|
| *ykvN* | 1,442,347 - 1,442,703 | putative transcriptional regulator (HGT island) | K |
| *ykvO* | 1,442,872 - 1,443,618 | putative oxidoreductase (HGT island) | I |
| *ykvP* | 1,444,099 - 1,445,298 | spore protein (HGT island) | M |
| *ykzQ* | 1,445,314 - 1,445,541 | putative peptidoglycan binding protein | M |
| *ykvQ* | 1,445,638 - 1,446,336 | putative sporulation-specific glycosylase (HGT island) | G |
| *ykzR* | 1,446,317 - 1,446,568 | putative spore-specific glycosyl hydrolase | S |
| *ykvR* | 1,447,251 - 1,447,541 | conserved hypothetical protein (HGT island) | S |
| *ykvS* | 1,447,662 - 1,447,847 | conserved protein of unknown function (HGT island) | S |
| *ykzS* | 1,448,013 - 1,448,207 | conserved hypothetical protein (HGT island) | - |
| *ykvT* | 1,448,506 - 1,449,132 | cell wall hydrolase related to sporecortex-lytic enzymes | M |
| *ykvU* | 1,449,250 - 1,450,587 | spore membrane protein involved in germination | S |

**Table S2. The genes information of the strain Δ13.**

| Fragment | Gene | position | Function | COG category |
|---|---|---|---|---|
| 13 | *yitF* | 1,174,861-1,175,976 | putative enolase superfamily enzyme | M |
| 13 | *yitG* | 1,175,985-1,177,253 | putative efflux transporter | E |

| | | | | |
|---|---|---|---|---|
| 13 | *yitH* | 1,177,365-1,178,213 | putative N-acetyltransferase | K |
| 13 | *yitI* | 1,178,218-1,178,667 | putative N-acetyltransferase | S |
| 13 | *samT* | 1,178,757-1,180,595 | bifunctional homocysteine S-methyltransferase using (R,S)AdoMet and methylenetetrahydrofolate reductase | E |
| 13 | *mswK* | 1,180,685-1,180,802 | RNA | S |
| 13 | *yitK* | 1,180,909-1,181,400 | putative RNA or cyclic d-GMP binding protein | S |
| 13 | *yitL* | 1,181,499-1,182,395 | RNA-binding protein | - |
| 13 | *yitM* | 1,182,448-1,183,032 | conserved hypothetical protein | - |
| 13 | *yitO* | 1,183,029-1,183,958 | putative integral inner membrane protein with HTTM domain | S |
| 13 | *yitP* | 1,183,943-1,184,479 | conserved hypothetical protein | K |
| 13 | *yizB* | 1,184,657-1,185,004 | putative transcriptional regulator | - |
| 13 | *yitQ* | 1,185,001-1,185,588 | conserved protein of unknown function | S |
| 13 | *yitR* | 1,185,608-1,185,901 | conserved protein of unknown function | E |
| 13 | *nprB* | 1,186,037-1,187,653 | extracellular neutral protease B | S |
| 13 | *fakBB* | 1,187,700-1,188,551 | fatty acid kinase fatty acid binding subunit B | |

| | | | | |
|---|---|---|---|---|
| 14 | *yitT* | 1,188,689-1,189,531 | putative integral membrane protein | S |
| 14 | *ipi* | 1,189,646-1,190,005 | intracellular proteinase inhibitor BsuPI | S |
| 14 | *yizC* | 1,190,036-1,190,233 | conserved hypothetical protein | S |
| 14 | *ribZC* | 1,190,490-1,191,302 | 5-amino-6-ribitylamino-2,4(1H,3H)-pyrimidinedione 5'-phosphate phosphatase | S |
| 14 | *yitV* | 1,191,423-1,192,190 | putative carboxylesterase | S |
| 14 | *yitW* | 1,192,254-1,192,562 | putative protein involved in Fe-S cluster assembly | S |
| 14 | *yitY* | 1,192,858-1,194,288 | putative FMN/FAD-binding oxidoreductase | C |
| 14 | *yitZ* | 1,194,333-1,194,827 | putative transport protein | G |
| 14 | *argC* | 1,195,034-1,196,071 | N-acetylglutamate gamma-semialdehyde dehydrogenase | E |
| 14 | *argJ* | 1,196,091-1,197,311 | ornithine acetyltransferas | E |
| 14 | *argB* | 1,197,326-1,198,102 | N-acetylglutamate 5-phosphotransferase | E |
| 14 | *argD* | 1,198,099-1,199,256 | N-acetylornithine aminotransferase | E |
| 14 | *carA* | 1,199,327-1,200,388 | arginine-specific carbamoyl-phosphate synthetase | F |

| | | | | |
|---|---|---|---|---|
| 14 | *carB* | 1,200,381-1,203,473 | arginine-specific carbamoyl-phosphate synthetase | E |
| 14 | *argF* | 1,203,461-1,204,420 | ornithine carbamoyltransferase | E |
| 14 | *yjzC* | 1,204,506-1,204,685 | conserved hypothetical protein | S |
| 14 | *yjzD* | 1,204,731-1,204,916 | forespore targeted protein | S |
| 14 | *yjaU* | 1,205,165-1,205,899 | conserved hypothetical protein | I |
| 14 | *yjaV* | 1,205,981-1,206,538 | putative NAD(P) binding enzyme | - |
| 25 | *cysP* | 1,631,095-1,632,159 | sulfate permease | P |
| 25 | *sat* | 1,632,208-1,633,356 | sulfate adenylyltransferase | P |
| 25 | *cysC* | 1,633,369-1,633,962 | adenylylsulfate kinase | P |
| 25 | *sumT* | 1,634,061-1,634,834 | uroporphyrinogen III and precorrin-1C-methyltransferase | H |
| 25 | *sirB* | 1,634,837-1,635,622 | sirohydrochlorin ferrochelatase | S |
| 25 | *sirC* | 1,635,603-1,636,091 | precorrin-2 dehydrogenase | H |
| 25 | *fbnA* | 1,636,131-1,637,849 | putative tRNA modification protein | K |
| 25 | *tcaB* | 1,637,965-1,640,637 | P-type calcium transport ATPase (sporulation) | P |

| | | | | |
|---|---|---|---|---|
| 46 | *pgoN* | 3,426,749-3,427,579 | promiscuous glyoxal/methylglyoxal reductase | S |
| 46 | *yvgO* | 3,427,802-3,428,287 | exported stress induced factor | - |
| 46 | *nhaK* | 3,428,331-3,430,343 | Na+/H+ antiporter | P |
| 46 | *cysI* | 3,430,598-3,432,313 | assimilatory sulfite reductase | P |
| 46 | *cysJ* | 3,432,339-3,434,156 | assimilatory sulfite reductase | P |
| 46 | *helD* | 3,434,327-3,436,651 | transcription factor | L |
| 46 | *yvgT* | 3,436,849-3,437,457 | putative integral membrane protein | S |

**Table S3. The strains used in this study.**

| **Strains** | **Description** | **Source** |
|---|---|---|
| *E. coli* JM109 | *recA1*, *endA1*, *thi*, *gyrA96*, *supE44*, *hsdR17*Δ (*lac-proAB*) /F'[*traD36*,*proAB*$^{+}$, *lacI*$^{q}$, *lacZ*Δ *M15*] | Laboratory stock |
| BS | Wild-type *Bacillus subtilis* 168 | Laboratory stock |
| BSΔ1 | Knockout genes including from *yceB* to *ycgG* | This study |
| BSΔ2 | Knockout genes including from *srfAC* to *sfpAn* | This study |
| BSΔ3 | Knockout genes including from *yclG* to *gabR* | This study |
| BSΔ5 | Knockout genes including from *dinB* to *ydhUn* | This study |
| BSΔ6 | Knockout genes including from *ydiM* to *ydjC* | This study |
| BSΔ7 | Knockout genes including from *pspA* to *yebA* | This study |
| BSΔ8 | Knockout genes including from *yefB* to *yetN* | This study |
| BSΔ9 | Knockout genes including from *sspH* to *yfiV* | This study |
| BSΔ10 | Knockout genes including from *yfhH* to *yfhP* | This study |

| | | |
|---|---|---|
| BSΔ11 | Knockout genes including from *yhfM* to *yhfP* | This study |
| BSΔ13 | Knockout genes including from *yitF* to *fakBB* | This study |
| BSΔ14 | Knockout genes including from *yitT* to *yjaV* | This study |
| BSΔ15 | Knockout genes including from *cotO* to *spoVIF* | This study |
| BSΔ16 | Knockout genes including from *manP* to *yjdJ* | This study |
| BSΔ17 | Knockout genes including from *cotT* to *yjlB* | This study |
| BSΔ18 | Knockout genes including from *uxaC* to *spoIISA* | This study |
| BSΔ19 | Knockout genes including from *ykzB* to *ykoT* | This study |
| BSΔ20 | Knockout genes including from *kinD* to *motA* | This study |
| BSΔ21 | Knockout genes including from *ykvN* to *ykvU* | This study |
| BSΔ22 | Knockout genes including from *splA* to *pbpH* | This study |
| BSΔ23 | Knockout genes including from *yktA* to *nprE* | This study |
| BSΔ24 | Knockout genes including from *bpr* to *sigG* | This study |
| BSΔ25 | Knockout genes including from *cysP* to *tcaB* | This study |
| BSΔ27 | Knockout genes including from *yoxC* to *yoaU* | This study |
| BSΔ28 | Knockout genes including from *walL* to *yozC* | This study |
| BSΔ29 | Knockout genes including from *bshBB* to *ldcB* | This study |
| BSΔ30 | Knockout genes including from *yoyE* to *sprB* | This study |
| BSΔ31 | Knockout genes including from *spsMc* to *ycgG* | This study |
| BSΔ32 | Knockout genes including from *ypwA* to *ypzG* | This study |
| BSΔ33 | Knockout genes including from *ypsA* to *sspM* | This study |
| BSΔ34 | Knockout genes including from *spoIIIAH* to *yqhV* | This study |
| BSΔ35 | Knockout genes including from *tasA* to *tapA* | This study |
| BSΔ36 | Knockout genes including from *yraJ* to *levR* | This study |
| BSΔ37 | Knockout genes including from *yrhK* to *yrhC* | This study |
| BSΔ38 | Knockout genes including from *sspI* to *frvX* | This study |
| BSΔ39 | Knockout genes including from *ytzH* to *ytcP* | This study |
| BSΔ40 | Knockout genes including from *yttA* to *ytoA* | This study |
| BSΔ41 | Knockout genes including from *yteA* to *ytaB* | This study |
| BSΔ42 | Knockout genes including from *rhaA* to *yugM* | This study |

| | | |
|---|---|---|
| BSΔ43 | Knockout genes including from *yuxO* to *comQ* | This study |
| BSΔ44 | Knockout genes including from *yueI* to *yukJ* | This study |
| BSΔ45 | Knockout genes including from *lytH* to *glxB* | This study |
| BSΔ46 | Knockout genes including from *pgoN* to *yvgT* | This study |
| BSΔ47 | Knockout genes including from *araE* to *lutP* | This study |
| BSΔ48 | Knockout genes including from *epsO* to *slrR* | This study |
| BSΔ49 | Knockout genes including from *padC* to *cotNP* | This study |
| BSΔ50 | Knockout genes including from *mdxM* to *crh* | This study |
| BSΔ51 | Knockout genes including from *nagA* to *yvzB* | This study |
| BSΔ52 | Knockout genes including from *lytD* to *ywsA* | This study |
| BSΔ53 | Knockout genes including from *ywrJ* to *ywqG* | This study |
| BSΔ54 | Knockout genes including from *mta* to *rapB* | This study |
| BSΔ55 | Knockout genes including from *sboA* to *albG* | This study |
| BSΔ56 | Knockout genes including from *bacG* to *bacA* | This study |
| BSΔ57 | Knockout genes including from *bslB* to *ywdH* | This study |
| BSΔ58 | Knockout genes including from *ywbB* to *epr* | This study |
| BSΔ59 | Knockout genes including from *ywzH* to *dltE* | This study |
| BSΔ60 | Knockout genes including from *licH* to *nnrA* | This study |
| BSΔ61 | Knockout genes including from *cimH* to *wapA* | This study |
| BSΔ62 | Knockout genes including from *yxiE* to *hutM* | This study |
| BSΔ63 | Knockout genes including from *yybP* to *yyaJ* | This study |
| BSΔ64 | Knockout genes including from *bshBB* to *ypmR* | This study |
| BSΔ65 | Knockout genes including from *walL* to *ilvA* | This study |
| BS-p566-sfGFP | Wild-type *Bacillus subtilis* 168, Harboring pHT01-P566-sfGFP | This study |
| BS-alsSD | Wild-type *Bacillus subtilis* 168, Harboring pHT01-P43-*alsS-alsD* | Laboratory stock |
| BS-CrtEBI | Wild-type *Bacillus subtilis* 168, Harboring pHT01-P43-*CrtEBI* | This study |
| BS-Δ21-1 | Knockout genes *yKvN* | This study |
| BS-Δ21-2 | Knockout genes *ykvO* | This study |
| BS-Δ21-3 | Knockout genes *yKvP* | This study |

| | | |
|---|---|---|
| BS-Δ21-4 | Knockout genes *yKzQ* | This study |
| BS-Δ21-5 | Knockout genes *ykvQ* | This study |
| BS-Δ21-6 | Knockout genes *ykzR* | This study |
| BS-Δ21-7 | Knockout genes *ykvR* | This study |
| BS-Δ21-8 | Knockout genes *ykvS* | This study |
| BS-Δ21-9 | Knockout genes *ykzS* | This study |
| BS-Δ21-10 | Knockout genes *ykvT* | This study |
| BS-Δ21-11 | Knockout genes *ykvU* | This study |

**Table S4. The plasmids used in this study.**

| Plasmids | Description | Source |
|---|---|---|
| p8999-cas9-2N20-HA-Δ1 | A genome-scale knockout plasmid harboring two sgRNAs and HA, targeting the Δ1 regions | This study |
| p8999-cas9-2N20-HA-Δ2 | A genome-scale knockout plasmid harboring two sgRNAs and HA, targeting the Δ2 regions | This study |
| p8999-cas9-2N20-HA-Δ3 | A genome-scale knockout plasmid harboring two sgRNAs and HA, targeting the Δ3 regions | This study |
| p8999-cas9-2N20-HA-Δ5 | A genome-scale knockout plasmid harboring two sgRNAs and HA, targeting the Δ5 regions | This study |
| p8999-cas9-2N20-HA-Δ6 | A genome-scale knockout plasmid harboring two sgRNAs and HA, targeting the Δ6 regions | This study |
| p8999-cas9-2N20-HA-Δ7 | A genome-scale knockout plasmid harboring two sgRNAs and HA, targeting the Δ7 regions | This study |
| p8999-cas9-2N20-HA-Δ8 | A genome-scale knockout plasmid harboring two sgRNAs and HA, targeting the Δ8 regions | This study |
| p8999-cas9-2N20-HA-Δ9 | A genome-scale knockout plasmid harboring two sgRNAs and HA, targeting the Δ9 regions | This study |
| p8999-cas9-2N20-HA-Δ10 | A genome-scale knockout plasmid harboring two sgRNAs and HA, targeting the Δ10 regions | This study |
| p8999-cas9-2N20-HA-Δ11 | A genome-scale knockout plasmid harboring two sgRNAs and HA, targeting the Δ11 regions | This study |

| | | |
|---|---|---|
| p8999-cas9-2N20-HA-Δ13 | A genome-scale knockout plasmid harboring two sgRNAs and HA, targeting the Δ13 regions | This study |
| p8999-cas9-2N20-HA-Δ14 | A genome-scale knockout plasmid harboring two sgRNAs and HA, targeting the Δ14 regions | This study |
| p8999-cas9-2N20-HA-Δ15 | A genome-scale knockout plasmid harboring two sgRNAs and HA, targeting the Δ15 regions | This study |
| p8999-cas9-2N20-HA-Δ16 | A genome-scale knockout plasmid harboring two sgRNAs and HA, targeting the Δ16 regions | This study |
| p8999-cas9-2N20-HA-Δ17 | A genome-scale knockout plasmid harboring two sgRNAs and HA, targeting the Δ17 regions | This study |
| p8999-cas9-2N20-HA-Δ18 | A genome-scale knockout plasmid harboring two sgRNAs and HA, targeting the Δ18 regions | This study |
| p8999-cas9-2N20-HA-Δ19 | A genome-scale knockout plasmid harboring two sgRNAs and HA, targeting the Δ19 regions | This study |
| p8999-cas9-2N20-HA-Δ20 | A genome-scale knockout plasmid harboring two sgRNAs and HA, targeting the Δ20 regions | This study |
| p8999-cas9-2N20-HA-Δ21 | A genome-scale knockout plasmid harboring two sgRNAs and HA, targeting the Δ21 regions | This study |
| p8999-cas9-2N20-HA-Δ22 | A genome-scale knockout plasmid harboring two sgRNAs and HA, targeting the Δ22 regions | This study |
| p8999-cas9-2N20-HA-Δ23 | A genome-scale knockout plasmid harboring two sgRNAs and HA, targeting the Δ23 regions | This study |
| p8999-cas9-2N20-HA-Δ24 | A genome-scale knockout plasmid harboring two sgRNAs and HA, targeting the Δ24 regions | This study |
| p8999-cas9-2N20-HA-Δ25 | A genome-scale knockout plasmid harboring two sgRNAs and HA, targeting the Δ25 regions | This study |
| p8999-cas9-2N20-HA-Δ27 | A genome-scale knockout plasmid harboring two sgRNAs and HA, targeting the Δ27 regions | This study |
| p8999-cas9-2N20-HA-Δ28 | A genome-scale knockout plasmid harboring two sgRNAs and HA, targeting the Δ28 regions | This study |
| p8999-cas9-2N20-HA-Δ29 | A genome-scale knockout plasmid harboring two sgRNAs and HA, targeting the Δ29 regions | This study |

| | | |
|---|---|---|
| p8999-cas9-2N20-HA-Δ30 | A genome-scale knockout plasmid harboring two sgRNAs and HA, targeting the Δ30 regions | This study |
| p8999-cas9-2N20-HA-Δ31 | A genome-scale knockout plasmid harboring two sgRNAs and HA, targeting the Δ31 regions | This study |
| p8999-cas9-2N20-HA-Δ32 | A genome-scale knockout plasmid harboring two sgRNAs and HA, targeting the Δ32 regions | This study |
| p8999-cas9-2N20-HA-Δ33 | A genome-scale knockout plasmid harboring two sgRNAs and HA, targeting the Δ33 regions | This study |
| p8999-cas9-2N20-HA-Δ34 | A genome-scale knockout plasmid harboring two sgRNAs and HA, targeting the Δ34 regions | This study |
| p8999-cas9-2N20-HA-Δ35 | A genome-scale knockout plasmid harboring two sgRNAs and HA, targeting the Δ35 regions | This study |
| p8999-cas9-2N20-HA-Δ36 | A genome-scale knockout plasmid harboring two sgRNAs and HA, targeting the Δ36 regions | This study |
| p8999-cas9-2N20-HA-Δ37 | A genome-scale knockout plasmid harboring two sgRNAs and HA, targeting the Δ37 regions | This study |
| p8999-cas9-2N20-HA-Δ38 | A genome-scale knockout plasmid harboring two sgRNAs and HA, targeting the Δ38 regions | This study |
| p8999-cas9-2N20-HA-Δ39 | A genome-scale knockout plasmid harboring two sgRNAs and HA, targeting the Δ39 regions | This study |
| p8999-cas9-2N20-HA-Δ40 | A genome-scale knockout plasmid harboring two sgRNAs and HA, targeting the Δ40 regions | This study |
| p8999-cas9-2N20-HA-Δ41 | A genome-scale knockout plasmid harboring two sgRNAs and HA, targeting the Δ41 regions | This study |
| p8999-cas9-2N20-HA-Δ42 | A genome-scale knockout plasmid harboring two sgRNAs and HA, targeting the Δ42 regions | This study |
| p8999-cas9-2N20-HA-Δ43 | A genome-scale knockout plasmid harboring two sgRNAs and HA, targeting the Δ43 regions | This study |
| p8999-cas9-2N20-HA-Δ44 | A genome-scale knockout plasmid harboring two sgRNAs and HA, targeting the Δ44 regions | This study |
| p8999-cas9-2N20-HA-Δ45 | A genome-scale knockout plasmid harboring two sgRNAs and HA, targeting the Δ45 regions | This study |

| | | |
|---|---|---|
| p8999-cas9-2N20-HA-Δ46 | A genome-scale knockout plasmid harboring two sgRNAs and HA, targeting the Δ46 regions | This study |
| p8999-cas9-2N20-HA-Δ47 | A genome-scale knockout plasmid harboring two sgRNAs and HA, targeting the Δ47 regions | This study |
| p8999-cas9-2N20-HA-Δ48 | A genome-scale knockout plasmid harboring two sgRNAs and HA, targeting the Δ48 regions | This study |
| p8999-cas9-2N20-HA-Δ49 | A genome-scale knockout plasmid harboring two sgRNAs and HA, targeting the Δ49 regions | This study |
| p8999-cas9-2N20-HA-Δ50 | A genome-scale knockout plasmid harboring two sgRNAs and HA, targeting the Δ50 regions | This study |
| p8999-cas9-2N20-HA-Δ51 | A genome-scale knockout plasmid harboring two sgRNAs and HA, targeting the Δ51 regions | This study |
| p8999-cas9-2N20-HA-Δ52 | A genome-scale knockout plasmid harboring two sgRNAs and HA, targeting the Δ52 regions | This study |
| p8999-cas9-2N20-HA-Δ53 | A genome-scale knockout plasmid harboring two sgRNAs and HA, targeting the Δ53 regions | This study |
| p8999-cas9-2N20-HA-Δ54 | A genome-scale knockout plasmid harboring two sgRNAs and HA, targeting the Δ54 regions | This study |
| p8999-cas9-2N20-HA-Δ55 | A genome-scale knockout plasmid harboring two sgRNAs and HA, targeting the Δ55 regions | This study |
| p8999-cas9-2N20-HA-Δ56 | A genome-scale knockout plasmid harboring two sgRNAs and HA, targeting the Δ56 regions | This study |
| p8999-cas9-2N20-HA-Δ57 | A genome-scale knockout plasmid harboring two sgRNAs and HA, targeting the Δ57 regions | This study |
| p8999-cas9-2N20-HA-Δ58 | A genome-scale knockout plasmid harboring two sgRNAs and HA, targeting the Δ58 regions | This study |
| p8999-cas9-2N20-HA-Δ59 | A genome-scale knockout plasmid harboring two sgRNAs and HA, targeting the Δ59 regions | This study |
| p8999-cas9-2N20-HA-Δ60 | A genome-scale knockout plasmid harboring two sgRNAs and HA, targeting the Δ60 regions | This study |
| p8999-cas9-2N20-HA-Δ61 | A genome-scale knockout plasmid harboring two sgRNAs and HA, targeting the Δ61 regions | This study |

| | | |
|---|---|---|
| p8999-cas9-2N20-HA-Δ62 | A genome-scale knockout plasmid harboring two sgRNAs and HA, targeting the Δ62 regions | This study |
| p8999-cas9-2N20-HA-Δ63 | A genome-scale knockout plasmid harboring two sgRNAs and HA, targeting the Δ63 regions | This study |
| p8999-cas9-2N20-HA-Δ64 | A genome-scale knockout plasmid harboring two sgRNAs and HA, targeting the Δ64 regions | This study |
| p8999-cas9-2N20-HA-Δ65 | A genome-scale knockout plasmid harboring two sgRNAs and HA, targeting the Δ65 regions | This study |

**Table S5. The Primers used in this study.**

| Primers | Sequences (5'-3') |
|---|---|
| Δ1-N20-1 | ATGCGGGCGATCATGATTTCGTTTTAGAGCTAGAAATAGCAAGTTAAAATAAGGC |
| Δ1-N20-2 | GAAATCATGATCGCCCGCATCGTAGGTACATTTTACTCAATTCTCTAATCA |
| Δ1-N20-3 | TTTCATTCTGTGCGCTACACGTTTTAGAGCTAGAAATAGCAAGTTAAAATAAGGC |
| Δ1-N20-4 | GTGTAGCGCACAGAATGAAACGTAGGTACATTTTACTCAATTCTCTAATCA |
| Δ1-HA-1 | TTCTCCCCCATTACATCACTTCTACCTGCTGACGTTTGAT |
| Δ1-HA-2 | AGCTGCAGGATGTCTCTTAT |
| Δ1-HA-3 | ATAAGAGACATCCTGCAGCTATGGAAGCAGTATTTCCTCC |
| Δ1-HA-4 | GGTGACTGAAGTATACCGAATCAATCTACGCAGTACTTGG |
| Δ1-Test-1 | TTAATGGCAGCTGGTTTGTCC |
| Δ1-Test-2 | TGTACTGGACATCAGAAATGCTG |
| Δ1-Test-3 | TCCTCCTGATCGATTATTTAGGAT |
| Δ1-Test-4 | TTGAGTAATACGGTCCAGGGT |
| Δ2-N20-1 | CGTCGTACAGGATCTATTTAGTTTTAGAGCTAGAAATAGCAAGTTAAAATAAGG |

| | |
|---|---|
| Δ2-N20-2 | TAAATAGATCCTGTACGACGCGTAGGTACATTTTACTCAATTCTCTAATC |
| Δ2-N20-3 | GCGTTTGATTCACAGCCGATGTTTTAGAGCTAGAAATAGCAAGTTAAAATAAGG |
| Δ2-N20-4 | ATCGGCTGTGAATCAAACGCCGTAGGTACATTTTACTCAATTCTCTAATC |
| Δ2-HA-1 | AGTGGTTCTATCGAGAGTCGTTACAGCAGGCTGAATACTGG |
| Δ2-HA-2 | ATATCACCTGAATCGGCGATAGGTAATACATATCTT |
| Δ2-HA-3 | TATCGCCGATTCAGGTGATATGAAAGACATGAACC |
| Δ2-HA-4 | CGGCAAATCCAATCGCATAGTGACAACGAATGCAAGGGTT |
| Δ2-Test-1 | CTACATGCAGGCTGAGAAAGAA |
| Δ2-Test-2 | TTGTGATTTTCAGCGTGATTGAA |
| Δ2-Test-3 | TATCATTTCTGCAATCCAGCCG |
| Δ2-Test-4 | TTCTGACACTGAAACGGTCGA |
| Δ3-N20-1 | TTCCTATCGTAATCATACAGGTTTTAGAGCTAGAAATAGCAAGTTAAAATAAGGC |
| Δ3-N20-2 | CTGTATGATTACGATAGGAACGTAGGTACATTTTACTCAATTCTCTAATCACGG |
| Δ3-N20-3 | GGCATCGTCACTTCAGCCATGTTTTAGAGCTAGAAATAGCAAGTTAAAATAAGGC |
| Δ3-N20-4 | ATGGCTGAAGTGACGATGCCCGTAGGTACATTTTACTCAATTCTCTAATCACGG |
| Δ3-HA-1 | GCTTCTCCCCCATTACATCACTTGCAGGAAACCAAACTATATACTGGA |
| Δ3-HA-2 | ATTCCTTACAAGACGCGTTGATTGACGTACTC |
| Δ3-HA-3 | GTCAATCAACGCGTCTTGTAAGGAATACAGCCAGACCG |
| Δ3-HA-4 | AGGAGGTGACTGAAGTATACCGAAATCAGCTACGATGACAGGGTTT |
| Δ3-Test-1 | ATCACAGGAATCAAAATACCGAG |
| Δ3-Test-2 | ATCCAGATCGTACAGCTCG |

| | |
|---|---|
| Δ3-Test-3 | TGTTAAAAGCTGAAATCCAGCAG |
| Δ3-Test-4 | TGTGATATTCCGAAGCTTTATCATG |
| Δ4-N20-1 | TTCGCTTTGCTTGTCGCAATGTTTTAGAGCTAGAAATAGCAAGTTAAAATAAGG |
| Δ4-N20-2 | ATTGCGACAAGCAAAGCGAACGTAGGTACATTTTACTCAATTCTCTAATC |
| Δ4-N20-3 | GCTGACAGCGATCATCCATAGTTTTAGAGCTAGAAATAGCAAGTTAAAATAAGG |
| Δ4-N20-4 | TATGGATGATCGCTGTCAGCCGTAGGTACATTTTACTCAATTCTCTAATC |
| Δ4-HA-1 | AGTGGTTCTATCGAGAGTCGTTGAAGAATAATGCGGCAGC |
| Δ4-HA-2 | GGCTCTACATGATTATATGGGACCTTATCACGAATCC |
| Δ4-HA-3 | GTCCCATATAATCATGTAGAGCCTGACGACAT |
| Δ4-HA-4 | CGGCAAATCCAATCGCATAGTATCCATGCCAGCTGAGGAAG |
| Δ4-Test-1 | AGACATAGGGCTGACAAGATG |
| Δ4-Test-2 | TTCGGCATTTGCTGATTCCG |
| Δ4-Test-3 | CAGAGGAATTGACTGGTGTCAG |
| Δ4-Test-4 | AGAATGACGAGATCTGGCTG |
| Δ5-N20-1 | TCGGTTGAATATTTGTTGATGTTTTAGAGCTAGAAATAGCAAGTTAAAATAAGG |
| Δ5-N20-2 | ATCAACAAATATTCAACCGACGTAGGTACATTTTACTCAATTCTCTAATC |
| Δ5-N20-3 | GCATTCATATCGCCTAACCCGTTTTAGAGCTAGAAATAGCAAGTTAAAATAAGG |
| Δ5-N20-4 | GGGTTAGGCGATATGAATGCCGTAGGTACATTTTACTCAATTCTCTAATC |
| Δ5-HA-1 | TTCTCCCCCATTACATCACTAAGGATAGATGCTCATAATCACGAG |
| Δ5-HA-2 | TTGGAGCGCAAAGTAACTGTTTCCGAACTGCTTCC |
| Δ5-HA-3 | ACAGTTACTTTGCGCTCCAATAATATAATGA |

| | |
|---|---|
| Δ5-HA-4 | GAGGTGACTGAAGTATACCGAAACTTGCATGTATTAGGCACG |
| Δ5-Test-1 | TACTGAATACAGCGCTGTTGC |
| Δ5-Test-2 | TCCATCTGTAAGTGGTAGCCG |
| Δ5-Test-3 | CAAAGACGTCACTTGTCGTG |
| Δ5-Test-4 | TGCTTGTATGAAGTAACGGGATC |
| Δ6-N20-1 | ATAGGTATCATAGTCGATGCGTTTTAGAGCTAGAAATAGCAAGTTAAAATAAGG |
| Δ6-N20-2 | GCATCGACTATGATACCTATCGTAGGTACATTTTACTCAATTCTCTAATC |
| Δ6-N20-3 | ATTTACGGGCGTTCATTCACGTTTTAGAGCTAGAAATAGCAAGTTAAAATAAGG |
| Δ6-N20-4 | GTGAATGAACGCCCGTAAATCGTAGGTACATTTTACTCAATTCTCTAATC |
| Δ6-HA-1 | CTTCTCCCCCATTACATCACTTTAGGTCGGTCTAGGCAAGTAC |
| Δ6-HA-2 | CGGAATGACCACATTGCAATGGTGTATACTTTGTGG |
| Δ6-HA-3 | ATTGCAATGTGGTCATTCCGGTATTGTT |
| Δ6-HA-4 | GAGGTGACTGAAGTATACCGAAACTCTATCTATTGGAGCTACCTGC |
| Δ6-Test-1 | ACGATATCAACAGATCCTGTTG |
| Δ6-Test-2 | ACACAATACTTAGCGCACCAA |
| Δ6-Test-3 | TAATCCTCAACAGTGGTAACCTTG |
| Δ6-Test-4 | ATGAACTACAAGCATGTAGCGG |
| Δ7-N20-1 | GACAAATTGGTATCTGACATGTTTTAGAGCTAGAAATAGCAAGTTAAAATAAGG |
| Δ7-N20-2 | ATGTCAGATACCAATTTGTCCGTAGGTACATTTTACTCAATTCTCTAATC |
| Δ7-N20-3 | GTTACATTTGAGCCGACTAAGTTTTAGAGCTAGAAATAGCAAGTTAAAATAAGG |
| Δ7-N20-4 | TTAGTCGGCTCAAATGTAACCGTAGGTACATTTTACTCAATTCTCTAATC |

| | |
|---|---|
| Δ7-HA-1 | CTTCTCCCCCATTACATCACTTGTACATGTCGATACCTCTATGCTG |
| Δ7-HA-2 | GTCTTCTATGTCGTACCTTAGCCAAATCGCTATTCA |
| Δ7-HA-3 | GGCTAAGGTACGACATAGAAGACATGTCAAAGC |
| Δ7-HA-4 | GGAGGTGACTGAAGTATACCGAATACGGAGAAGACCATGGTCTACA |
| Δ7-Test-1 | TTGTACCGATGTTGACGTTGAT |
| Δ7-Test-2 | ACACGTAAATGAATTCATTACCGAT |
| Δ7-Test-3 | TTACTTGGTATAATGGAGTCGGGA |
| Δ7-Test-4 | TTCCTAAGCGGAAATAGATGGT |
| Δ8-N20-1 | AAGAGATGTACTTCAGCTTGGTTTTAGAGCTAGAAATAGCAAGTTAAAATAAGG |
| Δ8-N20-2 | CAAGCTGAAGTACATCTCTTCGTAGGTACATTTTACTCAATTCTCTAATC |
| Δ8-N20-3 | GAGAAGACGCGTCAGAGATGGTTTTAGAGCTAGAAATAGCAAGTTAAAATAAGG |
| Δ8-N20-4 | CATCTCTGACGCGTCTTCTCCGTAGGTACATTTTACTCAATTCTCTAATC |
| Δ8-HA-1 | CTTCTCCCCCATTACATCACTATGACATCATCGACATGAGCG |
| Δ8-HA-2 | GTTCAAGCACTTAGAGCTTAGATAAATTCATGAAGGAAGG |
| Δ8-HA-3 | CTAAGCTCTAAGTGCTTGAACTGCTTGTTC |
| Δ8-HA-4 | GGAGGTGACTGAAGTATACCGAAAACGTGATGATCTGAAAGCG |
| Δ8-Test-1 | AAGACGTCTTAGAACGCATCG |
| Δ8-Test-2 | TCATCCGTATGTATGTAAGCTCCA |
| Δ8-Test-3 | TATCGTCAAAGGCAAATCAGCG |
| Δ8-Test-4 | AGCACATACGTAAACGTCACC |
| Δ9-N20-1 | CCGATCCAGTCATACAGATCGTTTTAGAGCTAGAAATAGCAAGTTAAAATAAGG |

| | |
|---|---|
| Δ9-N20-2 | GATCTGTATGACTGGATCGGCGTAGGTACATTTTACTCAATTCTCTAATC |
| Δ9-N20-3 | TCAACTGAACAACTACAGAGGTTTTAGAGCTAGAAATAGCAAGTTAAAATAAGG |
| Δ9-N20-4 | CTCTGTAGTTGTTCAGTTGACGTAGGTACATTTTACTCAATTCTCTAATC |
| Δ9-HA-1 | CTTCTCCCCCATTACATCACTTCTGTTTGTCGGGTGAAACG |
| Δ9-HA-2 | ACTGGATATGATTCGTCAAGCGGATAAATTCTTGC |
| Δ9-HA-3 | GCTTGACGAATCATATCCAGTTTCATCCACTCGG |
| Δ9-HA-4 | GAGGTGACTGAAGTATACCGAAACATACCTGATATCTGCATGAATGC |
| Δ9-Test-1 | TCAATCGAAAATGCTGTCACCAT |
| Δ9-Test-2 | TCTTGATGATAGCCAAGCTGC |
| Δ9-Test-3 | AACTGATGATCGTAATTGGCCA |
| Δ9-Test-4 | ACATGCATAGGAAAGATTTCCG |
| Δ10-N20-1 | TTTCATTGCCTTCAGTCGTCGTTTTAGAGCTAGAAATAGCAAGTTAAAATAAGGC |
| Δ10-N20-2 | CGACGACTGAAGGCAATGAAACGTAGGTACATTTTACTCAATTCTCTAATCACGG |
| Δ10-N20-3 | TGTCACTGGATCCTAAGGAAGTTTTAGAGCTAGAAATAGCAAGTTAAAATAAGGC |
| Δ10-N20-4 | ACTTCCTTAGGATCCAGTGACACGTAGGTACATTTTACTCAATTCTCTAATCACGG |
| Δ10-HA-1 | AGTGGTTCTATCGAGAGTCGTATGAGGAAGTTCCTGCTCCG |
| Δ10-HA-2 | GGATCAGATATGCGGTGTCATTTGACTGTAT |
| Δ10-HA-3 | TGACACCGCATATCTGATCCGACAACAGGATC |
| Δ10-HA-4 | CGGCAAATCCAATCGCATAGTACACTTGTCCGTCTTCATCAG |
| Δ10-Test-1 | TTCACGCGTCGCATTGATAC |
| Δ10-Test-2 | AACGCAAATTGTTCGAGCTC |

| | |
|---|---|
| Δ10-Test-3 | AGCAATTGTCATCGGTCATGA |
| Δ10-Test-4 | ATCCTTGCTTCATTAAGGCAC |
| Δ11-N20-1 | ACGGAATATCGTGGGAAATCGTTTTAGAGCTAGAAATAGCAAGTTAAAATAAGGC |
| Δ11-N20-2 | GATTTCCCACGATATTCCGTCGTAGGTACATTTTACTCAATTCTCTAATCACGG |
| Δ11-N20-3 | GGATATCATATAAAGGGGGAGGTTTTAGAGCTAGAAATAGCAAGTTAAAATAAGGC |
| Δ11-N20-4 | ACCTCCCCCTTTATATGATATCCGTAGGTACATTTTACTCAATTCTCTAATCACGG |
| Δ11-HA-1 | TAATCGAATTGGTGCAGGAAAGGAC |
| Δ11-HA-2 | ATGCAAGAAGTGTGCCAGTCT |
| Δ11-HA-3 | TGATTCAGCAACATCAGGGCTG |
| Δ11-HA-4 | ATTTGTATCCTTCTAACGGGGTG |
| Δ11-Test-1 | AGAACCTGTACCTGATGATACC |
| Δ11-Test-2 | ATCCACGTGGTATACATGCC |
| Δ11-Test-3 | TATTGTGCAGCTGCTTGTACG |
| Δ11-Test-4 | ATTGACTCTTCTCATCCTGACTT |
| Δ12-N20-1 | AGTGTCTCCTTTTTCTGGGTGTTTTAGAGCTAGAAATAGCAAGTTAAAATAAGG |
| Δ12-N20-2 | ACCCAGAAAAAGGAGACACTCGTAGGTACATTTTACTCAATTCTCTAATC |
| Δ12-N20-3 | CATAGAAGCGAATGGATATTGTTTTAGAGCTAGAAATAGCAAGTTAAAATAAGG |
| Δ12-N20-4 | AATATCCATTCGCTTCTATGCGTAGGTACATTTTACTCAATTCTCTAATC |
| Δ12-HA-1 | AGTGGTTCTATCGAGAGTCGCAGGAAGCTGATGATCATATCGA |
| Δ12-HA-2 | GCATTACCCGAAACTATGGCGGTCATTCTAGTAG |
| Δ12-HA-3 | GCCATAGTTTCGGGTAATGCCTCTTTAAACAG |
| Δ12-HA-4 | CGGCAAATCCAATCGCATAGTATAGTCAGCCTCGTTCAAATGATG |

| | |
|---|---|
| Δ12-Test-1 | AAGGCTCCCTGACATGAAG |
| Δ12-Test-2 | TGGCCATGCTGTCAGGATATGT |
| Δ12-Test-3 | TCCCTACTGTCTGTAACATACG |
| Δ12-Test-4 | TCCGCATACAATCGACGA |
| Δ13-N20-1 | TGCGTCCAGAATCATCGTAAGTTTTAGAGCTAGAAATAGCAAGTTAAAATAAGG |
| Δ13-N20-2 | TTACGATGATTCTGGACGCACGTAGGTACATTTTACTCAATTCTCTAATCAC |
| Δ13-N20-3 | ATGATTTAACATATCTCGCGGTTTTAGAGCTAGAAATAGCAAGTTAAAATAAGG |
| Δ13-N20-4 | CGCGAGATATGTTAAATCATCGTAGGTACATTTTACTCAATTCTCTAATCAC |
| Δ13-HA-1 | AGTGGTTCTATCGAGAGTCGTTCTGTAAAATAGCGGCTGCG |
| Δ13-HA-2 | CATGCGGATCAGTTATTTCGGCGTCAGAGCA |
| Δ13-HA-3 | CGAAATAACTGATCCGCATGAATTGTGACTG |
| Δ13-HA-4 | CGGCAAATCCAATCGCATAGCATACCTCGTGATGACGATAATCA |
| Δ13-Test-1 | TGTTGACGACATCATTTCCTCG |
| Δ13-Test-2 | ACTTCCACCTGCTTTCGTCA |
| Δ13-Test-3 | TCGCTTAGCGCTTCAATCAT |
| Δ13-Test-4 | AAATCCAGCAAGCTCGAACA |
| Δ14-N20-1 | GCAGATTCTCAGATTCAAGAAGTTTTAGAGCTAGAAATAGCAAGTTAAAATAAGGC |
| Δ14-N20-2 | CTTCTTGAATCTGAGAATCTGCGTAGGTACATTTTACTCAATTCTCTAATCACGG |
| Δ14-N20-3 | GATTTAATCGCTGCGACGAGGTTTTAGAGCTAGAAATAGCAAGTTAAAATAAGGC |
| Δ14-N20-4 | CTCGTCGCAGCGATTAAATCCGTAGGTACATTTTACTCAATTCTCTAATCACGG |
| Δ14-HA-1 | GCTTCTCCCCCATTACATCACTATCCATATATGCACCGCGATACAA |

| | |
|---|---|
| Δ14-HA-2 | TATCATTCATGATGACGGAACATCATAATTGAATTCCTGAGCGAT |
| Δ14-HA-3 | TATGATGTTCCGTCATCATGAATGATAATCGTCA |
| Δ14-HA-4 | AGGAGGTGACTGAAGTATACCGAAACAGCCTCAATCGTATCCATCA |
| Δ14-Test-1 | TCGAAGCAATCAGCTTCAGTC |
| Δ14-Test-2 | TGTTCAGGTCGTAGTAATGATCGC |
| Δ14-Test-3 | ATTCTTTCGCATCATCCTCATG |
| Δ14-Test-4 | TACTGACCTTTTGGCCTCACTC |
| Δ15-N20-1 | GCATTGTTCTGAGAGAATGAAGTTTTAGAGCTAGAAATAGCAAGTTAAAATAAGGC |
| Δ15-N20-2 | CTTCATTCTCTCAGAACAATGCGTAGGTACATTTTACTCAATTCTCTAATCACGG |
| Δ15-N20-3 | GAGTGTGCGTCCGATCGATTGTTTTAGAGCTAGAAATAGCAAGTTAAAATAAGGC |
| Δ15-N20-4 | AATCGATCGGACGCACACTCCGTAGGTACATTTTACTCAATTCTCTAATCACGG |
| Δ15-HA-1 | TTCTCCCCCATTACATCACTTCACTTGAAGGCCGTAACATTG |
| Δ15-HA-2 | AATCACACTTTGATGACAATCTCCTGCATGG |
| Δ15-HA-3 | GATTGTCATCAAAGTGTGATTAACCGCGTAGC |
| Δ15-HA-4 | GAGGTGACTGAAGTATACCGAAAGGCCGTGCTATATATGAAAGG |
| Δ15-Test-1 | TTGCGTTTGCTACGGTATGC |
| Δ15-Test-2 | AACTGCGAACTACAGGTCTAC |
| Δ15-Test-3 | ATTAAGTGCTGTTACAACTGATGT |
| Δ15-Test-4 | AGAAGAGCTTGCAAAGCTTC |
| Δ16-N20-1 | CCAATTGCCATGGCACTGACGTTTTAGAGCTAGAAATAGCAAGTTAAAATAAGGC |
| Δ16-N20-2 | GTCAGTGCCATGGCAATTGGCGTAGGTACATTTTACTCAATTCTCTAATCACGG |

| | |
|---|---|
| Δ16-N20-3 | CCATGGGAGTGAATATGCTGGTTTTAGAGCTAGAAATAGCAAGTTAAAATAAGGC |
| Δ16-N20-4 | CAGCATATTCACTCCCATGGCGTAGGTACATTTTACTCAATTCTCTAATCACGG |
| Δ16-HA-1 | GCTTCTCCCCCATTACATCACTAAGACGCGTACACAGGAATCT |
| Δ16-HA-2 | AAGGAAGCAATTATGTGTGTGACGTCCTATACAACAG |
| Δ16-HA-3 | CGTCACACACATAATTGCTTCCTTTACGTGGCAG |
| Δ16-HA-4 | AGGAGGTGACTGAAGTATACCGAAAGAATGATGTTCGGCGATGCA |
| Δ16-Test-1 | ACGGAAATATCACCAAATCGC |
| Δ16-Test-2 | CGGTCACGCTATTATTTCCTTTC |
| Δ16-Test-3 | TGATTATGACCGAAAAGATGCAGA |
| Δ16-Test-4 | ATTGTCCACAACTTCAACAACG |
| Δ17-N20-1 | TTACCATGAGAAGCAGCAGGGTTTTAGAGCTAGAAATAGCAAGTTAAAATAAGGC |
| Δ17-N20-2 | CCTGCTGCTTCTCATGGTAACGTAGGTACATTTTACTCAATTCTCTAATCACGG |
| Δ17-N20-3 | GGTTAAACTGAAGTTCACGGAGTTTTAGAGCTAGAAATAGCAAGTTAAAATAAGGC |
| Δ17-N20-4 | CTCCGTGAACTTCAGTTTAACCGTAGGTACATTTTACTCAATTCTCTAATCACGG |
| Δ17-HA-1 | TTCTCCCCCATTACATCACTTGACTACTGGCAAGAGCGGAA |
| Δ17-HA-2 | TATCGTACTGAACAGACAGCCTTATTATTATCCGCG |
| Δ17-HA-3 | GCTGTCTGTTCAGTACGATATGAAGACGGG |
| Δ17-HA-4 | GAGGTGACTGAAGTATACCGAAAGCGCCTAGAATGACAATATGT |
| Δ17-Test-1 | ATCCATATGCCTCCGAGAAC |
| Δ17-Test-2 | TTGTTTACCACTGTCACGCG |
| Δ17-Test-3 | AGTATATCTGCCAATGAACGATTC |
| Δ17-Test-4 | AGCTCCTTTGTATCTTGTCCC |

| | |
|---|---|
| Δ18-N20-1 | CTCACGGCTTAGAAGTTGAAGTTTTAGAGCTAGAAATAGCAAGTTAAAATAAGGC |
| Δ18-N20-2 | CTTCAACTTCTAAGCCGTGAGCGTAGGTACATTTTACTCAATTCTCTAATCACGG |
| Δ18-N20-3 | AGCGACTCTCTCGCTTATACGTTTTAGAGCTAGAAATAGCAAGTTAAAATAAGGC |
| Δ18-N20-4 | GTATAAGCGAGAGAGTCGCTCGTAGGTACATTTTACTCAATTCTCTAATCACGG |
| Δ18-HA-1 | TTCTCCCCCATTACATCACTAGTCTTCCAACATGTTGAAGATC |
| Δ18-HA-2 | ACATATGAGCCTCGCATAATTGTGATAGAGGCTG |
| Δ18-HA-3 | ATTATGCGAGGCTCATATGTCCAGTATACCAT |
| Δ18-HA-4 | GAGGTGACTGAAGTATACCGAAACATCGTAGATCCTTATACGCTG |
| Δ18-Test-1 | TTACGATGCGCTTGTTGTAGG |
| Δ18-Test-2 | ATTGCGACTTCGGTATCAACA |
| Δ18-Test-3 | ATCGGACAATACTGTACACGG |
| Δ18-Test-4 | TGTTCATAATCGGCAACATACTCT |
| Δ19-N20-1 | AAATGACGATTTCTGAGCTTGTTTTAGAGCTAGAAATAGCAAGTTAAAATAAGG |
| Δ19-N20-2 | AAGCTCAGAAATCGTCATTTCGTAGGTACATTTTACTCAATTCTCTAATC |
| Δ19-N20-3 | ATGCTATCCTTCGCTTTTAAGTTTTAGAGCTAGAAATAGCAAGTTAAAATAAGG |
| Δ19-N20-4 | TTAAAAGCGAAGGATAGCATCGTAGGTACATTTTACTCAATTCTCTAATC |
| Δ19-HA-1 | CTTCTCCCCCATTACATCACTTTATCGGAGCGGTTTGTTCTG |
| Δ19-HA-2 | CTCCGATGATTCCAATTTACGAAGATTTTCGGTATTGTCCG |
| Δ19-HA-3 | TCGTAAATTGGAATCATCGGAGAGTATATTGG |

| | |
|---|---|
| Δ19-HA-4 | GGAGGTGACTGAAGTATACCGAAAGAGAGGTATCGTTTATCGGTTT |
| Δ19-Test-1 | TCTCTTGCGGTTGTCATCAGAG |
| Δ19-Test-2 | AGATTATGGAACAGCACGGAC |
| Δ19-Test-3 | AAGTGTCTCATGTGCCGATT |
| Δ19-Test-4 | ATCAGCCTGTGGACATCATT |
| Δ20-N20-1 | CTCCTTCAGGCATTGACTCGGTTTTAGAGCTAGAAATAGCAAGTTAAAATAAGG |
| Δ20-N20-2 | CGAGTCAATGCCTGAAGGAGCGTAGGTACATTTTACTCAATTCTCTAATC |
| Δ20-N20-3 | TGAGCCCATTCAGAGAACATGTTTTAGAGCTAGAAATAGCAAGTTAAAATAAGG |
| Δ20-N20-4 | ATGTTCTCTGAATGGGCTCACGTAGGTACATTTTACTCAATTCTCTAATC |
| Δ20-HA-1 | CTTCTCCCCCATTACATCACTTGCACAATAACCCAATCAAACTTG |
| Δ20-HA-2 | TCAGTTACGGAGGCTCTATTCATATTGAAAG |
| Δ20-HA-3 | AATAGAGCCTCCGTAACTGACGCCTTTCAGAACC |
| Δ20-HA-4 | GGAGGTGACTGAAGTATACCGAATACAGCATTGTGTTGCTGGATG |
| Δ20-Test-1 | AAGTAGTGATGTGCTGGATACAT |
| Δ20-Test-2 | AGACGTTCAGCATATGTTCCTG |
| Δ20-Test-3 | TGCATCGCTCCAACATACAC |
| Δ20-Test-4 | AAGCCTTGTGACATATCAGGC |
| Δ21-N20-1 | TGATGGAAATATCGGCTGTAGTTTTAGAGCTAGAAATAGCAAGTTAAAATAAGGC |
| Δ21-N20-2 | TACAGCCGATATTTCCATCACGTAGGTACATTTTACTCAATTCTCTAATCACGG |
| Δ21-N20-3 | ACGTGAGCGCGTTCTGCCAAGTTTTAGAGCTAGAAATAGCAAGTTAAAATAAGGC |
| Δ21-N20-4 | TTGGCAGAACGCGCTCACGTCGTAGGTACATTTTACTCAATTCTCTAATCACGG |

| | |
|---|---|
| Δ21-HA-1 | TTCTCCCCCATTACATCACTTAGAAGGTGTAACATTGCTAGGC |
| Δ21-HA-2 | ATGCTTTGCAGATAGGCTATTATGCGGAGCAAGA |
| Δ21-HA-3 | AATAGCCTATCTGCAAAGCATTGAAGGTATC |
| Δ21-HA-4 | GAGGTGACTGAAGTATACCGAATGATACAAGTGCTGCAATAAGCT |
| Δ21-Test-1 | AGCTGAATCTCGTCAGAGTG |
| Δ21-Test-2 | TCCTTCCTTCGCTTTAGCAAATC |
| Δ21-Test-3 | AAGTCCATTCAATATGTTCTGACC |
| Δ21-Test-4 | TAGTTAGTCGTGCGCTTCCAT |
| Δ22-N20-1 | AGAAACGACATCCCACAATCGTTTTAGAGCTAGAAATAGCAAGTTAAAATAAGG |
| Δ22-N20-2 | GATTGTGGGATGTCGTTTCTCGTAGGTACATTTTACTCAATTCTCTAATC |
| Δ22-N20-3 | ATCTCAGTAATCGCCAATGGGTTTTAGAGCTAGAAATAGCAAGTTAAAATAAGG |
| Δ22-N20-4 | CCATTGGCGATTACTGAGATCGTAGGTACATTTTACTCAATTCTCTAATC |
| Δ22-HA-1 | AGTGGTTCTATCGAGAGTCGTGTCAATGAACCGACTGTCTC |
| Δ22-HA-2 | ACAACAACACAGTATCTCCTGCGTTAAATTGGT |
| Δ22-HA-3 | AGGAGATACTGTGTTGTTGTTCCATCAGTAGATG |
| Δ22-HA-4 | CGGCAAATCCAATCGCATAGATCAGCAGCTGATATCGTTCAC |
| Δ22-Test-1 | TCAGCTGCCTACAGAAGATGAG |
| Δ22-Test-2 | ATCAGCAGCTGATATCGTTCAC |
| Δ22-Test-3 | TTCAGCCATATCACAGGAATCAG |
| Δ22-Test-4 | ACCATAATCGCGAGTGTTCC |
| Δ23-N20-1 | CGATACGCTGAAGGAATTCCGTTTTAGAGCTAGAAATAGCAAGTTAAAATAAGG |
| Δ23-N20-2 | GGAATTCCTTCAGCGTATCGCGTAGGTACATTTTACTCAATTCTCTAATC |

| | |
|---|---|
| Δ23-N20-3 | ATCGAAAGTGCTTGTATCCAGTTTTAGAGCTAGAAATAGCAAGTTAAAATAAGG |
| Δ23-N20-4 | TGGATACAAGCACTTTCGATCGTAGGTACATTTTACTCAATTCTCTAATC |
| Δ23-HA-1 | TTCTCCCCCATTACATCACTTGCACTCTGTCTTTAGATACAAGG |
| Δ23-HA-2 | CAGCTTAATTCATATGCAAGCTCAACCTGC |
| Δ23-HA-3 | GCTTGCATATGAATTAAGCTGATGACCTTCAGCAG |
| Δ23-HA-4 | GAGGTGACTGAAGTATACCGAAATTTGATTGGGATATGGCGCAA |
| Δ23-Test-1 | TCTGATTGGCAAGCTTGAGTG |
| Δ23-Test-2 | ATCGGATTTAACAGCAGGACA |
| Δ23-Test-3 | TGAAGCCTTTAGCAGAAGCTG |
| Δ23-Test-4 | ACAGTATGTCATTCTTGGAGCG |
| Δ24-N20-1 | TATTAAAGCAACTGACGGTGGTTTTAGAGCTAGAAATAGCAAGTTAAAATAAGG |
| Δ24-N20-2 | CACCGTCAGTTGCTTTAATACGTAGGTACATTTTACTCAATTCTCTAATCAC |
| Δ24-N20-3 | GAAATCCCTGTAAAATCAAGGTTTTAGAGCTAGAAATAGCAAGTTAAAATAAGG |
| Δ24-N20-4 | CTTGATTTTACAGGGATTTCCGTAGGTACATTTTACTCAATTCTCTAATCAC |
| Δ24-HA-1 | TTCTCCCCCATTACATCACTACGACCGTATCCTTGAAATTGT |
| Δ24-HA-2 | ATTCCGAAGTGAACTGATGACAACTGTACT |
| Δ24-HA-3 | TCATCAGTTCACTTCGGAATTTCTCAAGCGCAG |
| Δ24-HA-4 | GAGGTGACTGAAGTATACCGAATTGTCAGGCATTCGTTATCGG |
| Δ24-Test-1 | TGTGACAAGACCGTTTACCTT |
| Δ24-Test-2 | TGGCATGATCCAGATCAGGTC |
| Δ24-Test-3 | GATCAGACGACACAAGCTAATT |
| Δ24-Test-4 | AATCTCAGGTACAGCAGCTT |

| | |
|---|---|
| Δ25-N20-1 | TGATCAGGGTGTTGACTGAAGTTTTAGAGCTAGAAATAGCAAGTTAAAATAAGGC |
| Δ25-N20-2 | CTTCAGTCAACACCCTGATCACGTAGGTACATTTTACTCAATTCTCTAATCACGG |
| Δ25-N20-3 | ACATAAGCCTGAAGCGCAGGGTTTAGAGCTAGAAATAGCAAGTTAAAATAAGGC |
| Δ25-N20-4 | CCTGCGCTTCAGGCTTATGTCGTAGGTACATTTTACTCAATTCTCTAATCACGG |
| Δ25-HA-1 | GCTTCTCCCCCATTACATCACTTGCATCGCCTCAATACGCAA |
| Δ25-HA-2 | AGTATCTCCATGACCGATGAGTATCGTGCAGGATCAGG |
| Δ25-HA-3 | TACTCATCGGTCATGGAGATACTTGTGAAT |
| Δ25-HA-4 | AGGAGGTGACTGAAGTATACCGAAATCATGTCACCTGAATCAGTCAC |
| Δ25-Test-1 | ATGATACGCTCACAGAGGTC |
| Δ25-Test-2 | ATGTAGCGTCAATTAGCATTCC |
| Δ25-Test-3 | TGCTGATGAATGAAATCGAACG |
| Δ25-Test-4 | AATCGCCGTCAATCAGCTTAT |
| Δ26-N20-1 | CGTCGTACAGGATCTATTTAGTTTTAGAGCTAGAAATAGCAAGTTAAAATAAGG |
| Δ26-N20-2 | TAAATAGATCCTGTACGACGCGTAGGTACATTTTACTCAATTCTCTAATC |
| Δ26-N20-3 | GCGTTTGATTCACAGCCGATGTTTTAGAGCTAGAAATAGCAAGTTAAAATAAGG |
| Δ26-N20-4 | ATCGGCTGTGAATCAAACGCCGTAGGTACATTTTACTCAATTCTCTAATC |
| Δ26-HA-1 | AGTGGTTCTATCGAGAGTCGAACCATCGGACATGCATATGG |
| Δ26-HA-2 | CTGTCTTGAAGTCGCAAAGGTTTCAGTTGCA |
| Δ26-HA-3 | CCTTTGCGACTTCAAGACAGGCTTCGATATTAC |
| Δ26-HA-4 | CGGCAAATCCAATCGCATAGATGTACCGGTCATCGGGAAA |

| | |
|---|---|
| Δ26-Test-1 | TGGCAGCTGAAATCTAATGATGT |
| Δ26-Test-2 | TAGACATTCCGCAAAGCCAG |
| Δ26-Test-3 | TCCGATTACGGTCATCATCTT |
| Δ26-Test-4 | ATCAGCAGACATACTATGACCAT |
| Δ27-N20-1 | GCAGAGCTTAACCATGTGACGTTTTAGAGCTAGAAATA GCAAGTTAAAATAAGGC |
| Δ27-N20-2 | GTCACATGGTTAAGCTCTGCCGTAGGTACATTTTACTCA ATTCTCTAATCA |
| Δ27-N20-3 | GAGTCTCGTACACCTTGGAAGTTTTAGAGCTAGAAATA GCAAGTTAAAATAAGGC |
| Δ27-N20-4 | TTCCAAGGTGTACGAGACTCCGTAGGTACATTTTACTC AATTCTCTAATCA |
| Δ27-HA-1 | AGTGGTTCTATCGAGAGTCGAGCCTGCATTTACTGCTA TCT |
| Δ27-HA-2 | CCATCTTCACAACTGCTAATACTGCAAGACTGAT |
| Δ27-HA-3 | TATTAGCAGTTGTGAAGATGGTTCATGTTCATAACC |
| Δ27-HA-4 | CGGCAAATCCAATCGCATAGTCTCGTAATAGTCTTCTC CTGTCAC |
| Δ27-Test-1 | TCTGTCACTGACAACACTATGT |
| Δ27-Test-2 | AGTATCTGGATTCTCACCAGC |
| Δ27-Test-3 | TACGGTTATGATGAGCTTTGCT |
| Δ27-Test-4 | ACATTAATGGTGTCGACAAGAAG |
| Δ28-N20-1 | CGCTGCACCATAATATCCTGGTTTTAGAGCTAGAAATA GCAAGTTAAAATAAGGC |
| Δ28-N20-2 | CAGGATATTATGGTGCAGCGCGTAGGTACATTTTACTC AATTCTCTAATCA |
| Δ28-N20-3 | GGAGATGCGAGCGGATTTATGTTTTAGAGCTAGAAATA GCAAGTTAAAATAAGGC |
| Δ28-N20-4 | ATAAATCCGCTCGCATCTCCCGTAGGTACATTTTACTCA ATTCTCTAATCA |
| Δ28-HA-1 | TTCTCCCCCATTACATCACTACTTTCAGAGAGCTATTAA AGCTG |

| | |
|---|---|
| Δ28-HA-2 | GCGGCTATTATGTACCAAGTAAAAGTGATGCTTCT |
| Δ28-HA-3 | ACTTGGTACATAATAGCCGCCGACTTTATTCG |
| Δ28-HA-4 | GAGGTGACTGAAGTATACCGAAATCCTGGGATGATGTT GATTACAC |
| Δ28-Test-1 | TGAAGCTGCTGAGGAGCTAA |
| Δ28-Test-2 | ATGTCACACGCTTAATGCTT |
| Δ28-Test-3 | TTGTAACGATAGGAGTCTTGGC |
| Δ28-Test-4 | AGCTCGCAGACGTATGAAAT |
| Δ29-N20-1 | TGTCAGCCGTAATCTCAATCGTTTTAGAGCTAGAAATA GCAAGTTAAAATAAGGC |
| Δ29-N20-2 | GATTGAGATTACGGCTGACACGTAGGTACATTTTACTC AATTCTCTAATCACGG |
| Δ29-N20-3 | GCGAAGGGTGCATTCTTAGTGTTTTAGAGCTAGAAATA GCAAGTTAAAATAAGGC |
| Δ29-N20-4 | ACTAAGAATGCACCCTTCGCCGTAGGTACATTTTACTC AATTCTCTAATCACGG |
| Δ29-HA-1 | GCTTCTCCCCCATTACATCACTTGTCGTTCAATTTAAGG ACATTCCG |
| Δ29-HA-2 | GGTCAACTAGAAGCTGCTTCTGAAGAATACAACT |
| Δ29-HA-3 | TTCAGAAGCAGCTTCTAGTTGACCCTAACTGGACG |
| Δ29-HA-4 | AGGAGGTGACTGAAGTATACCGAAATGGTATACAATG ACATCGAATTGC |
| Δ29-Test-1 | AGAAGCCGAAGACCTGTATCA |
| Δ29-Test-2 | TGATAGAAGTGGCTTTACATTCCG |
| Δ29-Test-3 | ACCGTACTTACATGACTCACC |
| Δ29-Test-4 | ATATAGCCTTGGCTTGCTAATTC |
| Δ30-N20-1 | GGAAAAACAGAAGCAAGCCGGTTTTAGAGCTAGAAAT AGCAAGTTAAAATAAGG |
| Δ30-N20-2 | CGGCTTGCTTCTGTTTTTCCCGTAGGTACATTTTACTCA ATTCTCTAATCAC |
| Δ30-N20-3 | TCGCTCCGCAAAAGTAAATCGTTTTAGAGCTAGAAATA GCAAGTTAAAATAAGG |

| | |
|---|---|
| Δ30-N20-4 | GATTTACTTTTGCGGAGCGACGTAGGTACATTTTACTCAATTCTCTAATCAC |
| Δ30-HA-1 | GCTTCTCCCCCATTACATCACTTCAGCGGCATGACGATATTCT |
| Δ30-HA-2 | CTGATGCTGGTGTATTGTAGGTTTAGGGCTGATCG |
| Δ30-HA-3 | TAAACCTACAATACACCAGCATCAGGGTTTACA |
| Δ30-HA-4 | AGGAGGTGACTGAAGTATACCGAAAGAGTAGCTATATTGAAGAGGTCCA |
| Δ30-Test-1 | TATTGCCGTTACACTCAGAGCG |
| Δ30-Test-2 | ATGCATGAGCTCATTGAGTCTT |
| Δ30-Test-3 | TCAATGTCTTGCATCCTCATTCC |
| Δ30-Test-4 | TCTGAAGAAATGATTGTCACACCG |
| Δ31-N20-1 | TGCACACCTCTTCGTACGTAGTTTTAGAGCTAGAAATAGCAAGTTAAAATAAGGC |
| Δ31-N20-2 | TACGTACGAAGAGGTGTGCACGTAGGTACATTTTACTCAATTCTCTAATCA |
| Δ31-N20-3 | ATGCTGTCACCCTACTCTTTGTTTTAGAGCTAGAAATAGCAAGTTAAAATAAGGC |
| Δ31-N20-4 | AAAGAGTAGGGTGACAGCATCGTAGGTACATTTTACTCAATTCTCTAATCA |
| Δ31-HA-1 | AGTGGTTCTATCGAGAGTCGTGAACCTCTGACCATCACAGAC |
| Δ31-HA-2 | GGATGATTTATTATCAGCGCTACCGGACA |
| Δ31-HA-3 | GCGCTGATAATAAATCATCCGAACGGTTCCC |
| Δ31-HA-4 | CGGCAAATCCAATCGCATAGATGGATGTCATGAAGGTTATCAAG |
| Δ31-Test-1 | TGTTCTCGGTTCCAGAGGTTC |
| Δ31-Test-2 | TAAAGCACAACGCGTAACATG |
| Δ31-Test-3 | ATCACCAGCATGCAGGAAATA |
| Δ31-Test-4 | TAACAGGCCTAAGCATCGGAT |
| Δ32-N20-1 | CTACTTCCCATCCTATGCGCGTTTTAGAGCTAGAAATAGCAAGTTAAAATAAGG |

| | |
|---|---|
| Δ32-N20-2 | GCGCATAGGATGGGAAGTAGCGTAGGTACATTTTACTCAATTCTCTAATC |
| Δ32-N20-3 | TAAGACGGATAATCCGGAGAGTTTTAGAGCTAGAAATAGCAAGTTAAAATAAGG |
| Δ32-N20-4 | TCTCCGGATTATCCGTCTTACGTAGGTACATTTTACTCAATTCTCTAATC |
| Δ32-HA-1 | CTTCTCCCCCATTACATCACTTAAGTAAGTCAGCTGCTCGACAG |
| Δ32-HA-2 | CATTTCAGTGTGAAAGATGCAACAGGAGAAGAGC |
| Δ32-HA-3 | TGCATCTTTCACACTGAAATGTCCCAAACC |
| Δ32-HA-4 | GAGGTGACTGAAGTATACCGAAAATCGGTAAAATCAACACTTGCA |
| Δ32-Test-1 | ATACAGCTGTAAAGGTACAGCC |
| Δ32-Test-2 | TGATGCACTTGCCATCTGC |
| Δ32-Test-3 | TCATGCAATCTGACAAGCCC |
| Δ32-Test-4 | ACAGGCCATACAACCATCAA |
| Δ33-N20-1 | CCATACTAGTAGTCGCAGCAGTTTTAGAGCTAGAAATAGCAAGTTAAAATAAGG |
| Δ33-N20-2 | TGCTGCGACTACTAGTATGGCGTAGGTACATTTTACTCAATTCTCTAATC |
| Δ33-N20-3 | AGCAACTTCAATCATTTGCTGTTTTAGAGCTAGAAATAGCAAGTTAAAATAAGG |
| Δ33-N20-4 | AGCAAATGATTGAAGTTGCTCGTAGGTACATTTTACTCAATTCTCTAATC |
| Δ33-HA-1 | GCTTCTCCCCCATTACATCACTTCCGCATATTGTGAAGACGATC |
| Δ33-HA-2 | TAACCTGTCTTGAGATGTATACCGCTATGCCACAG |
| Δ33-HA-3 | CGGTATACATCTCAAGACAGGTTATTGTACAGG |
| Δ33-HA-4 | AGGAGGTGACTGAAGTATACCGAAATATGAATACGCCAATTCTACCGG |
| Δ33-Test-1 | TTGCGACAACAGCTTCAATG |
| Δ33-Test-2 | ACGCGGATAACATTTCAAGAAC |

| | |
|---|---|
| Δ33-Test-3 | ATCGAGCTTCTCAACTTGATACT |
| Δ33-Test-4 | CAGAAGTGAATGAACTGGCGAA |
| Δ34-N20-1 | TCCTTCGTGTCTTGCTTGGCGTTTTAGAGCTAGAAATAG CAAGTTAAAATAAGG |
| Δ34-N20-2 | GCCAAGCAAGACACGAAGGACGTAGGTACATTTTACTC AATTCTCTAATC |
| Δ34-N20-3 | GTTGTTCCGGTATTTCAGAAGTTTTAGAGCTAGAAATA GCAAGTTAAAATAAGG |
| Δ34-N20-4 | TTCTGAAATACCGGAACAACCGTAGGTACATTTTACTC AATTCTCTAATC |
| Δ34-HA-1 | AGTGGTTCTATCGAGAGTCGACGTGACATTCACTTCTTC ACA |
| Δ34-HA-2 | ATTGAGCTGACGAGATGTCGCTGTCACATTTGAA |
| Δ34-HA-3 | CGACATCTCGTCAGCTCAATCATAGAGGAT |
| Δ34-HA-4 | CGGCAAATCCAATCGCATAGATCCGTCAGATCTTCAGC TATC |
| Δ34-Test-1 | TCACTTCTTCACATAATTCAGCGA |
| Δ34-Test-2 | AGGCTACGGTGACTATTTCG |
| Δ34-Test-3 | TGATTACTGTATCTGTTGCGTCTG |
| Δ34-Test-4 | ATAAATCATTCTCACCGCAAGAAT |
| Δ35-N20-1 | CCGCGACTATGTTATTGGACGTTTTAGAGCTAGAAATA GCAAGTTAAAATAAGGC |
| Δ35-N20-2 | GTCCAATAACATAGTCGCGGCGTAGGTACATTTTACTC AATTCTCTAATCACGG |
| Δ35-N20-3 | ATCGAATAGAGCGAGCCTGTGTTTTAGAGCTAGAAATA GCAAGTTAAAATAAGGC |
| Δ35-N20-4 | ACAGGCTCGCTCTATTCGATCGTAGGTACATTTTACTCA ATTCTCTAATCACGG |
| Δ35-HA-1 | GCTTCTCCCCCATTACATCACTTACCGCAAGCAGCCAA TATACA |
| Δ35-HA-2 | AACAAGCATCAGTATCATATCCAACGTAATGACGCC |
| Δ35-HA-3 | GTTGGATATGATACTGATGCTTGTTGAAGATGC |

| | |
|---|---|
| Δ35-HA-4 | AGGAGGTGACTGAAGTATACCGAAATGTAGGGATAGCAGGAGACAC |
| Δ35-Test-1 | GTGGAAGAATGATATGACATTGCT |
| Δ35-Test-2 | ACTGCTGAGATGGACAGAATAA |
| Δ35-Test-3 | TTGATAAGTTCACACTCGCTGAA |
| Δ35-Test-4 | CAGTGTTTGGGTATACGCTGA |
| Δ36-N20-1 | TTCATTAGTTTGGCGCCTGAGTTTTAGAGCTAGAAATAGCAAGTTAAAATAAGGC |
| Δ36-N20-2 | TCAGGCGCCAAACTAATGAACGTAGGTACATTTTACTCAATTCTCTAATCACGG |
| Δ36-N20-3 | TGAGAGCCTCCCGTTATGAGGTTTTAGAGCTAGAAATAGCAAGTTAAAATAAGGC |
| Δ36-N20-4 | CTCATAACGGGAGGCTCTCACGTAGGTACATTTTACTCAATTCTCTAATCACGG |
| Δ36-HA-1 | AGTGGTTCTATCGAGAGTCGACACGAACCAATCTTATTGCAG |
| Δ36-HA-2 | GTGAGAAGAATAGATGTCGGTTACAAATGCAATTAGC |
| Δ36-HA-3 | ACCGACATCTATTCTTCTCACGTCTATCATCCT |
| Δ36-HA-4 | CGGCAAATCCAATCGCATAGATAAACCTGTACCGATTGCAC |
| Δ36-Test-1 | ATCAGATTGAGGGTTACCGTG |
| Δ36-Test-2 | TTGATAAGAGCAGTTCTCCGAG |
| Δ36-Test-3 | TACAGACCAATTCGAAGACATCG |
| Δ36-Test-4 | ATGATCGTCTTGTGATTGATGC |
| Δ37-N20-1 | GGCATCTGAAATCCGGACTGGTTTTAGAGCTAGAAATAGCAAGTTAAAATAAGGC |
| Δ37-N20-2 | CAGTCCGGATTTCAGATGCCCGTAGGTACATTTTACTCAATTCTCTAATCA |
| Δ37-N20-3 | AGAAGAACTGAAAGAGCTGCGTTTTAGAGCTAGAAATAGCAAGTTAAAATAAGGC |
| Δ37-N20-4 | GCAGCTCTTTCAGTTCTTCTCGTAGGTACATTTTACTCAATTCTCTAATCA |

| | |
|---|---|
| Δ37-HA-1 | AGGCTGTCATCATTGAGCTT |
| Δ37-HA-2 | AGCTGCAAGATCAAAGACGT |
| Δ37-HA-3 | AAAGCCGATCCTGAGTTTAC |
| Δ37-HA-4 | AGCTGCATTTTGTGCAAAAC |
| Δ37-Test-1 | ATAGGAGATTCCGTCATGCTT |
| Δ37-Test-2 | TCAACAGCCGCTTACTTTAGG |
| Δ37-Test-3 | TAATGCAAGCCTAATATCGGTGC |
| Δ37-Test-4 | AATCAACGTTATTGTGAGGCAG |
| Δ38-N20-1 | TGTATTCCAGCGTATTCCAAGTTTTAGAGCTAGAAATAGCAAGTTAAAATAAG |
| Δ38-N20-2 | TTGGAATACGCTGGAATACACGTAGGTACATTTTACTCAATTCTCTAATC |
| Δ38-N20-3 | GGTCGACGGCCAACGTGATCAGAAATTGAGGTCCTCTG |
| Δ38-N20-4 | CGACTCTCGATAGAACCACTTGGTGTTGATTAATCTGTGCGG |
| Δ38-HA-1 | TTCTCCCCCATTACATCACTTGATCAGAAATTGAGGTCCTCTG |
| Δ38-HA-2 | TGAGGTGACATGATTATTGCCGGTGACATTGG |
| Δ38-HA-3 | GCAATAATCATGTCACCTCATCAGCAAATGG |
| Δ38-HA-4 | GAGGTGACTGAAGTATACCGAATGGTGTTGATTAATCTGTGCGG |
| Δ38-Test-1 | ATAACGGAATCAGATTCCGGT |
| Δ38-Test-2 | ATTCAATCAGCTTGCTGATGC |
| Δ38-Test-3 | TAATCATCGCACCGAAAGCC |
| Δ38-Test-4 | TTGTATCAGCAGGGACTTGC |
| Δ39-N20-1 | GGGTCTTCCTGAAATCCTGAGTTTTAGAGCTAGAAATAGCAAGTTAAAATAAGGC |
| Δ39-N20-2 | TCAGGATTTCAGGAAGACCCCGTAGGTACATTTTACTCAATTCTCTAATCA |
| Δ39-N20-3 | ATAGACAAGCAGAACAGGGAGTTTTAGAGCTAGAAATAGCAAGTTAAAATAAGGC |

| | |
|---|---|
| Δ39-N20-4 | TCCCTGTTCTGCTTGTCTATCGTAGGTACATTTTACTCAATTCTCTAATCA |
| Δ39-HA-1 | CTTCTCCCCCATTACATCACTAATTGGCCTTTTCCTGTACC |
| Δ39-HA-2 | TTTGTCTCAGCTGAAGCAGG |
| Δ39-HA-3 | CCTGCTTCAGCTGAGACAAAGTCAATTTTCGCGGACTCTT |
| Δ39-HA-4 | AGGAGGTGACTGAAGTATACCGAATGACCTCTGAACAGATCGGC |
| Δ39-Test-1 | TGTCCGCATCTATATTCAGAAGC |
| Δ39-Test-2 | AAGCTATTATCTGTGCCGTGC |
| Δ39-Test-3 | TCTGTCAGCTCACGGTATTCT |
| Δ39-Test-4 | ATGCAGGCTGATGAACTAGATAT |
| Δ40-N20-1 | TCCAGTGTCTTCATGAGAGCGTTTTAGAGCTAGAAATAGCAAGTTAAAATAAGG |
| Δ40-N20-2 | GCTCTCATGAAGACACTGGACGTAGGTACATTTTACTCAATTCTCTAATC |
| Δ40-N20-3 | ATTACCGGTGATGTCGTGATGTTTTAGAGCTAGAAATAGCAAGTTAAAATAAGG |
| Δ40-N20-4 | ATCACGACATCACCGGTAATCGTAGGTACATTTTACTCAATTCTCTAATC |
| Δ40-HA-1 | AGTGGTTCTATCGAGAGTCGAGCACGCTGTATAAGGAAAGAG |
| Δ40-HA-2 | TTGTTGAAGAAGTGCAATGAGACATGCCAGA |
| Δ40-HA-3 | CTCATTGCACTTCTTCAACAAACATAAGACCTGCT |
| Δ40-HA-4 | CGGCAAATCCAATCGCATAGAATCGTTAGCCAAAGCAATCG |
| Δ40-Test-1 | ATAGTCAGGATCACCATCGCA |
| Δ40-Test-2 | ATTGCAACCGGAATCAGCAA |
| Δ40-Test-3 | TCCAACGGACGACAACACAA |
| Δ40-Test-4 | AGTATTGAAGGATGTTTCATTGAGC |

| | |
|---|---|
| Δ41-N20-1 | CAGAGCGCCATATGACAGTGGTTTTAGAGCTAGAAATAGCAAGTTAAAATAAGG |
| Δ41-N20-2 | CACTGTCATATGGCGCTCTGCGTAGGTACATTTTACTCAATTCTCTAATC |
| Δ41-N20-3 | GAACTGCGATCGGCATCATTGTTTTAGAGCTAGAAATAGCAAGTTAAAATAAGG |
| Δ41-N20-4 | AATGATGCCGATCGCAGTTCCGTAGGTACATTTTACTCAATTCTCTAATC |
| Δ41-HA-1 | TTCTCCCCCATTACATCACTTGTCTGCTGATCGTCCAGAAG |
| Δ41-HA-2 | CTCCATAAATGCTGAAGCTGTTCTTTCGT |
| Δ41-HA-3 | CAGCTTCAGCATTTATGGAGTGCATTTGCGACG |
| Δ41-HA-4 | GAGGTGACTGAAGTATACCGAAATCACTGATCCGTTTAGAACCG |
| Δ41-Test-1 | TGCCAGACAGGAAGACATGAC |
| Δ41-Test-2 | TTGTCTGCCATCATTACAGAGGC |
| Δ41-Test-3 | TCATTAAGCAGCACCTTCAGC |
| Δ41-Test-4 | ATTCGCTCACTGTTGTGATC |
| Δ42-N20-1 | CGGGATATCCTTATAGCCGTGTTTTAGAGCTAGAAATAGCAAGTTAAAATAAGG |
| Δ42-N20-2 | ACGGCTATAAGGATATCCCGCGTAGGTACATTTTACTCAATTCTCTAATC |
| Δ42-N20-3 | AACACCTGTCTCAGCATAAAGTTTTAGAGCTAGAAATAGCAAGTTAAAATAAGG |
| Δ42-N20-4 | TTTATGCTGAGACAGGTGTTCGTAGGTACATTTTACTCAATTCTCTAATC |
| Δ42-HA-1 | TTCTCCCCCATTACATCACTAAGCCGTTCTCTTTATGAACCA |
| Δ42-HA-2 | TTTTTTCGATGCCAGCATTAACC |
| Δ42-HA-3 | TAATGCTGGCATCGAAAAAACATAAACAACCGTTGCATAGAGT |

| | |
|---|---|
| Δ42-HA-4 | GGTGACTGAAGTATACCGAAAACGAAGAAGGCTTTGAATCATT |
| Δ42-Test-1 | TCTTGAGTCAGCAGCGGATAT |
| Δ42-Test-2 | AATCAGGTACGACAACTGAACC |
| Δ42-Test-3 | TGTCACAACCATCGTATTTGTCG |
| Δ42-Test-4 | TAGTTCATCTGCACTGACGC |
| Δ43-N20-1 | TGTACCGTCCGATGATCCACGTTTTAGAGCTAGAAATAGCAAGTTAAAATAAGG |
| Δ43-N20-2 | AGTGATGTAATGGGGGAGAAGC |
| Δ43-N20-3 | ATCGTCATGTTGACCTTGCAGTTTTAGAGCTAGAAATAGCAAGTTAAAATAAGG |
| Δ43-N20-4 | TGGTAGCTCTTGATCCGGCA |
| Δ43-HA-1 | GCTTCTCCCCCATTACATCACTCGCAGGATTCATTACAGGCT |
| Δ43-HA-2 | CTGTCCGGATCAAGGAGAAAAAAAACAGCCGGAACTCTGC |
| Δ43-HA-3 | TTTCTCCTTGATCCGGACAG |
| Δ43-HA-4 | GGAGGTGACTGAAGTATACCGAAATCAGGCGCCTTATAGTTTGT |
| Δ43-Test-1 | TACTGTACTCTGTGCTGAGG |
| Δ43-Test-2 | AACTGGGACACTTGCTCAGA |
| Δ43-Test-3 | CCATCAATCCATCACTGGTC |
| Δ43-Test-4 | TCGCCACATATCAAAATGGG |
| Δ44-N20-1 | TCGTCAGGCTTTGTCTCCAAGTTTTAGAGCTAGAAATAGCAAGTTAAAATAAGG |
| Δ44-N20-2 | TTGGAGACAAAGCCTGACGACGTAGGTACATTTTACTCAATTCTCTAATC |
| Δ44-N20-3 | AAGTGGCACTGGATTATGTGGTTTTAGAGCTAGAAATAGCAAGTTAAAATAAGG |
| Δ44-N20-4 | CACATAATCCAGTGCCACTTCGTAGGTACATTTTACTCAATTCTCTAATC |

| | |
|---|---|
| Δ44-HA-1 | TTCTCCCCCATTACATCACTAGATAAGCCGCGCAAATATTC |
| Δ44-HA-2 | CTGAATCAGATGATATCGCTGTGAATCGTGAAT |
| Δ44-HA-3 | CAGCGATATCATCTGATTCAGTTTGCAGATCAGTG |
| Δ44-HA-4 | GAGGTGACTGAAGTATACCGAATAGGAACAGATTATCCGATCTACGG |
| Δ44-Test-1 | TCACAATGTTGAGCAAAGCTGT |
| Δ44-Test-2 | TGTTCGAAGCACCTACTGTAG |
| Δ44-Test-3 | GAATGTAGACAATCGCATCGTAT |
| Δ44-Test-4 | TCTGCGATTCTCACAAGTCGT |
| Δ45-N20-1 | ACAGGACAAATCGTTGAGCCGTTTTAGAGCTAGAAATAGCAAGTTAAAATAAGGC |
| Δ45-N20-2 | GGCTCAACGATTTGTCCTGTCGTAGGTACATTTTACTCAATTCTCTAATCA |
| Δ45-N20-3 | CGCGTCAGTATGAATGCTGAGTTTTAGAGCTAGAAATAGCAAGTTAAAATAAGGC |
| Δ45-N20-4 | TCAGCATTCATACTGACGCGCGTAGGTACATTTTACTCAATTCTCTAATCA |
| Δ45-HA-1 | AGTGGTTCTATCGAGAGTCGATAATATCCACGTTGTTCGCCA |
| Δ45-HA-2 | AGACATGGCATTCGACAGCACTGGATCTGAA |
| Δ45-HA-3 | TGCTGTCGAATGCCATGTCTTGATCTTGTACA |
| Δ45-HA-4 | CGGCAAATCCAATCGCATAGTCTAACATCGAGTGTGATACGCT |
| Δ45-Test-1 | TCTGCAAATTCATCAGGATGGT |
| Δ45-Test-2 | TGATTCACCTTGCACCAAACA |
| Δ45-Test-3 | AGATATGGCATTCAACACCTGT |
| Δ45-Test-4 | TCAAGCATCTGACACAGTCT |
| Δ46-N20-1 | ATGTCGCAGACAACTACTGCGTTTTAGAGCTAGAAATAGCAAGTTAAAATAAGGC |
| Δ46-N20-2 | GCAGTAGTTGTCTGCGACATCGTAGGTACATTTTACTCAATTCTCTAATCA |

| | |
|---|---|
| Δ46-N20-3 | CCATCGTTTCAATCGCTTCAGTTTTAGAGCTAGAAATAGCAAGTTAAAATAAGGC |
| Δ46-N20-4 | TGAAGCGATTGAAACGATGGCGTAGGTACATTTTACTCAATTCTCTAATCA |
| Δ46-HA-1 | AGTGGTTCTATCGAGAGTCGATACACACAGCCCAATTGTCT |
| Δ46-HA-2 | TTGGCATCATATCTGCAGCATGGAGTTGTAA |
| Δ46-HA-3 | TGCTGCAGATATGATGCCAATCACACTAAGC |
| Δ46-HA-4 | CGGCAAATCCAATCGCATAGTCGTGAACGTCATGTTCCATG |
| Δ46-Test-1 | TTCCAGCAGCCTGTCATTTTC |
| Δ46-Test-2 | TCTTATCGCTATGCTGGGCAG |
| Δ46-Test-3 | TTGCCGTCAGTTAGTCCGTAC |
| Δ46-Test-4 | ATCCACTCAGAGGTATACGAGT |
| Δ47-N20-1 | ATCGGAGACCTCGTATCACAGTTTTAGAGCTAGAAATAGCAAGTTAAAATAAGGC |
| Δ47-N20-2 | TGTGATACGAGGTCTCCGATCGTAGGTACATTTTACTCAATTCTCTAATCA |
| Δ47-N20-3 | GCGGACAGATTTTTAAGGCTGTTTTAGAGCTAGAAATAGCAAGTTAAAATAAGGC |
| Δ47-N20-4 | AGCCTTAAAAATCTGTCCGCCGTAGGTACATTTTACTCAATTCTCTAATCA |
| Δ47-HA-1 | CTTCTCCCCCATTACATCACTCGAGCAGAAGATCAGGCAAT |
| Δ47-HA-2 | CGAAAGCGGGCAAAACAAAT |
| Δ47-HA-3 | ATTTGTTTTGCCCGCTTTCGTGAAGCGGTTTACAAGTTGG |
| Δ47-HA-4 | AGGAGGTGACTGAAGTATACCGAAGGCAGCGAAAACAGACAATC |
| Δ47-Test-1 | TGATTCCGAGACTGGCAGAC |
| Δ47-Test-2 | TGATTCCGAGACTGGCAGAC |
| Δ47-Test-3 | ACATAGAACGAATTCCGAATGG |

| | |
|---|---|
| Δ47-Test-4 | TTATATGCTGCTGGCTTTAGGC |
| Δ48-N20-1 | GCAGTAGCACTTCAATGACCGTTTTAGAGCTAGAAATAGCAAGTTAAAATAAGGC |
| Δ48-N20-2 | GGTCATTGAAGTGCTACTGCCGTAGGTACATTTTACTCAATTCTCTAATCA |
| Δ48-N20-3 | AACCACTTTTCCTTTCAGCGGTTTTAGAGCTAGAAATAGCAAGTTAAAATAAGGC |
| Δ48-N20-4 | CGCTGAAAGGAAAAGTGGTTCGTAGGTACATTTTACTCAATTCTCTAATCA |
| Δ48-HA-1 | AGTGGTTCTATCGAGAGTCGAAGAGCCGCGATATCATCTGA |
| Δ48-HA-2 | CCGATTTCTCTTTCATACCTGGATGGAAGGCG |
| Δ48-HA-3 | AGGTATGAAAGAGAAATCGGCTTGTCTGTC |
| Δ48-HA-4 | CGGCAAATCCAATCGCATAGTTCTTTCGTCATCGTTCGAGAAG |
| Δ48-Test-1 | TATTCAGTTCCGCTGCGATG |
| Δ48-Test-2 | ACCGTATGCAATTTATCCAGTTG |
| Δ48-Test-3 | TCAACGATTGTAACGACATGTC |
| Δ48-Test-4 | ATTCAGACGATTGGATTGAGC |
| Δ49-N20-1 | CTTGTCTCACCAATCGTGATGTTTTAGAGCTAGAAATAGCAAGTTAAAATAAGGC |
| Δ49-N20-2 | ATCACGATTGGTGAGACAAGCGTAGGTACATTTTACTCAATTCTCTAATCA |
| Δ49-N20-3 | CTGATCATTTCTGCCAGGGAGTTTTAGAGCTAGAAATAGCAAGTTAAAATAAGGC |
| Δ49-N20-4 | TCCCTGGCAGAAATGATCAGCGTAGGTACATTTTACTCAATTCTCTAATCA |
| Δ49-HA-1 | CTTCTCCCCCATTACATCACTACGCATTAGAGCTTCCTTTT |
| Δ49-HA-2 | TTGAAACAGGCAGTGAAGTC |
| Δ49-HA-3 | GACTTCACTGCCTGTTTCAAGGCACGATGAACTTTGAATG |

| | |
|---|---|
| Δ49-HA-4 | AGGAGGTGACTGAAGTATACCGAAAATGCAAGGGCAGTAAATGA |
| Δ49-Test-1 | TGTCTGGATGTACAGGTTCGA |
| Δ49-Test-2 | ATTGTTGGTTGAAAACAGGTCC |
| Δ49-Test-3 | AATGCCGATATCCTATTGGCA |
| Δ49-Test-4 | TGTGTAATGCCAGTCCTTAAGAT |
| Δ50-N20-1 | GACGCCGGAAGATCTTTTACGTTTTAGAGCTAGAAATAGCAAGTTAAAATAAGG |
| Δ50-N20-2 | GTAAAAGATCTTCCGGCGTCCGTAGGTACATTTTACTCAATTCTCTAATC |
| Δ50-N20-3 | TACCTCGTTGATATCGGCTTGTTTTAGAGCTAGAAATAGCAAGTTAAAATAAGG |
| Δ50-N20-4 | AAGCCGATATCAACGAGGTACGTAGGTACATTTTACTCAATTCTCTAATC |
| Δ50-HA-1 | AGTGGTTCTATCGAGAGTCGTTCCTGTTGTTCTATTGGACGC |
| Δ50-HA-2 | GGAAGTTCGATTGTTACATGAGGAATGGGAGCA |
| Δ50-HA-3 | TCATGTAACAATCGAACTTCCACTTTCTGTTGA |
| Δ50-HA-4 | CGGCAAATCCAATCGCATAGATACGTCACGATACACATAAAGTGG |
| Δ50-Test-1 | TGTTACTCTAAGGTATGCAATTCCC |
| Δ50-Test-2 | TGATATCGAAGAGCTGAAAGCG |
| Δ50-Test-3 | TGAACGGCAATGACATCGTG |
| Δ50-Test-4 | TAGGCAGGCTGTATCTACTGA |
| Δ51-N20-1 | ATTCATATCCACGGCGGCTAGTTTTAGAGCTAGAAATAGCAAGTTAAAATAAGG |
| Δ51-N20-2 | TAGCCGCCGTGGATATGAATCGTAGGTACATTTTACTCAATTCTCTAATC |
| Δ51-N20-3 | TGTTGATTGTGTGCTCTAGAGTTTTAGAGCTAGAAATAGCAAGTTAAAATAAGG |
| Δ51-N20-4 | TCTAGAGCACACAATCAACACGTAGGTACATTTTACTCAATTCTCTAATC |

| | |
|---|---|
| Δ51-HA-1 | TTCTCCCCCATTACATCACTAGCTTGTTGGTAAATTGTTCAGC |
| Δ51-HA-2 | ATTCAACGGCATCATTGATTCCGACATAGCCGT |
| Δ51-HA-3 | GAATCAATGATGCCGTTGAATTCAGTACGAT |
| Δ51-HA-4 | GAGGTGACTGAAGTATACCGAATTGATGACATTCGTGATGTATTCGC |
| Δ51-Test-1 | TCGGATTGAGTGGTTCTATCG |
| Δ51-Test-2 | AGCTTCATAAAGCGAAAGCG |
| Δ51-Test-3 | TGCAGATACCATCCATTGGCA |
| Δ51-Test-4 | TCTTGGTATCAAGAGCGCTT |
| Δ52-N20-1 | GAATTTAGCTCCGCCGATAAGTTTTAGAGCTAGAAATAGCAAGTTAAAATAAGG |
| Δ52-N20-2 | TTATCGGCGGAGCTAAATTCCGTAGGTACATTTTACTCAATTCTCTAATC |
| Δ52-N20-3 | CTTGATCGGCAGTTGTTTGCGTTTTAGAGCTAGAAATAGCAAGTTAAAATAAGG |
| Δ52-N20-4 | GCAAACAACTGCCGATCAAGCGTAGGTACATTTTACTCAATTCTCTAATC |
| Δ52-HA-1 | CTTCTCCCCCATTACATCACTAGAACTTCAGCGATTCTCGAA |
| Δ52-HA-2 | CATGAGCGAGTTATGTACAGCCTGTATTCATTGCT |
| Δ52-HA-3 | GCTGTACATAACTCGCTCATGTTGATGTCT |
| Δ52-HA-4 | GAGGTGACTGAAGTATACCGAATTCTTGACACCAGCTTATGCAA |
| Δ52-Test-1 | TGGCACCATTTATATTTGGTCTCT |
| Δ52-Test-2 | GTGCAGACAAGAACAGTAGAACT |
| Δ52-Test-3 | TATATGGAAGAACGATCACAGCG |
| Δ52-Test-4 | TATCTATAATGGCGCTCAGCAC |
| Δ53-N20-1 | CGCACAAGAGGAACTTTGGAGTTTTAGAGCTAGAAATAGCAAGTTAAAATAAGG |
| Δ53-N20-2 | TCCAAAGTTCCTCTTGTGCGCGTAGGTACATTTTACTCAATTCTCTAATC |

| | |
|---|---|
| Δ53-N20-3 | AACTGCAGGATACCTGAAGAGTTTTAGAGCTAGAAATAGCAAGTTAAAATAAGG |
| Δ53-N20-4 | TCTTCAGGTATCCTGCAGTTCGTAGGTACATTTTACTCAATTCTCTAATC |
| Δ53-HA-1 | AGTGGTTCTATCGAGAGTCGCCGATTGGCGTAAGTAATATTGT |
| Δ53-HA-2 | ACGTGAAACTGAATTGAGCTTATTAACGCAAGTGG |
| Δ53-HA-3 | AGCTCAATTCAGTTTCACGTATTCTTTCGCTG |
| Δ53-HA-4 | CGGCAAATCCAATCGCATAGTTACCAAACGGATATCAGAAGCG |
| Δ53-Test-1 | TGCTCTGCTTTCTCATGACC |
| Δ53-Test-2 | TGTCAGTATTTCGGTCGCATC |
| Δ53-Test-3 | TACATCTGCAAGAGACAGACG |
| Δ53-Test-4 | TCAATAACACCGGCTTCAATTAAT |
| Δ54-N20-1 | CAATTCGTCTGTAAATCGAAGTTTTAGAGCTAGAAATAGCAAGTTAAAATAAGG |
| Δ54-N20-2 | TTCGATTTACAGACGAATTGCGTAGGTACATTTTACTCAATTCTCTAATCA |
| Δ54-N20-3 | ACAGGCGCTGGAGACGTATAGTTTTAGAGCTAGAAATAGCAAGTTAAAATAAGG |
| Δ54-N20-4 | TATACGTCTCCAGCGCCTGTCGTAGGTACATTTTACTCAATTCTCTAATCA |
| Δ54-HA-1 | TTCTCCCCCATTACATCACTGAGTGCTTTATGATGAACTCGG |
| Δ54-HA-2 | GGGAAGTGTATATCACTGACGAG |
| Δ54-HA-3 | CTCGTCAGTGATATACACTTCCCTCCTCTTCCATCTGTTCGATATCTAG |
| Δ54-HA-4 | GGTGACTGAAGTATACCGAACGGGTATTCGTGACATCGC |
| Δ54-Test-1 | TCAGGTCAGAGTGCTTTATGA |
| Δ54-Test-2 | AACTTCCGATGCACGTACTG |
| Δ54-Test-3 | AGAGGCGAGCTTCATAATATCC |

| | |
|---|---|
| Δ54-Test-4 | TCACAGTATACGGTGAAGAAATGATC |
| Δ55-N20-1 | AAGAGTTCACCAGCTTCCTGGTTTTAGAGCTAGAAATAGCAAGTTAAAATAAGG |
| Δ55-N20-2 | CAGGAAGCTGGTGAACTCTTCGTAGGTACATTTTACTCAATTCTCTAATC |
| Δ55-N20-3 | TATGATGACTTGGCACAACAGTTTTAGAGCTAGAAATAGCAAGTTAAAATAAGG |
| Δ55-N20-4 | TGTTGTGCCAAGTCATCATACGTAGGTACATTTTACTCAATTCTCTAATC |
| Δ55-HA-1 | AGTGGTTCTATCGAGAGTCGATCGCCATATTCGTTGAGTAGT |
| Δ55-HA-2 | CGGCAAATCCAATCGCATAGACACTTGTTTACCGCCATCA |
| Δ55-HA-3 | GTCCTATAGTGAACTGCTGTTTCTGATTATCC |
| Δ55-HA-4 | GAGGTGACTGAAGTATACCGAAACACTTGTTTACCGCCATCA |
| Δ55-Test-1 | TGCTTCCAGTACAGACGGAA |
| Δ55-Test-2 | ATTATGCAGCACAGTCTGGC |
| Δ55-Test-3 | TCGGAGTTGTGCTAAGCAAC |
| Δ55-Test-4 | TCATCCGCAGTTAACCTGATATAC |
| Δ56-N20-1 | GTCTGACCAGCATACGAGAGGTTTTAGAGCTAGAAATAGCAAGTTAAAATAAGG |
| Δ56-N20-2 | CTCTCGTATGCTGGTCAGACCGTAGGTACATTTTACTCAATTCTCTAATCA |
| Δ56-N20-3 | GGGACGATGGTATATTCCAGGTTTTAGAGCTAGAAATAGCAAGTTAAAATAAGG |
| Δ56-N20-4 | CTGGAATATACCATCGTCCCCGTAGGTACATTTTACTCAATTCTCTAATCAC |
| Δ56-HA-1 | GCTTCTCCCCCATTACATCACTGGGATGTTCAAGGTTTCTCC |
| Δ56-HA-2 | AAGATTGGTTGGTGCTCATGTGCGGTTCGTTTTGACAAAT |

| | |
|---|---|
| Δ56-HA-3 | CATGAGCACCAACCAATCTT |
| Δ56-HA-4 | GGAGGTGACTGAAGTATACCGAATCGGTGATCGTGTTCCTTAT |
| Δ56-Test-1 | CTAGCATACTCTTCTTCCAATCAAAT |
| Δ56-Test-2 | AATACTCCAGAGAACTGAGCT |
| Δ56-Test-3 | ACAGAAACAGCGAATATCGGT |
| Δ56-Test-4 | TAGACAGAATTAAGATCATGCTGTGG |
| Δ57-N20-1 | ACGACAAGAGATTCAGTGAAGTTTTAGAGCTAGAAATAGCAAGTTAAAATAAGG |
| Δ57-N20-2 | TTCACTGAATCTCTTGTCGTCGTAGGTACATTTTACTCAATTCTCTAATC |
| Δ57-N20-3 | GAAATTGAGCGCGCTGTTTTGTTTTAGAGCTAGAAATAGCAAGTTAAAATAAGG |
| Δ57-N20-4 | AAAACAGCGCGCTCAATTTCCGTAGGTACATTTTACTCAATTCTCTAATC |
| Δ57-HA-1 | AGTGGTTCTATCGAGAGTCGTCCTCATTTGTGATGACGTCAG |
| Δ57-HA-2 | CGAAGTCGATAAAGGCGATGTAGTAAACGGT |
| Δ57-HA-3 | CATCGCCTTTATCGACTTCGCATTTCGTTATCC |
| Δ57-HA-4 | CGGCAAATCCAATCGCATAGAGCGTGATTTATGCCTTCAGT |
| Δ57-Test-1 | TGACGCCTGAATGTTATGTGC |
| Δ57-Test-2 | ATACCGCTGACTATGAGGATATG |
| Δ57-Test-3 | TGATCAAGCTTCCCGTCATG |
| Δ57-Test-4 | ATACTGTACGCCAAATGGATG |
| Δ58-N20-1 | ACATAACTGAAGCCAACTTGGTTTTAGAGCTAGAAATAGCAAGTTAAAATAAGG |
| Δ58-N20-2 | CAAGTTGGCTTCAGTTATGTCGTAGGTACATTTTACTCAATTCTCTAATC |
| Δ58-N20-3 | GCCTGCTTGAAGCTGATTGAGTTTTAGAGCTAGAAATAGCAAGTTAAAATAAGG |

| | |
|---|---|
| Δ58-N20-4 | TCAATCAGCTTCAAGCAGGCCGTAGGTACATTTTACTCAATTCTCTAATC |
| Δ58-HA-1 | AGTGGTTCTATCGAGAGTCGAGATCCATCTCATCCAATGCG |
| Δ58-HA-2 | GGATGTCTTTGAACATCCATGAATTGGTTGAGC |
| Δ58-HA-3 | ATGGATGTTCAAAGACATCCGCTCAGTCT |
| Δ58-HA-4 | CGGCAAATCCAATCGCATAGACCAATGAAACACCTTAATCATGC |
| Δ58-Test-1 | TATTGAATGCCATTCTCTTGGATGA |
| Δ58-Test-2 | TAGGTATGGGTTGCTGCCAAA |
| Δ58-Test-3 | ATCATCGCCATAATATCAAACGAAG |
| Δ58-Test-4 | ACATCAGCATCACTGTCCAG |
| Δ59-N20-1 | CGCTCACGTATCAGGAATTAGTTTTAGAGCTAGAAATAGCAAGTTAAAATAAGG |
| Δ59-N20-2 | TAATTCCTGATACGTGAGCGCGTAGGTACATTTTACTCAATTCTCTAATC |
| Δ59-N20-3 | GATTGAAATGGCACCGCCTAGTTTTAGAGCTAGAAATAGCAAGTTAAAATAAGG |
| Δ59-N20-4 | TAGGCGGTGCCATTTCAATCCGTAGGTACATTTTACTCAATTCTCTAATC |
| Δ59-HA-1 | AGTGGTTCTATCGAGAGTCGTGTATACAGGTAGCCGATCGC |
| Δ59-HA-2 | ATAATCGGCGTGTATGACATGTCCACTTGACAGC |
| Δ59-HA-3 | CATGTCATACACGCCGATTATGACAGGTTATTC |
| Δ59-HA-4 | CGGCAAATCCAATCGCATAGTGACCTCATCGATCGCAATT |
| Δ59-Test-1 | ACGAAATCAGCACAAACACC |
| Δ59-Test-2 | TGTTCCGAAGGCTTCTTCCAG |
| Δ59-Test-3 | ACCGTCTGCTGTATTTCATCC |
| Δ59-Test-4 | AGCAAACGAAATGGAATGTGAT |
| Δ60-N20-1 | CACACAGTTCACTTCAACTGGTTTTAGAGCTAGAAATAGCAAGTTAAAATAAGG |

| | |
|---|---|
| Δ60-N20-2 | CAGTTGAAGTGAACTGTGTGCGTAGGTACATTTTACTCAATTCTCTAATC |
| Δ60-N20-3 | AATTGTTCCTGTTTTGCCCGGTTTTAGAGCTAGAAATAGCAAGTTAAAATAAGG |
| Δ60-N20-4 | CGGGCAAAACAGGAACAATTCGTAGGTACATTTTACTCAATTCTCTAATC |
| Δ60-HA-1 | AGTGGTTCTATCGAGAGTCGTTATCAATGGATCCGGCTGC |
| Δ60-HA-2 | ATGTTCCTGTATAGACGAAATGCTAGAGGCGC |
| Δ60-HA-3 | GCATTTCGTCTATACAGGAACATCAAGGTGATACAGG |
| Δ60-HA-4 | CGGCAAATCCAATCGCATAGTAAAACGATGATTGCCGCTT |
| Δ60-Test-1 | TCGCGTTGAATTGACATCAAC |
| Δ60-Test-2 | TGGCCTTATTTTGCTGATGCT |
| Δ60-Test-3 | AGAAACGCCATGGATAATCGC |
| Δ60-Test-4 | TGATGATCATCTTTCAATGGTTGG |
| Δ61-N20-1 | ATTGTCATTCCGATCATGGCGTTTTAGAGCTAGAAATAGCAAGTTAAAATAAGG |
| Δ61-N20-2 | GCCATGATCGGAATGACAATCGTAGGTACATTTTACTCAATTCTCTAATC |
| Δ61-N20-3 | CACATCATTGACACAGCTGAGTTTTAGAGCTAGAAATAGCAAGTTAAAATAAGG |
| Δ61-N20-4 | TCAGCTGTGTCAATGATGTGCGTAGGTACATTTTACTCAATTCTCTAATC |
| Δ61-HA-1 | TTCTCCCCCATTACATCACTACGCCAATAATGCTTCAATCG |
| Δ61-HA-2 | TTCGGCTCCATTTATATTCATGGCTTTCGCAAACC |
| Δ61-HA-3 | TGAATATAAATGGAGCCGAAAATCACCATAG |
| Δ61-HA-4 | GAGGTGACTGAAGTATACCGAATGTAGACAAGGTTGAAGAGGACG |
| Δ61-Test-1 | ATAAAGAAAGCTCCAGTGGCAA |
| Δ61-Test-2 | TACTACATGTCGATGGCGGC |

| | |
|---|---|
| Δ61-Test-3 | TTCTTGGAATACGATGATGACCG |
| Δ61-Test-4 | ATGCCTTGCTCAGGAGTAAAC |
| Δ62-N20-1 | TGATTTAGTATCTCATGCGCGTTTTAGAGCTAGAAATA GCAAGTTAAAATAAGG |
| Δ62-N20-2 | GCGCATGAGATACTAAATCACGTAGGTACATTTTACTC AATTCTCTAATC |
| Δ62-N20-3 | GATTTGCAGGAGTCGTGGTCGTTTTAGAGCTAGAAATA GCAAGTTAAAATAAGG |
| Δ62-N20-4 | GACCACGACTCCTGCAAATCCGTAGGTACATTTTACTC AATTCTCTAATC |
| Δ62-HA-1 | AGTGGTTCTATCGAGAGTCGTGTGAAGACGGTATCGAG CAC |
| Δ62-HA-2 | CAAAGGATGATCAATTATCAACATGCCCGGT |
| Δ62-HA-3 | GTTGATAATTGATCATCCTTTGCTATCTCATCTACCAT |
| Δ62-HA-4 | CGGCAAATCCAATCGCATAGGTGGAACTGACGAAGGA TGAA |
| Δ62-Test-1 | AGAGTTTCTTCTGGATCATCGG |
| Δ62-Test-2 | ACGATGGCTATGGTGAACTC |
| Δ62-Test-3 | ACTTACTCTGTCGAGTTCAGATACGCTCATATTCCCGGT TTACTGTGTACATTCCTCTCTTACCTATAATGG |
| Δ62-Test-4 | TGTTATATTATCAGAAAGGAGGTGATAAAATGAAACAA ACCCATACCAGGATGTAGATGTAGAAATACAAGGTTAC AT |
| Δ63-N20-1 | GACAAATTGGTATCTGACATGTTTTAGAGCTAGAAATA GCAAGTTAAAATAAGG |
| Δ63-N20-2 | ATGTCAGATACCAATTTGTCCGTAGGTACATTTTACTCA ATTCTCTAATC |
| Δ63-N20-3 | GTTACATTTGAGCCGACTAAGTTTTAGAGCTAGAAATA GCAAGTTAAAATAAGG |
| Δ63-N20-4 | TTAGTCGGCTCAAATGTAACCGTAGGTACATTTTACTC AATTCTCTAATC |

| | |
|---|---|
| Δ63-HA-1 | TTCTCCCCCATTACATCACTAGATACAGATACGATTTGTTCTGCG |
| Δ63-HA-2 | AGTTCGAGGTAGATTCATCGAACCATTCAGGAG |
| Δ63-HA-3 | CGATGAATCTACCTCGAACTTATACGCCTATACA |
| Δ63-HA-4 | GAGGTGACTGAAGTATACCGAATCACAAACAATGACAGGCTGG |
| Δ63-Test-1 | ATGATTACACCTGCAGCAAGAC |
| Δ63-Test-2 | ATGGCGAAGCGAATGATTTG |
| Δ63-Test-3 | AGTATTATGAACAGCCTGAACG |
| Δ63-Test-4 | TTGATATAGAGGAACAGGACGAGG |
| Δ64-N20-1 | ATTCATCATCAGGGTGAGGCGTTTTAGAGCTAGAAATAGCAAGTTAAAATAAGG |
| Δ64-N20-2 | GCCTCACCCTGATGATGAATCGTAGGTACATTTTACTCAATTCTCTAATC |
| Δ64-N20-3 | ATGCTGTCACCCTACTCTTTGTTTTAGAGCTAGAAATAGCAAGTTAAAATAAGG |
| Δ64-N20-4 | CAAAGAGTAGGGTGACAGCATCGTAGGTACATTTTACTCAATTCTCTAATC |
| Δ64-HA-1 | AGTGGTTCTATCGAGAGTCGATTATTCGAGTATCTTTCCCAGCAT |
| Δ64-HA-2 | TCGGATGATTTCGAAGCTGATGTCGTACTTGA |
| Δ64-HA-3 | TCAGCTTCGAAATCATCCGAACGGTTCCC |
| Δ64-HA-4 | CGGCAAATCCAATCGCATAGATGGATGTCATGAAGGTTATCAAG |
| Δ64-Test-1 | TGCATTCGGCTATTGGATTGG |
| Δ64-Test-2 | TAAAGCACAACGCGTAACAT |
| Δ64-Test-3 | TTGGCGGATAATCAACAAACC |
| Δ64-Test-4 | AGACAGTGGACATTGTTCGTTAT |
| Δ65-N20-1 | TGATATTAGGCTCTTGCAGTGTTTTAGAGCTAGAAATAGCAAGTTAAAATAAGG |
| Δ65-N20-2 | ACTGCAAGAGCCTAATATCACGTAGGTACATTTTACTCAATTCTCTAATC |

| | |
|---|---|
| Δ65-N20-3 | ACTATGTCCACCACATCTTGGTTTTAGAGCTAGAAATAGCAAGTTAAAATAAGG |
| Δ65-N20-4 | CAAGATGTGGTGGACATAGTCGTAGGTACATTTTACTCAATTCTCTAATC |
| Δ65-HA-1 | AGTGGTTCTATCGAGAGTCGTGATATACAAAGATCCCGAACAAGC |
| Δ65-HA-2 | TTTTGGTTCTTCACTTCAAGCTTTTCTAGAATCATTG |
| Δ65-HA-3 | CTTGAAGTGAAGAACCAAAAGGAGCCAATTG |
| Δ65-HA-4 | CGGCAAATCCAATCGCATAGTTCGATCGCTAGATCATTTGATTTT |
| Δ65-Test-1 | ACTGGTAAAATTAGAAGAACGTCAG |
| Δ65-Test-2 | AGAAACACTAATGCTGGGATGG |
| Δ65-Test-3 | TGCTCTTCACTACCGTTAACC |
| Δ65-Test-4 | AGAGAAGTCTGTGGAGTAGGA |
| Δ21-1N20-1 | ATCTTGCAGGAGCTCTGTCACGTAGGTACATTTTACTCAATTCTCTAATC |
| Δ21-1N20-2 | TGACAGAGCTCCTGCAAGATGTTTTAGAGCTAGAAATAGCAAGTTAAAATAAGG |
| Δ21-1HA-1 | GGTCGACGGCCAACGAGTGGAGTCACAGTAAAATCCG |
| Δ21-1HA-2 | GGATTGATGTAGCTGCTGAAGTAA |
| Δ21-1HA-3 | TTACTTCAGCAGCTACATCAATCCGTCTAAAGCTCTCCATTCCACA |
| Δ21-1HA-4 | ATTTCTTAATCTAGAAAGGCCTTATATGTCGAACGTTCTATCAACC |
| Δ21-2N20-1 | GTTGCGAACTAGTGCTCTTACGTAGGTACATTTTACTCAATTCTCTAATC |
| Δ21-2N20-2 | TAAGAGCACTAGTTCGCAACGTTTTAGAGCTAGAAATAGCAAGTTAAAATAAGG |
| Δ21-2HA-1 | GGTCGACGGCCAACGACTTCAGCAGCTACATCAATCC |
| Δ21-2HA-2 | ACAAGCGCTATTTTACCTTCA |
| Δ21-2HA-3 | TGAAGGTAAAATAGCGCTTGTAATTTCACTGCGCAGAGATTAC |

| | |
|---|---|
| Δ21-2HA-4 | ATTTCTTAATCTAGAAAGGCCTTATCATGGGATGACTCTCAAGGAA |
| Δ21-3N20-1 | CAAGCATTCTCTCGGCTCTACGTAGGTACATTTTACTCAATTCTCTAATC |
| Δ21-3N20-2 | TAGAGCCGAGAGAATGCTTGGTTTTAGAGCTAGAAATAGCAAGTTAAAATAAGG |
| Δ21-3HA-1 | GGTCGACGGCCAACGTAATTTCACTGCGCAGAGATT |
| Δ21-3HA-2 | GAAATCATTACGGTATGTCCTGC |
| Δ21-3HA-3 | GCAGGACATACCGTAATGATTTCCAGCAACTCATGGAGCTTAATC |
| Δ21-3HA-4 | ATTTCTTAATCTAGAAAGGCCTTATCGGAGAAGTAATGATCCATCTT |
| Δ21-4N20-1 | TAATGTATCGCCGGATTCCACGTAGGTACATTTTACTCAATTCTCTAATC |
| Δ21-4N20-2 | TGGAATCCGGCGATACATTAGTTTTAGAGCTAGAAATAGCAAGTTAAAATAAGG |
| Δ21-4HA-1 | GGTCGACGGCCAACGAACTGTGAAAGCCAACTTACAC |
| Δ21-4HA-2 | CCCTTCATTCATATTACTCTCCTTTAGA |
| Δ21-4HA-3 | TCTAAAGGAGAGTAATATGAATGAAGGGGATAATCTTTGAGGTTTCGGAAGC |
| Δ21-4HA-4 | ATTTCTTAATCTAGAAAGGCCTTATTCGTCCAGCCAAAAGGATTT |
| Δ21-5N20-1 | TCCTACACTTGCAGTAGGCCCGTAGGTACATTTTACTCAATTCTCTAATC |
| Δ21-5N20-2 | GGCCTACTGCAAGTGTAGGAGTTTTAGAGCTAGAAATAGCAAGTTAAAATAAGG |
| Δ21-5HA-1 | GGTCGACGGCCAACGTACTCCCGGTGAAACATTATCA |
| Δ21-5HA-2 | CGTTTGATCAACGTGTTGAAT |
| Δ21-5HA-3 | ATTCAACACGTTGATCAAACGAAGGCATAGCGGGACAATTAT |
| Δ21-5HA-4 | ATTTCTTAATCTAGAAAGGCCTTATTGAAATGCGATAACCCTGCTA |

| | |
|---|---|
| Δ21-6N20-1 | GACTGACGCATCTTCTTGTCCGTAGGTACATTTTACTCAATTCTCTAATC |
| Δ21-6N20-2 | GACAAGAAGATGCGTCAGTCGTTTTAGAGCTAGAAATAGCAAGTTAAAATAAGG |
| Δ21-6HA-1 | GGTCGACGGCCAACGAGCTAATCCCGATGTGATTAAAG |
| Δ21-6HA-2 | TTACTAACAACAAGGTAAAGGGG |
| Δ21-6HA-3 | CCCCTTTACCTTGTTGTTAGTAATCTCTAAATGGACATACTGAAAGGAG |
| Δ21-6HA-4 | ATTTCTTAATCTAGAAAGGCCTTATCACCTTTACAAACATACGTTCGTTA |
| Δ21-7N20-1 | GGTTCGTACAGGTTGGTCAACGTAGGTACATTTTACTCAATTCTCTAATC |
| Δ21-7N20-2 | TTGACCAACCTGTACGAACCGTTTTAGAGCTAGAAATAGCAAGTTAAAATAAGG |
| Δ21-7HA-1 | GGTCGACGGCCAACGAATGTTCTACACGAGGCATTACA |
| Δ21-7HA-2 | ATGTCTTCACTTCATATCACTCCT |
| Δ21-7HA-3 | AGGAGTGATATGAAGTGAAGACATAAGCGAATTTCATATGTAAGGTGT |
| Δ21-7HA-4 | ATTTCTTAATCTAGAAAGGCCTTATCTGGACTCTCAGTTCGTTTATG |
| Δ21-8N20-1 | TATCGCTTTACTTCTCAATGCGTAGGTACATTTTACTCAATTCTCTAATC |
| Δ21-8N20-2 | CATTGAGAAGTAAAGCGATAGTTTTAGAGCTAGAAATAGCAAGTTAAAATAAGG |
| Δ21-8HA-1 | GGTCGACGGCCAACGGATGGCAGCTCGTCTGTTT |
| Δ21-8HA-2 | CTGCAAGACGGACAATTTAACA |
| Δ21-8HA-3 | TGTTAAATTGTCCGTCTTGCAGCCCTTATGAATTGTTCTGTCGC |
| Δ21-8HA-4 | ATTTCTTAATCTAGAAAGGCCTTATCAGCCAGAGCAGTGAATTTC |
| Δ21-9N20-1 | GTGCGCAGCTGTTCAATCATCGTAGGTACATTTTACTCAATTCTCTAATC |

| | |
|---|---|
| Δ21-9N20-2 | ATGATTGAACAGCTGCGCACGTTTTAGAGCTAGAAATAGCAAGTTAAAATAAGG |
| Δ21-9HA-1 | GGTCGACGGCCAACGCAACATCATTGACCAACCTGT |
| Δ21-9HA-2 | TTCGGGACGCAACAAAAAT |
| Δ21-9HA-3 | ATTTTTGTTGCGTCCCGAAAGGTTTCTCATACGTGAGAAGC |
| Δ21-9HA-4 | ATTTCTTAATCTAGAAAGGCCTTATTCAGTGTGTGCTGATAGCTTAA |
| Δ21-10N20-1 | GTTCAGTACAACGCTTGCTACGTAGGTACATTTTACTCAATTCTCTAATC |
| Δ21-10N20-2 | TAGCAAGCGTTGTACTGAACGTTTTAGAGCTAGAAATAGCAAGTTAAAATAAGG |
| Δ21-10HA-1 | GGTCGACGGCCAACGATGAATTGTTCTGTCGCTATAAAGA |
| Δ21-10HA-2 | TTCGTTGTCATACAGTTTCCTCCT |
| Δ21-10HA-3 | GGAGGAAACTGTATGACAACGAAACAGCGTCAGATAATTGGATTC |
| Δ21-10HA-4 | ATTTCTTAATCTAGAAAGGCCTTATAGGAATTAAGCCGATGACAATC |
| Δ21-11N20-1 | CAAACTGGAACATGACCCAGCGTAGGTACATTTTACTCAATTCTCTAATC |
| Δ21-11N20-2 | CTGGGTCATGTTCCAGTTTGGTTTTAGAGCTAGAAATAGCAAGTTAAAATAAGG |
| Δ21-11HA-1 | GGTCGACGGCCAACGAATCGATTGAAGCTGCCGAA |
| Δ21-11HA-2 | AATGGCTGTCATTCTGAAAGC |
| Δ21-11HA-3 | GCTTTCAGAATGACAGCCATTTTTCTTACGGGAGGGACCA |
| Δ21-11HA-4 | ATTTCTTAATCTAGAAAGGCCTTATAAAATGATGTTGGGATCGTGAT |
| Δ21-1-test-1 | GCCGAATACCGTATGATGAAC |
| Δ21-1-test-2 | ACTTTGTCCGGGAATAGTGAT |
| Δ21-2-test-1 | ATTAAGGCGTACAATTTGTACGT |

| | |
|---|---|
| Δ21-2-test-2 | CATTACGGTATGTCCTGCTTC |
| Δ21-3-test-1 | CGGGTGTTGAATTGTTCGTAG |
| Δ21-3-test-2 | TCCCAAATCCTGATCCCAATG |
| Δ21-4-test-1 | GTGGCTAATGCTTACCCTAATTT |
| Δ21-4-test-2 | GGCTTTCATCATAGGCCATAAC |
| Δ21-5-test-1 | ATAGAGCCGAGAGAATGCTT |
| Δ21-5-test-2 | TCATGCTTCACCATGCGTAC |
| Δ21-6-test-1 | ACGTTGATCAAACGAACGGA |
| Δ21-6-test-2 | CCGCAATGTCTTCACTTCATAT |
| Δ21-7-test-1 | AACAGTGAGGGCATTCCATAG |
| Δ21-7-test-2 | ATCTTGTTTAGATAACACGGCC |
| Δ21-8-test-1 | CACTCGCCGCTTTCTAATGA |
| Δ21-8-test-2 | GTGAAGCTTGTGTATGCTCAATT |
| Δ21-9-test-1 | GAGTGAGACTTACCATGATATCGC |
| Δ21-9-test-2 | ATTCTCCTTTGGCCTCTGC |
| Δ21-10-test-1 | TCACAAGCTTCCTCTCCTTTAA |
| Δ21-10-test-2 | AACAAGGAAATCAGCACAGC |
| Δ21-11-test-1 | CTGTAGCAAGCGTTGTACTG |
| Δ21-11-test-2 | TTCTTCCGTCCATTCCTTCA |
| Δ21-N20-1 | TGATGGAAATATCGGCTGTAGTTTTAGAGCTAGAAATA GCAAGTTAAAATAAGGC |
| Δ21-N20-2 | TACAGCCGATATTTCCATCACGTAGGTACATTTTACTCA ATTCTCTAATCACGG |
| Δ21-N20-3 | ACGTGAGCGCGTTCTGCCAAGTTTTAGAGCTAGAAATA GCAAGTTAAAATAAGGC |
| Δ21-N20-4 | TTGGCAGAACGCGCTCACGTCGTAGGTACATTTTACTC AATTCTCTAATCACGG |
| Δ21-HA-1 | TTCTCCCCCATTACATCACTTAGAAGGTGTAACATTGCT AGGC |
| Δ21-HA-2 | ATGCTTTGCAGATAGGCTATTATGCGGAGCAAGA |
| Δ21-HA-3 | AATAGCCTATCTGCAAAGCATTGAAGGTATC |

| | |
|---|---|
| Δ21-HA-4 | GAGGTGACTGAAGTATACCGAATGATACAAGTGCTGCA ATAAGCT |
| Δ21-Test-1 | AGCTGAATCTCGTCAGAGTG |
| Δ21-Test-2 | TCCTTCCTTCGCTTTAGCAAATC |
| Δ21-Test-3 | AAGTCCATTCAATATGTTCTGACC |
| Δ21-Test-4 | TAGTTAGTCGTGCGCTTCCAT |

**Table S6. The construction price of Keio collection, CRISPRi, and CHASING.**

| Price | Keio collection | CRISPRi | CHASING |
|---|---|---|---|
| Primer | 52,000 $ | 6,000 $ | 2,000 $ |
| Gibson seamless cloning mix | 20,000 $ | 0 $ | 700 $ |
| High-fidelity DNA polymerase | 10,000 $ | 100 $ | 300 $ |
| Taq DNA polymerase | 1,000 $ | 0 $ | 50 $ |
| Competent cell | 700 $ | 50 $ | 150 $ |
| Sequencing | 20,000 $ | 0 $ | 300 $ |
| Total | 103,700 $ | 6,150 $ | 3,500 $ |

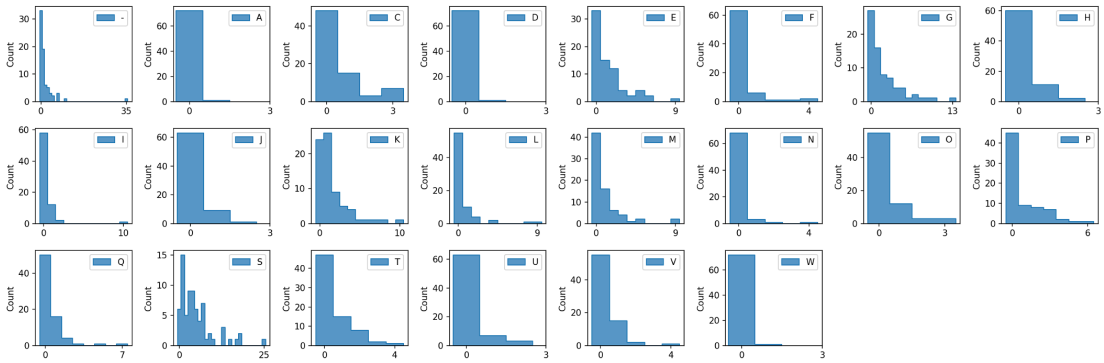


**Figure S1. The number of genes in different COG categories based on the deleted genes analysis of all chromosome segments.**

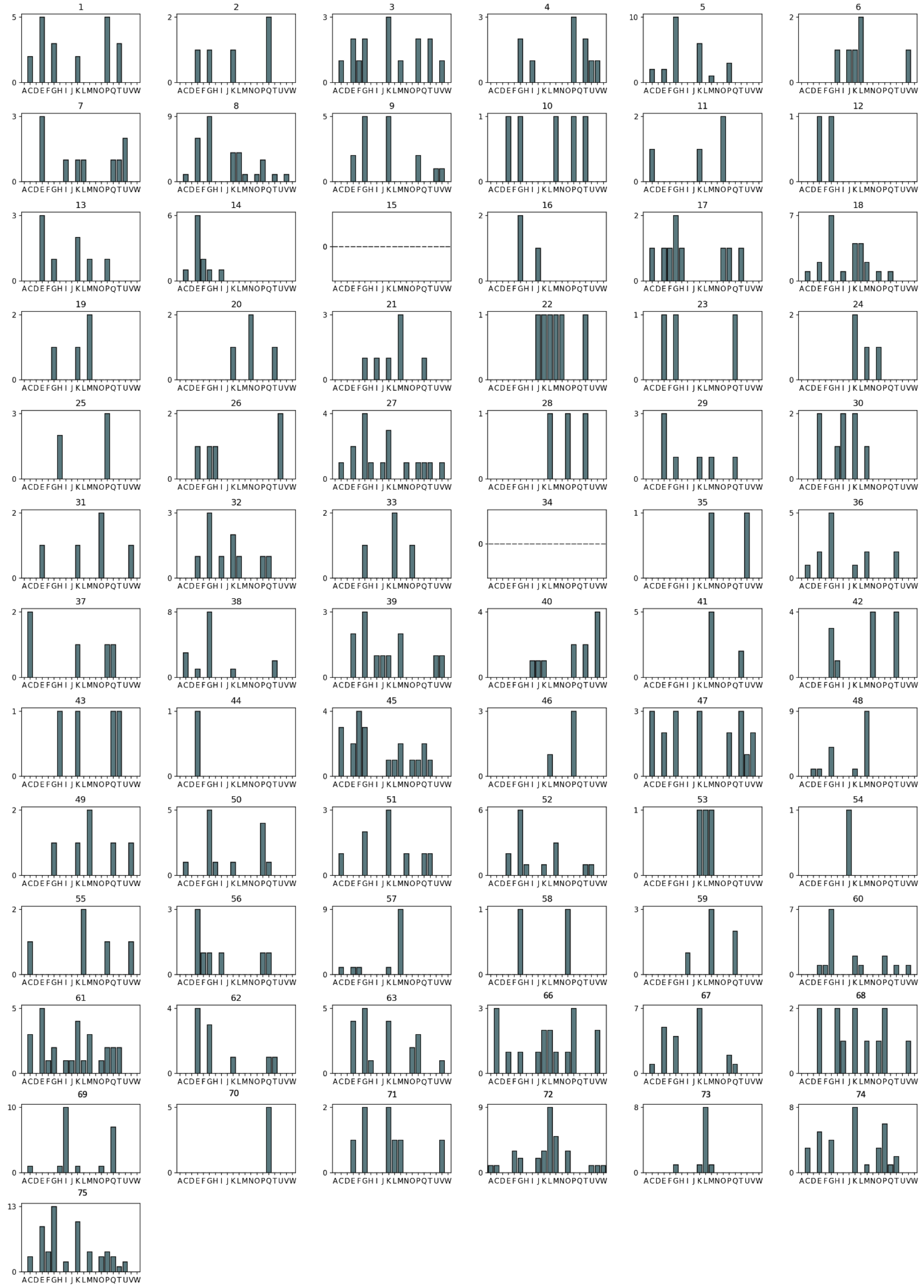

**Figure S2. Gene number analysis of each COG category in each chromosome segment.** The segments 15 and 34 with genes failed to group to any COG category were also included in this figure.

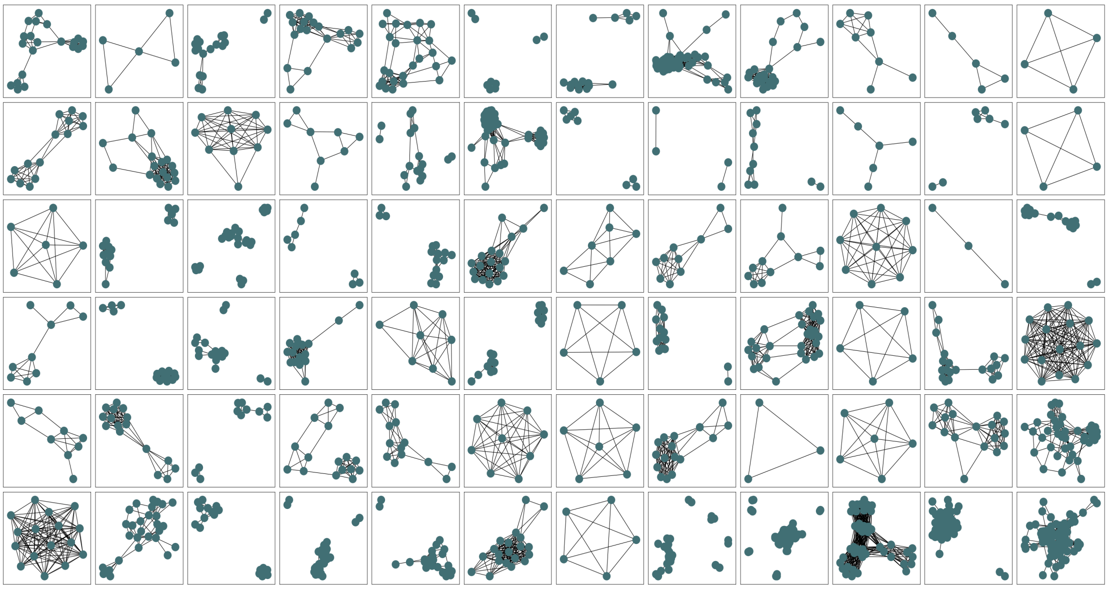

**Figure S3. The interconnection between the proteins in each chromosome segment.** The blue point represents the protein and the line between the blue points indicates the connection of proteins. The segments with no interconnection was excluded in this figure identified for the segment 54.

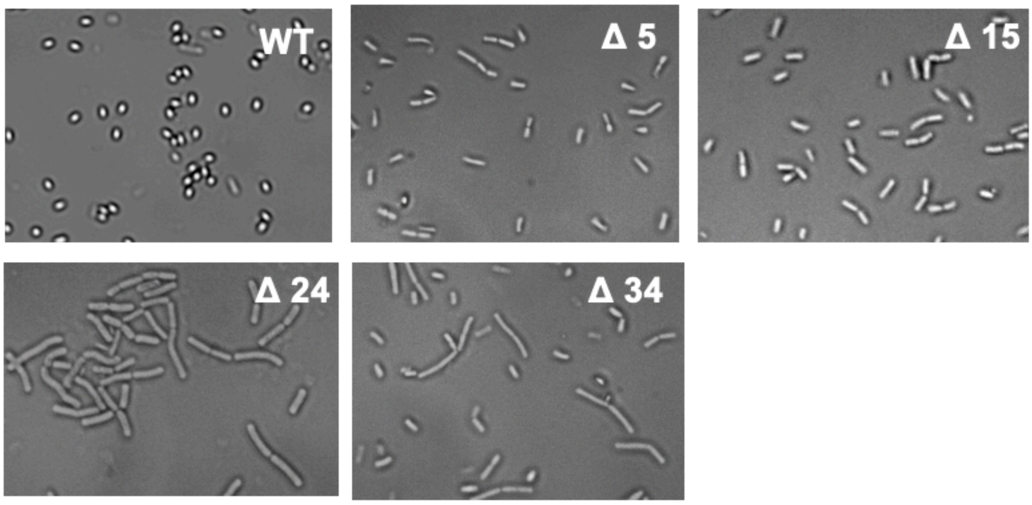


**Figure S4. Sporulation patterns of the wild-type strain and the deletion strains Δ5, Δ15, Δ24, and Δ34.** 48 h-cultivation sample was collected and the morphological pattern was visualized using microscopy.

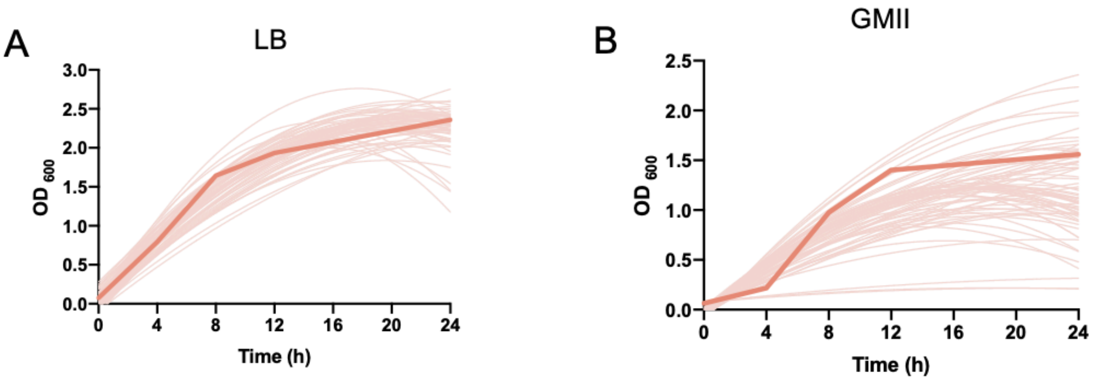


**Figure S5. Cell growth of each deletion strain in LB and GMII medium.** (a) The cell growth of the deletion strains in LB medium. (b) The cell growth of the deletion strains in GMII medium. The pink curve and the bold curve represent the deletion strains and the wild-type strain, respectively.

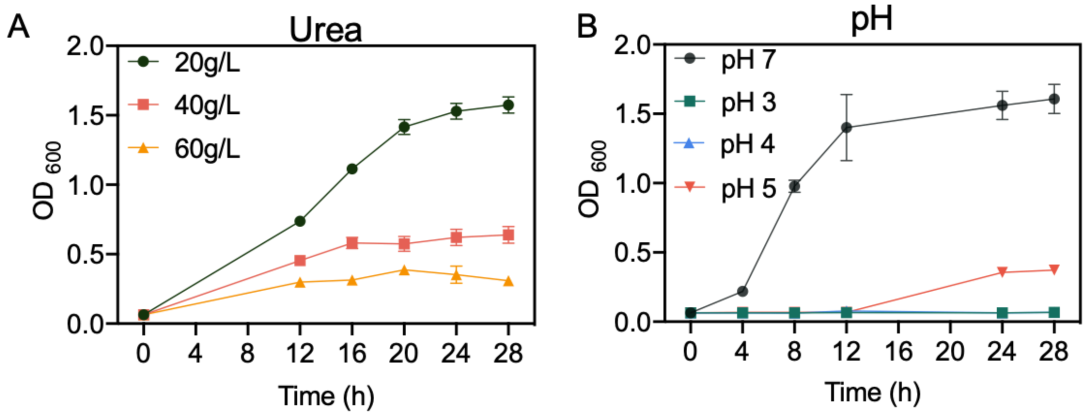


**Figure S6. The cell growth of the wild-type strains with urea shock and pH shock.** (a) The cell growth in the presence of 20, 40 and 60 g/L urea. (b) The cell growth under pH 3, pH 4, pH 5, and pH 7 conditions.

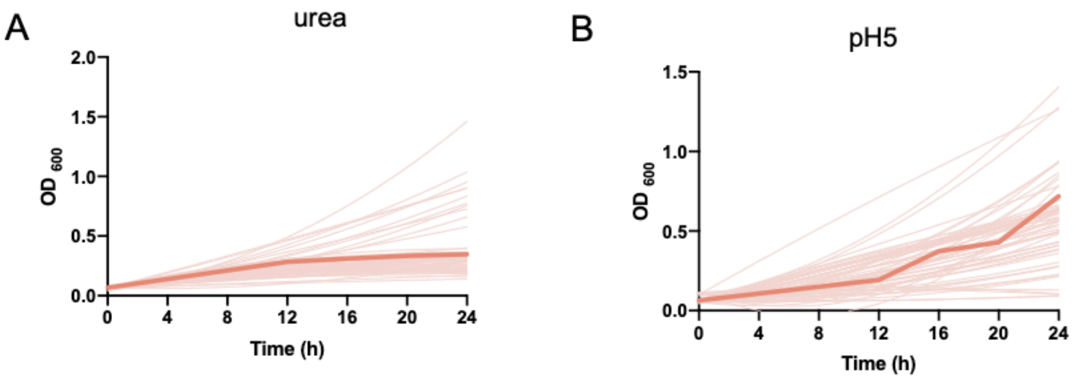


**Figure S7. The cell growth of the deletion strains under high urea and pH5 condition.** (a) The cell growth of the deletion strains under 60 g/L urea stress. (b) The cell growth of the deletion strains at pH 5 condition. The pink curve and the bold curve represent the deletion strains and the wild-type strain, respectively.

CHASING

Enter group number

21

Show large fragment

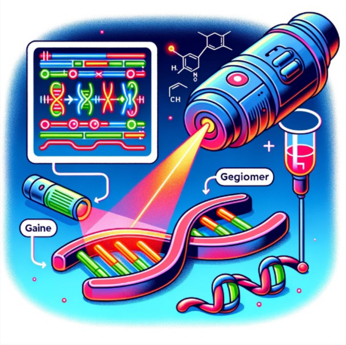


### Chromosome segment scanning for gain- or loss-of-function screening (CHASING) and its application in metabolic engineering

#### Strain No. 21 large fragment knockout

Deletion from 1442346 to 1450587, length: 8.241 kb

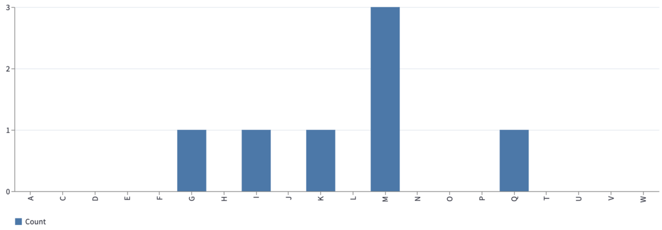


| gene | locus_tag | COG | COG cluster | COG pathway | NCBI COG | UniProt | BsuCyc | STRING |
|---|---|---|---|---|---|---|---|---|
| ykvN | BSU_13760 | K | COG1733 | none | Link to COG | Link to UniProt | Link to BsuCyc | Link to STRING |
| ykvO | BSU_13770 | IQ | COG1028 | Fatty acid biosynthesis | Link to COG | Link to UniProt | Link to BsuCyc | Link to STRING |
| ykvP | BSU_13780 | M | COG1388,COG4641 | none,none | Link to COG | Link to UniProt | Link to BsuCyc | Link to STRING |
| ykzQ | BSU_13789 | M | COG1388 | none | Link to COG | Link to UniProt | Link to BsuCyc | Link to STRING |
| ykvQ | BSU_13790 | G | COG3858 | none | Link to COG | Link to UniProt | Link to BsuCyc | Link to STRING |
| ykzR | BSU_13799 | S | COG3858 | none | Link to COG | Link to UniProt | Link to BsuCyc | Link to STRING |
| ykvR | BSU_13800 | S | | | Link to COG | Link to UniProt | Link to BsuCyc | Link to STRING |
| ykvS | BSU_13810 | S | COG4873 | none | Link to COG | Link to UniProt | Link to BsuCyc | Link to STRING |
| ykzS | BSU_13819 | - | | | Link to COG | Link to UniProt | Link to BsuCyc | Link to STRING |
| ykvT | BSU_13820 | M | COG3773 | none | Link to COG | Link to UniProt | Link to BsuCyc | Link to STRING |
| ykvU | BSU_13830 | S | COG2244 | none | Link to COG | Link to UniProt | Link to BsuCyc | Link to STRING |

**Figure S8. Analysis the fragment 21 COG functions by CHASING web server.**

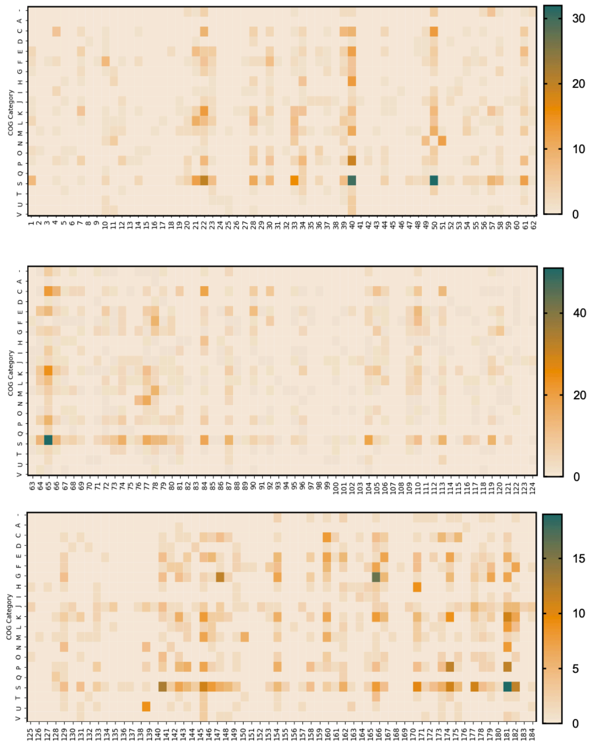


**Figure S9. The COG distribution patterns of the 184 proposed chromosome segments in *E. coli* MG1655.**

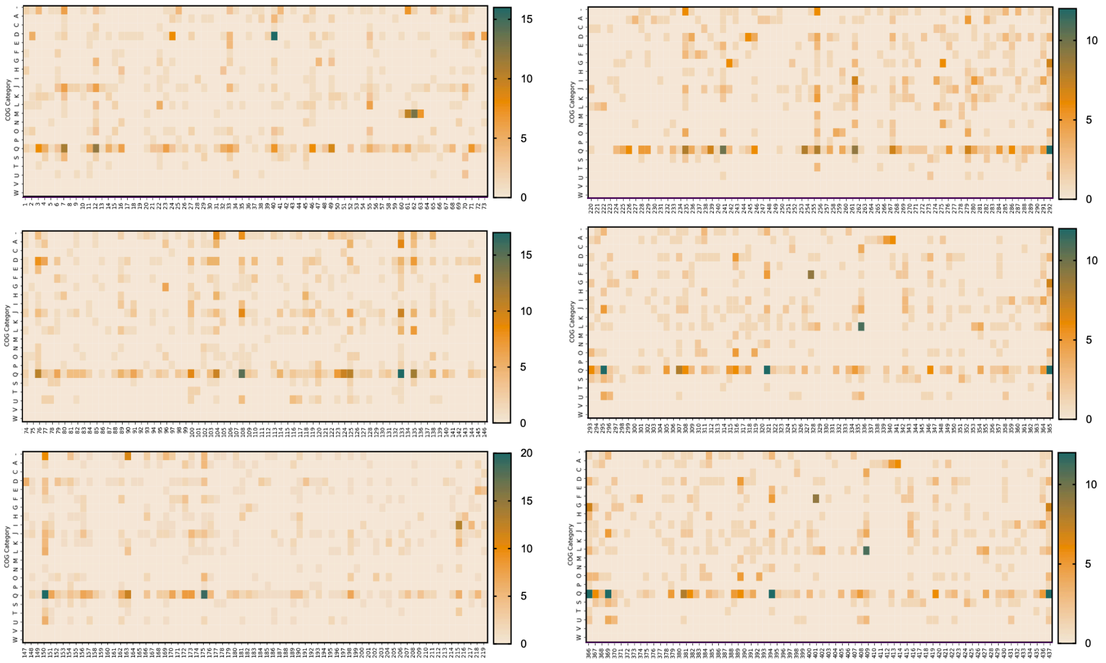


**Figure S10. The COG distribution patterns of the 437 proposed chromosome segments in *B. thuringiensis* BMB171.**

**Supplementary Note1:**

Using two-fold change as the cutoff, the differentially expressed transcripts were identified. Under pH5 condition, 276 downregulated genes and 176 upregulated genes were identified for the strain Δ5 while 121 downregulated genes and 202 upregulated genes were identified for the strain Δ21 (Figs. 5(b) and (c)). Under urea-rich condition, the differentially expressed transcripts included 516 downregulated genes and 238 upregulated genes in strain Δ5, and 253 downregulated genes and 88 upregulated genes in strain Δ21, respectively. To exclude the effect caused by the deletion of unrelated non-essential genes, we performed the cluster analysis to identify the differentially expressed transcripts that are specific for pH5 and urea-rich condition. For strain Δ5, the specific differentially-expressed genes were narrowed down to 561 and 259 for urea-rich and pH5 condition, respectively (Fig. 5(d)). For strain Δ21, the specific differentially-expressed genes were narrowed to 287 and 269 genes under urea-rich and pH5 condition, respectively. Then, the major pathways with the largest number of differentially-expressed genes were identified for the tolerance of acid shock, including the two-component system, quorum sensing protein, phosphotransferase system (PTS) in upregulation and flagellar assembly metabolism in downregulation (Fig. 5(e)). Similarly, the major pathways involved in the tolerance of urea shock were also identified, including the ABC transporters, inositol phosphate metabolism in upregulation, and bacterial chemotaxis in downregulation.
